# Multi-omics Analysis of Dsup Expressing Human Cells Reveals Open Chromatin Architectural Dynamics Underyling Radioprotection

**DOI:** 10.1101/2020.11.10.373571

**Authors:** Craig Westover, Deena Najjar, Cem Meydan, Kirill Grigorev, Mike T. Veling, Roger L Chang, Christopher Chin, Daniel Butler, Evan E. Afshin, Pamela A Silver, Christopher E. Mason

## Abstract

Spaceflight has been documented to produce detrimental effects to physiology and genomic stability, partly a result of Galactic Cosmic Radiation (GCR). In recent years, extensive research into extremotolerant organisms has begun to reveal how they survive harsh conditions, such as ionizing radiation. One such organism is the tardigrade (*Ramazzottius varieornatus*) which can survive up to 5kGy of ionizing radiation and the vacuum of space. In addition to their extensive network of DNA damage response mechanisms, the tardigrade also possesses a unique damage suppressor protein (Dsup) that co-localizes with chromatin in both tardigrade and transduced human cells to protect against DNA damage from reactive oxygen species induced by ionizing radiation. While Dsup has been shown to confer human cells with increased radiotolerance; much of the mechanism of how it does this in the context of human cells remains unknown. Until now there is no knowledge yet of how introduction of Dsup into human cells can perturb molecular networks and if there are any systemic risks associated with foreign gene introduction. Here, we created a stable HEK293 cell line expressing Dsup, validated its radioprotective phenotype, and performed multi-omic analyses across different time points and doses of radiation to delineate molecular mechanism of the radioprotection and assess molecular network pertubations. Dsup expressing human cells showed an enrichment for pathways seen in cells overexpressing HMGN1, a chromosomal architectural protein that has a highly similar nucleosome binding motif. As HMGN1 binding to nucleosomes promotes a less transcriptionally repressed chromatin state, we further explored the hypothesis that Dsup could behave similarly via ATAC-seq analysis and discovered overall selective differential opening and closing of the chromatin landscape. Cut&Run analysis further revealed global increases in histone post translational modifications indicative of open chromatin and global decreases in repressive marks, with Dsup binding preferentially towards promoter regions marked by H3K27ac and H3K4me3. We further validated some of the enriched pathways via in-vitro assays and revealed novel phenotypes that Dsup confers to human cells such as reduction in apoptosis, increased cell proliferation, and increased cell adhesion properties. Our analysis provides evidence that the Dsup protein in the context of HEK293 cells may behave as a chromatin architectural protein and that in addition to its nucleosome shielding effect, may confer radio-resistance via chromatin modulation. These results provide future insight into mitigating some of the major challenges involved with long term spaceflight as well as understanding some of the molecular architectural underpinnings that lead to radioresistant cancer phenotypes back home.

## Introduction

One of the major risks during long term spaceflight is exposure to Galactic Cosmic Rays (GCRs). GCRs are made up of mostly high-energy protons, and to a lesser extent, alpha particles, electrons, and highly damaging HZE nuclei ^1, 2^. As of now there is little data on the associated risks of exposure to space radiation for a three year Mars mission^8^, but it has been estimated that a return trip to Mars could expose astronauts to 600-1000mSv, which is near the NASA astronaut career limit of 800-1200mSv^1, 2^. Based on these estimates, the predicted attributable risk for GCR exposure would suggest a high likelihood of returning astronauts facing a higher risk of leukemia, stomach, colon, lung, bladder, ovarian, and esophageal cancers^2^.

Selection of radioresistant individuals for space travel has been proposed as one option to circumvent the effects of these massive doses of radiation exposure. Candidates could be selected based on their rate of DNA damage accumulation and repair, as measured by comprehensive multi-omic analyses, or prioritizing those with lower rate of mutations measured with clonal hematopoiesis^1^.

Protective mechanisms identified from studies on bacteria and multicellular extremophiles have also been proposed as a candidate defense against the extreme environment of space via genetic engineering of radioprotective components into human cells^1^. Studies on various bacteria have shown increases in mutation rates as well as increases in virulence, antibiotic resistance, metabolic activity, shorter lag phase time, and a number of beneficial adaptations in response to short orbital flights^8, 10, 11^. While there is not a selective pressure to withstand high doses of radiation on this planet, it is speculated that there is an evolutionary link between increased radiotolerance and the ability to enter an anhydrobiotic state as many radiotolerant eukaryotic species do have this ability to dehydrate for extended periods of time and so there is a selective pressure to withstand DNA damage from endogenous reactive oxygen species generated during times of desiccation^3, 5, 12, 13^. As such, desiccated tardigrades of the species *R. coronifer* and *M. tardigradum* have been shown to survive the combined effects of the vacuum of space, galactic cosmic radiation on the scale of 9.1Gy, and different spectra of UV radiation at a total dose of 7577kJ/m^2^ at low earth orbit^3, 5, 12, 13^. Anhydrobiosis in tardigrade species *R. varieornatus* has been demonstrated to provide better aid in the prevention of DNA damage accumulation than in the hydrated stage in response to UV-C radiation as measured by UV-induced thymine dimers^16^. However, hydrated tardigrades in another study were shown to be more resistant to heavy ion radiation than in their anhydrobiotic state as demonstrated as the LD_50_ for heavy ions being 4.4kGy to 5.2kGy for hydrated and dehydrated animals respectively^1^. As most of the damage from High-linear energy transfer (LET) radiation is deposited along linear tracks to pass right through cells and induce double stranded breaks (DSBs) as opposed to the indirect effects of ROS induced by Low-LET, there could be separate mechanisms for defending against varying types of radiation. An overlap of functional redundancy between proteins of both systems could be possible as adult tardigrades of various species have been shown to be just as tolerant to High-LET as they are tolerant to Low-LET radiation^1, 3, 5, 13, 16, 17^. One such protein that has been shown to protect against low-LET radiation in human cells is the recently discovered Damage suppressor protein, Dsup^18^.

Out of several novel genes identified from the complete *R. varieornatus* sequence, Dsup became a likely candidate for conferring tardigrades with increased radiotolerance as tandem mass spectrometry revealed Dsup colocalized with nuclear DNA and could potentially protect against radiation induced DNA breaks. By expressing Dsup in human cells, this hypothesis was confirmed as Dsup was shown to suppress about 40% of X-ray induced DNA breaks compared to WT cells^18^. It was then later discovered through a series of biochemical assays using purified protein that Dsup preferentially bound to nucleosome associated DNA through its highly basic c-terminal domain and competed against histone H1 for nucleosome binding. Moreover, this nucleosome binding region was shown to exhibit a highly similar amino acid sequence with the vertebrate HMGN1 protein in its intrinsically disordered RRSARLSA motif^19^. Interestingly, HMGN1 also competes against histone H1 for nucleosome binding sites in a dynamic flux of chromatin remodeling based on cellular context such as development, differentiation, and DNA damage response^24–26^. As a chromosomal architectural transcription factor, HMGN1 is known to affect histone post-translational modifications (PTMs) while activating ATM in response to DNA damage which leads to a more open chromatin state and allows for less sterically hindered access of DNA repair machinery to the site of the lesion. Conversely, binding of histone H1 to nucleosomes leads to a more condensed chromatin state and can inhibit the effects of HMGN1 through increasing histone deacetylase PCAF activity^35, 36, 38^. Interestingly, HMGN protein knock-out in in vitro and in vivo mouse models have shown increased radiosensitivity in response to UV and ionizing radiation and a lack of DNA damage repair leading to G2-M checkpoint arrest^21, 22^.

Understanding the molecular mechanisms underpinning extremophile tolerance could provide us clues on how human survival in space could be improved through genetic engineering. However, it is essential to understand how introducing a foreign gene from one species to another affects the overall system. Here, we developed a lentiviral-transduced HEK293 cell line containing Dsup (HEK293-Dsup) from which we assessed system wide stability via transcriptomic and epigenetic analyses and tested functional assays that measure biomarkers and cellular processes indicative of DNA damage. The current proposed mechanism for which Dsup prevents DNA damage is that is suppresses DNA breaks and acts as a physical protectant against damage inducing agents^18, 19^. Given some of the similarities to HMGN proteins we hypothesized that it is possible that Dsup may also act functionally similar to HMGN1 in terms of enhancing DNA repair by promoting a more open chromatin state in addition to its overall shielding effect^19^, but there is no functional genomics data to yet verify this hypothesis until now. Overall, these data show a range in functionality of the radioprotective effects, with changes in overall gene expression and chromatin landscape, and provide a resource of data on human cells regulatory changes when utilizing the foreign Dsup protein.

## Results

### Stable integration of Dsup into HEK293 cells reduces DNA damage biomarkers

Hashimoto and colleagues created a HEK293 Dsup cell line under a CAG promoter via a proprietary lipofectamine transfection^10^. We attempted our own stable integration via insertion of a human codon optimized CMV promoter Dsup via lentiviral transduction into HEK293 cells.

After serial dilution cloning, we confirmed expression by western blot (Figure 1A) and Dsup insertion site into the right arm of chromosome 5 as assessed by Promethion long read sequencing (Figure 1B).

**Figure 1.**
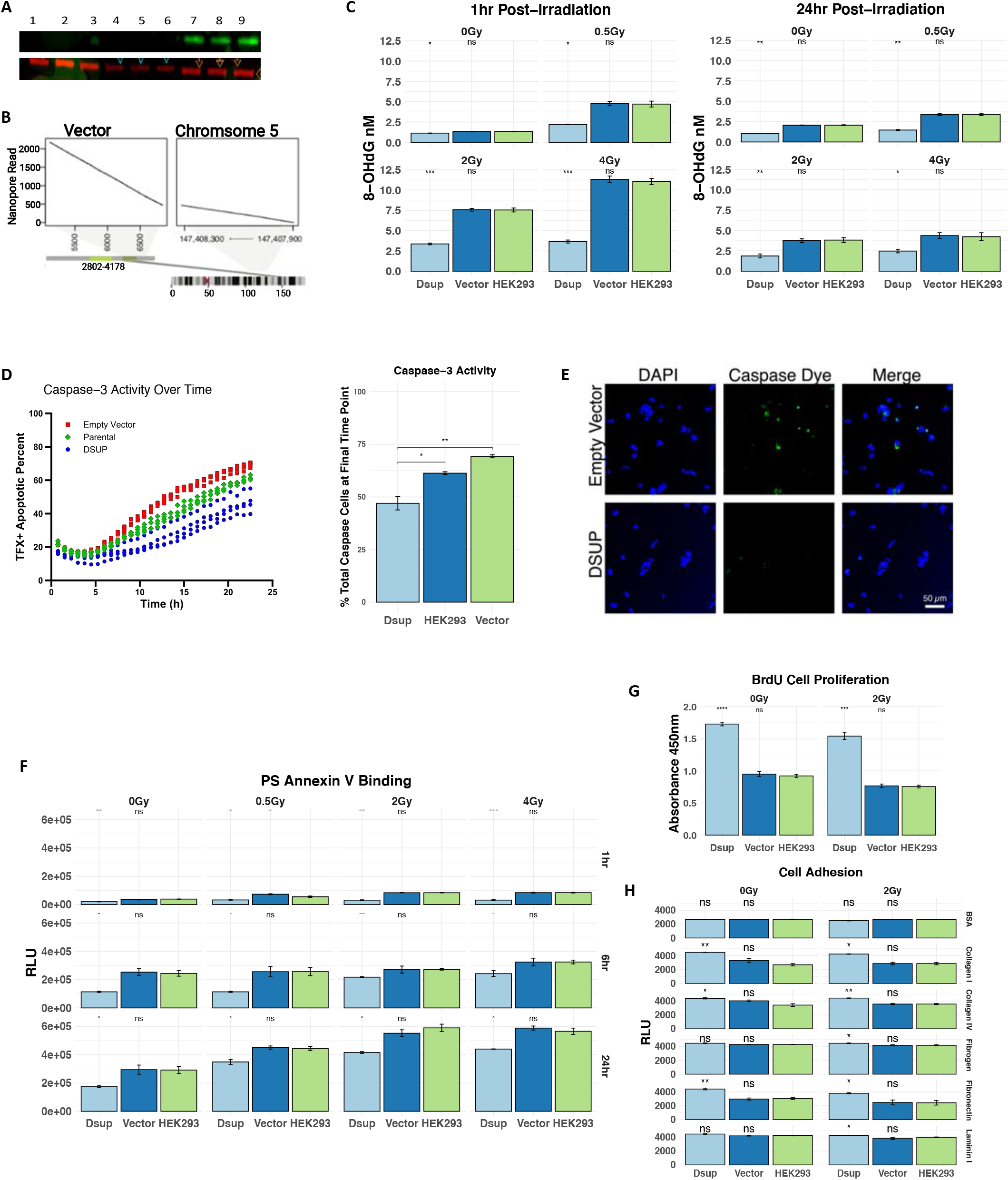
Introduction of Dsup into HEK293 cells results in prevention of oxidative stress and apoptosis while increasing cell proliferation and cell adhesive properties. (A) Confirmation of Dsup introduction into HEK293 cells via western blotting of lysates from HEK293 and HEK293 cells stably transduced with codon optimized pLVX-Puro Dsup and pLVX-Puro Empty Vector. Lanes 1-3 include HEK293, Lanes 4-6 include HEK293 Empty Vector, and lanes 7-9 include HEK293 Dsup. Primary R&D Systems recombinant mouse HA Tag primary antibody (MAB0601) was used to bind to HA-tagged Dsup at a 0.1ug/ml dilution and then visualized with Rabbit anti-Mouse IgG (H+L) Cross-adsorbed Secondary Antibody Alexa Fluor Plus 680 (A32729) at 0.1ug/ml. Bio-Rad Mouse anti Human Vinculin antibody (V284) was used as a control at a 1:500 dilution and visualized with Invitrogen Goat anti-Rabbit IgG (H+L) Cross-Absorbed Secondary Antibody DyLight 800 (SAS-10036) at a 1:5000 dilution. (B) Dsup integration site was assessed via Oxford Nanopore long read sequencing with 2000 reads mapping to the vector at position 5,500 bp. The first 500 bases map to the right arm of chromosome 5 while the remaining 1500 bases map to the vector site downstream of Dsup. Parental HEK293 cell line and HEK293 Dsup were sequenced with a genome coverage of 2.8x and 1.1x respectively. (C) 8-OHdG nM concentration was measured from cell lysates via ELISA assay for both 1hr and 24hrs post-irradiation with significant reductions in HEK293 Dsup as calculated by students t-test with HEK293 WT as the reference group. (D) Caspase-3 activity was monitored over time for all cell types. 13.2μM of CPT was added to each cell line and Hoechst and Caspase staining was used to discriminate live cells from apoptotic cells. Images were collected and FITC intensity was analyzed and plotted over time for all wells. Percentage of apoptotic cells at the end of the experiment was calculated at 22.5 hours. Error bars show 1 standard deviation of the apoptotic rate of the cells. A heteroscedastic two tailed student t-test was used to test the null hypothesis that the empty vector and Dsup containing cells have equal rates of apoptosis (p = 0.005). (E) Representative images of caspase-3 apoptosis assay. Left panels contain DAPI stained nuclei, center contains caspase-3 dye, and the right panel is the merged images. The top rows are HEK293 cells transduced with the empty pLVX-puro vector and bottom row are HEK293 Dsup cells. Scale bar is at 50μm. (F) Apoptosis via Annexin V binding to phosphatidyl serine was assessed over 0Gy, 0.5Gy, 2Gy, and 4Gy of irradiation and plotted across time at 1hr, 6hrs, and 24hrs (Top to bottom panels) for all cell types. Apoptotic activity was measured as relative luminescent output (RLU) (G) Cell proliferation as measured by total BrdU incorporation after a 24hr period and measured via absorbance at 420nm for both 0Gy (left) and after 2Gy irradiation (right) for all cell types. (H) Cell adhesive properties to different surfaces was measured as lysed adherent cell fluorescence as detected by dye. All cell types were added in triplicates to each of the ECM coated wells with BSA serving as a negative control. Unless otherwise noted all significant figures were obtained by a heteroscedastic two tailed student t-test was used to test the null hypothesis that the HEK293 and other cells for each condition are equal. ns=p>0.05, *p<0.05, **p<0.01, ***p<0.001.

As it was clear that Dsup expressing cells had a significant advantage in reducing double stranded DNA breaks and early DNA damage response indicators from ROS,^10^ we then looked at how well Dsup could reduce other molecular markers of oxidative stress such as 8-hydroxy-2-deoxoguanosine (8-OHdg)^52^. 8-OHdg is a large bulky adduct that results from ROS interaction with DNA and is typically repaired quickly through base excision repair^56^. While Dsup has been shown to reduce damage typically repaired by nonhomologous end joining and homologous recombination^20^, we tested if Dsup could also prevent the accumulation of 8-OHdg in cells. We assayed cell lysates via ELISA for presence of 8-OHdg in HEK293 Dsup, HEK293 WT, and HEK293 transduced with an empty pLVX vector. At 1hr post irradiation HEK293 Dsup had a significantly lower amount of 8-OHdg at baseline 0Gy (p<0.05) and at all doses of radiation (0.5Gy p<0.01, 2Gy p<0.001, & 4Gy<0.001). At 24hrs post irradiation most of the 8-OHdg levels were reduced and with significant reductions still in HEK293 Dsup in irradiated cells (0.5Gy p<0.001, 2Gy, p<0.001, & 4Gy p<0.01) (Figure 1C).

Hashimoto and colleagues first noted that Dsup expressing human cells continued to proliferate after a dose of 4Gy of radiation, which was enough to cause WT HEK293 cells to enter a senescent state and pause proliferation. Here they assessed proliferation in terms of increased cell viability and observed that even unirradiated Dsup expressing cells continued to proliferate slightly faster than controls^10^. As regulation of proliferation and apoptosis are tightly linked^46, 49, 57, 60^, we wanted to next examine if Dsup expressing cells are more viable as previously reported^10^ because they also have a reduction in programmed cell death. Here we examined caspase-3 activity directly via caspase staining and microscopy over a 22.5hr time frame by inducing DNA damage with 13.2 μM cisplatin. We observed a 37% reduction in caspase signal intensity for HEK293 Dsup vs empty vector control (p=0.001) and a 31% reduction for HEK293 Dsup vs HEK293 cells (p<0.005). The total percent of caspase-3 positive cells was also quantified at the end of the experiment using a cutoff of 650 intensity units measured against the background and classifying apoptotic vs non apoptotic cells in this way. Using this metric, we observed 32% reduction in percent apoptotic HEK293 Dsup cells compared to empty vector control (p=0.005) and 23% reduction compared to WT control (p<0.05) (Figure 1D). 10x objective representative images taken at the center of the wells show signal intensity for DAPI stained nuclei, FITC showing caspase-3 dye, and merged channels (Figure 1E).

To further validate apoptotic signal reduction in Dsup expressing cells, we also examined one of the earliest events in apoptotic signaling, the binding of Anexin V to phosphatidyl serine (PS)^57^ for different doses of radiation and at three time points. Here PS Annexin V binding was measured as two Annexin V luciferase components binding together when PS exposure resulting from flipping from the inner leaflet to outer membrane brings the Annexin V units into complementary proximity; the output of which was measured as Relative Luminescence Units (RLU). From these measurements we saw a decrease in early apoptosis induction in HEK293 Dsup vs control HEK293 cells at baseline 0Gy and 1hr (p<0.01). The effects of PS Annexin V binding became more pronounced over increasing doses of radiation especially at the 1hr timepoint with HEK293 Dsup cells having a 62% reduction at 4Gy (p<0.001). This trend continued into the 6hr and 24hr time points with apoptotic signal increasing over time for HEK293 Dsup having significant reductions across all the time points and at the higher doses of radiation. (Figure 1F).

We next looked more in depth at cell proliferation and hypothesized that Dsup could lead to an increased rate in cell proliferative ability particularly through regulation of the cell cycle at S phase as HMGN1, a higher eukaryotic protein with a similar nucleosome binding domain^19^, is known to directly enhance the rate of recruitment of PCNA during this phase to sites of replication to increase transcriptional output and sites of DNA damage to increase repair during this DNA damage checkpoint^96, 97^. Here we measured cell proliferation via the incorporation of 5’-bromo-2’-deoxyuridine (BrDU)^98^ into replicating cells at S phase and measured relative absorbance at 420nm for all cell types at either 0Gys or 2Gys 24hrs post irradiation. In the non-irradiated cells, HEK293 Dsup had nearly a 1.9x increase in proliferation compared to control HEK293 cells and at 2Gy as well (p<0.001 & p<0.001) (Figure 1G).

Cell proliferation and evasion of apoptosis has also been linked to regulation of Extra Cellular Matrix (ECM) dynamics. It has been demonstrated that increased collagen formation as well as LOX over expression can lead to ECM stiffness that then in turn facilitates upregulation of integrin signaling necessary for cell survival and proliferation^99^. Chromosomal architectural proteins like HMGN1 can directly modulate chromatin architecture, and so an increased adhesive phenotype could be likely as the nucleus plays an additional role as a mechanosensor in which its envelope, filaments, and chromatin are linked to the extracellular environment by integrin and cytoskeleton complexes^100^. We tested if Dsup expression could lead to a similar phenotype via a fluorometric cell adhesion assay in which unirradiated and cells irradiated with 2Gy were plated on wells containing different ECM components before. HEK293 DSUP had a 26% increase in cells adhered to Collagen I in comparison to control cells at 0Gy (p<0.01) and a 32% increase once irradiated compared to irradiated control (p<0.05). Similar increases were observed for cell adhesion to Collagen IV with HEK293 Dsup having an 8% increase in the number of cells adhering to the wells (p<0.05) and increasing to 19% compared to control after 2Gy of irradiation (p<0.01). Other notable shifts in cell adhesion properties were seen in HEK293 Dsup adherence to Fibronectin at 0Gy where a 32% increase was seen (p<0.01) and a 34% increase at 2Gy (p<0.05) (Figure 1H).

### Dsup expression promotes upregulation of pathways associated with a more transcriptionally permissive state

It has been well documented that chromosomal architectural transcription factors play a role in modulating transcriptional states through their direct interactions with the chromatin landscape^21^. In particular, the family of High Mobility Group proteins (HMGNs) bind either directly to nucleosomes or to the tails of core histones via their highly acidic c-terminal domain in order to promote a de-compacted and therefore active chromatin state where pol II and III transcription is facilitated^25^. We hypothesized that the similar c-terminus of Dsup could play a role in enhancing transcription in addition to its DNA damage shielding effect in human cells. We set out to probe this question through a transcriptomic analysis of Dsup expressing cells in comparison to WT HEK293 cells over a 24hr time period in order to gain a longitudinal understanding of how Dsup can affect transcriptional states over this period. Additionally, we collected samples at increasing doses of radiation to acquire insight into what cellular stress networks may be affected in the presence of Dsup and in relation to no treatment baseline expression.

HEK293 Dsup and HEK293 WT cells were then subjected to 0Gy, 0.5Gy, 1Gy, 2Gy, and 4Gy of X-ray irradiation and collected at 1hr, 6hrs, and 24hrs post irradiation for RNAseq. When comparing overall gene expression dynamics at each time point, there was an overall upregulation of significantly expressed genes in HEK293 Dsup vs WT with most downregulation occurring at 6hrs (Figure 2A, Supplemental Figure S1A). Unsupervised variance stabilizing transformation (VST) clustering revealed a strong Euclidean distance gene correlation driven by the presence of the Dsup genotype itself, followed by time and then radiation dose. When separated by time point, the greatest magnitude of separation by genotype was at 6hrs as revealed by PCA and t-SNE plots (Supplemental Figures S1B & S1C).

**Figure 2.**
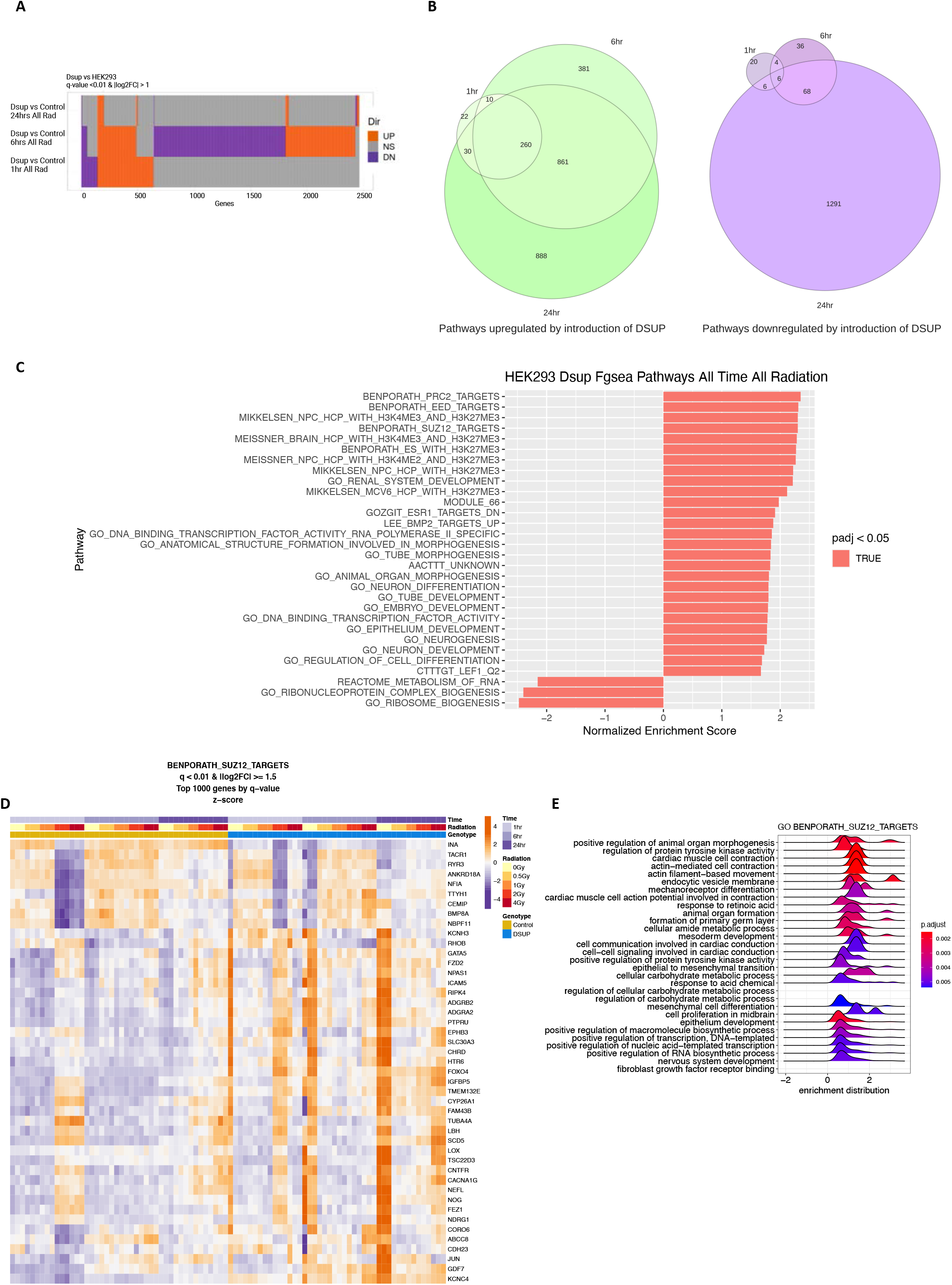
RNA-seq analysis reveals a unique transcriptionally permissive state. **(A)** Altuna plot shows comparison of differentially expressed genes at different time points of Dsup expressing HEK293 cells vs WT Control for all doses of radiation, q-value <0.01 and absolute value of Log_2_FC ≥ 1. X-axis represents total gene counts with approximate counts visualized for each up/ns/down groups by their relative sizes. **(B)** Venn diagrams show the intersections and exclusive counts of upregulated and downregulated pathways as assessed by Fgsea analysis based on differentially expressed signatures obtained by DEseq2 and Limma Voom, q-value < 0.01 and absolute value of NES > 1.2. Gene lists were ranked by absolute value of Log_2_FC ≥ 1.5 for all doses of radiation across each time point. **(C)** Enrichment plots based on running score and pre-ranked Fgsea results were used to visualize top upregulated pathways after normalization for time and radiation dose with NES, p-values, and p-adjusted values plotted alongside. **(D)** Heatmap representing enriched pathways via Fgsea analysis for top differentially expressed genes belonging to BENPORATH_SUZ12_TARGETS, q-value<0.01 and absolute value of Log_2_FC ≥ 1.5. Gradient represents degree of normalized enrichment scores with orange representing upregulated pathways and purple representing downregulated pathways. Hierarchical clustering of genes is split by time, radiation dose, and genotype. **(E)** Ridgeplot shows further GO analysis of pre-ranked leading edge genes belonging to BENPORATH_SUZ12_TARGETS with p-adjusted value cutoff=0.05 and Bonferroni p-adjusted method. Gradient shows p-adjusted values from red to blue going from smallest to largest with height of peaks representing density of enrichment distribution scores.

To gain insight regarding what kind of genes expression changes occur post irradiation we looked at some of the top pathway changes occurring at 1hr 0Gy and 1hr 4Gy. Separating by time and dose for 1hr and 0Gys for HEK293 Dsup vs WT revealed upregulated pathways at baseline involved in transcription with some top upregulated genes coding for Zinc Finger proteins (Supplemental Figures S2A, S2B, & S2C). When taking a closer at the top differentially expressed Dsup vs Control at 1hr and 4Gys, enrichment analysis revealed an upregulation of GO pathways related to DNA replication and nucleosome organization and a downregulation of pathways related to kidney development and extracellular matrix organization (Supplemental Figure S3A & S3B). Genes belonging to each pathway revealed an upregulation of several genes coding for histone components and a downregulation of collagen related genes (Supplemental Figure S3C).

We next explored gene trajectory dynamics affected by time and radiation dose (Supplemental Figure S4A). Log_2_FC expression scores were assessed for both control and Dsup cells with comparison to zero mean, in these cases 1hr and 0Gys were considered the baseline from which Log_2_FC was normalized to. Cluster 01 and Cluster 11 exhibited some of the greatest shifts in Log_2_FC for genes in the time centric plot and so were further investigated for assessment of gene dynamics belonging to these groups. Genes were filtered initially from DEG analysis and only those with an absolute value Log_2_FC cutoff of 1 were normalized and the top differentially expressed genes for each group were plotted via heatmap. Enricher analysis produced associated pathways for cluster 1 with top hits including chromatin organization and DNA damage response pathways while cluster 11 were pathways involved in extracellular matrix organization and collagen formation (Supplemental Figures S4B & S4C). Control WT HEK293 cells had a more negative Log_2_FC at 4Gy and 1hr while HEK293 Dsup at this time point and dose of radiation had a more positive Log_2_FC so this cluster was further explored across additional time points. At 1hr all genes belonging to DNA repair pathways were downregulated for WT HEK293 cells while DNA repair genes such as INO80D (normalized Control Log_2_FC = -1.0, Dsup Log_2_FC = 0.08) and chromatin organization genes such as BRPF3 (normalized Control Log_2_FC = -0.70, Dsup Log_2_FC = 0.30) and KAT7 (normalized Control Log_2_FC = -0.54, Dsup Log_2_FC = 0.30) were upregulated (Supplemental Figures S5A & S5B). At 6hrs post-irradiation WT HEK293 cells had a delayed response for upregulation of DNA damage repair genes while Dsup expressing cells had a lower Log_2_FC for many of these genes and a downregulation of RIF1(normalized Control Log_2_FC = 0.24, Dsup Log_2_FC = -0.10) (Supplemental Figure S6A & S6B). At 24hrs and 4Gys of radiation HEK293 Dsup had a greater upregulation of chromatin organization genes such as BRWD1 (normalized Control Log_2_FC = 0.21, Dsup Log_2_FC = 0.55) and OGT (normalized Control Log_2_FC = 0.05, Dsup Log_2_FC = 0.41)(Supplemental Figure S7A & S7B).

After observing an overall unique and more transcriptionally active state in Dsup expressing cells we assessed the overall enriched pathways across conditions to understand how introduction of Dsup may perturb human transcriptomic networks. Fast gene set enrichment analysis revealed a high degree of overlap of upregulated pathways across each time point for all radiation doses and in this case most of the downregulated pathways occurred at 24hrs (Figure 2B). Plotting the gene list ranks by normalized enrichment score and p-adjusted value for each subset of genes generated a list of top upregulated and downregulated pathways. At 1hr there was enrichment for gene targets of Polycomb Repressor complex such as those belonging to ZNF740_TARGET_GENES (NES=1.30, & p-adjusted value = 1.47e-03), PCGF1_TARGET_GENES (NES=1.44, & p-adjusted value = 1.13e-02), and DOUGLAS_BMI1_TARGETS_UP (NES=1.39, & p-adjusted value = 1.71e-02) (Supplemental Figure S8A). Of overall downregulated pathways at 1hr DACOSTA_UV_RESPONSE_VIA_ERCC3_COMMON_DN (NES=-1.71, & p-adjusted value = 1.71e-05) was downregulated compared to HEK293 Control cells (Supplemental Figure S8B). At 6hrs and all doses of radiation Dsup expressing cells had an upregulation of GO_REGULATION_OF_GROWTH (NES=1.30, & p-adjusted value = 9.79e-04) (Supplemental Figure S8C) with downregulations of GO_HOMOLOGOUS_RECOMBINATION (NES=-2.22, & p-adjusted value = 9.08e-04) and DACOSTA_UV_RESPONSE_VIA_ERCC3_XPCS_DN (NES=-1.65, & p-adjusted value = 3.63e-03) (Supplemental Figure S8D). At 24hrs and all doses of radiation Dsup expressing cells had an upregulation of hypoxia related pathways such as MENSE_HYPOXIA_UP (NES=1.97, & p-adjusted value = 2.77e-07) and collagen synthesis pathways such as REACTOME_COLLAGEN_FORMATION (NES=1.94, & p-adjusted value = 1.40e-05) (Supplemental Figure S9A). Downregulated pathways in HEK293 Dsup include GO_RRNA_METABOLIC_PROCESS (NES=-3.41, & p-adjusted value = 1.12e-08) and GO_SPLICEOSOMAL_COMPLEX (NES=-2,42, & p-adjusted value = 1.12e-08) (Supplemental Figure S9B). When comparing the overall effect of Dsup integration into the human genome across time points with all radiation doses included there was a strong enrichment for upregulated pathways related to a more open chromatin state and specific transcriptional activation such as MARTENS_TRETINOIN_RESPONSE_UP, MEISSNER_BRAIN_HCP_WITH_HEK4ME3_AND_H3K27ME3, and BENPORATH_PRC2_TARGETS. Other pathways were related to extracellular matrix components and development (Supplemental Figure S9C). Overall, we saw a downregulation of pathways related to mRNA splicing and processing (Supplemental figure S9D). When normalizing for time and radiation, all top upregulated enriched pathways were gene targets belonging to bivalently marked PRC2 targets followed by differentiation pathways and increased transcriptional activity (Figure 2C). A heatmap was generated to take a closer look at genes belonging to BENPORATH_SUZ12_TARGETS and track how these genes are regulated over time and radiation dose (Figure 2D). We then analyzed these same genes belonging to BENPORATH_SUZ12_TARGETS with KEGG enrichment terms and discovered an enrichment for developmental pathways, transcription, and proliferation (Figure 2E). We also observed genes in MARTENS_TRETINOIN_RESPONSE_UP that were enriched in Hippo, Wnt, and PI3K-AKT signaling pathways as well as an upregulation in focal adhesion (Supplemental Figures S10A, S10B, & S10C). Genes associated with MIKKELSEN_NCP_HCP_WITH_H3K27ME3 were enriched in upregulated KEGG pathways for developmental pathways and metabolism of reactive oxygen species while cell death pathways were downregulated (Supplemental Figures S10D, S10E, & S10F). KEGG Pathways associated with MEISSNER_NCP_HCP_WITH_H3K4ME2_AND_H3K27ME3 were enriched for developmental and Wnt signaling pathways (Supplemental Figures S11A, S11B, & S11C).

We further evaluated some of these findings with KEGG analysis and GO enrichment. Here, we ranked our Limma-voom results for Dsup genotype correcting for radiation and time effects by adjusted p-value multiplied by Log_2_FC (p-adjusted < 0.01 and |Log_2_FC| 1.5) and plotted KEGG pathways and GO enrichments by multiple categories. Overall, there was a downregulation of certain KEGG terms of interest such as Oxidative phosphorylation and pathways related to RNA transport and splicing. Upregulated KEGG pathways included those involved in proliferation, differentiation, and cell survival such as ErbB signaling, breast cancer pathways, and acute myeloid leukemia pathways, and focal adhesion (Supplemental Figure S12A) Of the biological processes plotted, most of the top categories belonged to broad terms such as biological regulation, metabolic process, and response to stimuli, but also more specific terms such as cell proliferation and growth were also enriched for (Supplemental Figure S12B). There were more enriched gene ontology sets that contributed to membrane components followed by nucleus and then several components that contribute to extracellular space (Supplemental Figure S12C). As for molecular function categories, the most enriched terms were protein binding, ion binding, and then nucleic acid binding (Supplemental Figure S12D). When looking at just baseline HEK293 Dsup expression, Shiny GO Network analysis via hypergeometric distribution testing of DEGs belonging to 0Gy and 1hr revealed several enrichments of developmental related pathways (Supplemental Figure S12E).

### Dsup expression in HEK293 cells induces dynamic opening and closing of ATAC-seq peaks

Given that Dsup expression led to a transcriptionally active and similar transcriptional profile to HMGN1 over expression^27^, we hypothesized that introduction of Dsup into human cells could induce a more open and permissive chromatin state in specific locations that would normally be condensed. As time and radiation did not have as strong an effect as the Dsup genotype itself on HEK293 cells it is likely that Dsup could have a direct effect on shaping chromatin state dynamics. This effect could overshadow typical responses to time and radiation through promoting a baseline radio-tolerant state or due to Dsup’s shielding effect of nucleosomes. To test this, we generated ATAC-seq data across the three time points for comparison of HEK293 Dsup vs WT for 0Gys baseline and 2Gys of radiation. When performing unsupervised clustering on the samples a clear segregation between genotype was seen mainly for PCA, more than what was observed in the RNAseq dataset. Just like the 6hr time point for the RNA-seq, the ATAC-seq profile also separated strongly by Euclidean distance at this time point (Supplemental Figures S13A & S13B). Differential expression analysis was performed with DESeq2 and edgeR and MA and volcano plots were then used to visualize fold change deviations based on a FDR ≤ 0.05 and revealed nuanced changes in peak deviations from M=0. Slight shifts towards positive LogFC were seen at the 1hr and 24hr time points at baseline 0Gy, with 2Gys of radiation having more upregulated peaks at these time points while 6hrs had a shift towards negative LogFC values at 2Gys (Supplemental Figures S14A & S14B).

After differential peak analysis using Limma-voom and DESeq2, we categorized the peak count distribution into 3 main categories for CpGs, enhancers, and genic regions. While most regions remained stable, particularly in enhancer regions, there was variation in CpG island distribution where the largest peak count could be seen in closing peaks at 0Gys and 6hrs. When looking at genic regions, there was an enrichment for intronic regions in opening peaks while most closing regions had a greater enrichment for coding sequences and 5′ UTRs at 6hrs (Figure 3A). The same data can be viewed for only significant differential closing and opening of peaks via peak count or as the fraction of differential peaks where most opening of peaks can be seen at HEK293 Dsup 0Gy 1hr baseline with closing of peaks occurring mainly in 5′ UTR and coding sequence regions. Upon irradiation at 1hr 5′UTR and CDS regions became slightly more open, while most closing peaks occurred at 6hrs with 5kb upstream regions of TSS becoming more open and 3′UTR peaks becoming stabilized post-irradiation (Supplemental Figure S15A & S15B). To gain a more nuanced insight into Dsup induced chromatin accessibility dynamics, peaks were annotated as either the distribution of gene TSS distance to the nearest peak within 5kb (Figure 3B) or the distribution of peaks to the nearest TSS within 5kb (Supplemental Figure S15C). The former annotation matches one gene per peak while the latter can match one peak to multiple genes. Plotting these distributions revealed similar profiles where across all time points and 0Gy and 2Gys of radiation the density of peaks was mostly stable and were highest closest to the TSS and 2.5kb away. Peaks closest to the nearest TSS were generally stable except at 6hrs for both doses, where most of the density of peaks were closed. Most opening peaks were typically 2.5kb away from the nearest TSS and there was no change in peak stability at 0Gys and 24hrs. Peak opening enrichment was also plotted via heatmap where HEK293 Dsup and HEK293 WT ATACseq signals were plotted 2kb upstream and downstream of the TSS for all genes belonging to GRCh38. Here we observed an enrichment of opening around genes occurring mostly in HEK293 Dsup at 0Gy and 1hr compared to control cells but becoming less enriched following irradiation. At 6hrs both HEK293 Dsup and WT have the same enrichment of signal but HEK293 post irradiation remains more open around these genes than HEK293 Dsup. At 24hrs HEK293 Dsup at 0Gy and 2Gy has greater signal enrichment than WT HEK293 cells (Supplemental Figure S15D). Peak stability was also plotted via Altuna plot and Venn diagrams to further illustrate shared and exclusive peak counts between the cell types at different doses and time points. Most shared upregulated peaks occurred between Dsup 0Gy and 6hrs and Dsup 2Gy and 24hrs, while most shared downregulated peaks occurred between Dsup 2Gy and 1hr and Dsup 2Gy and 24hr, q<0.01 and absolute value Log_2_FC ≥ 0.58 (Supplemental Figures 16A & 16B).

**Figure 3.**
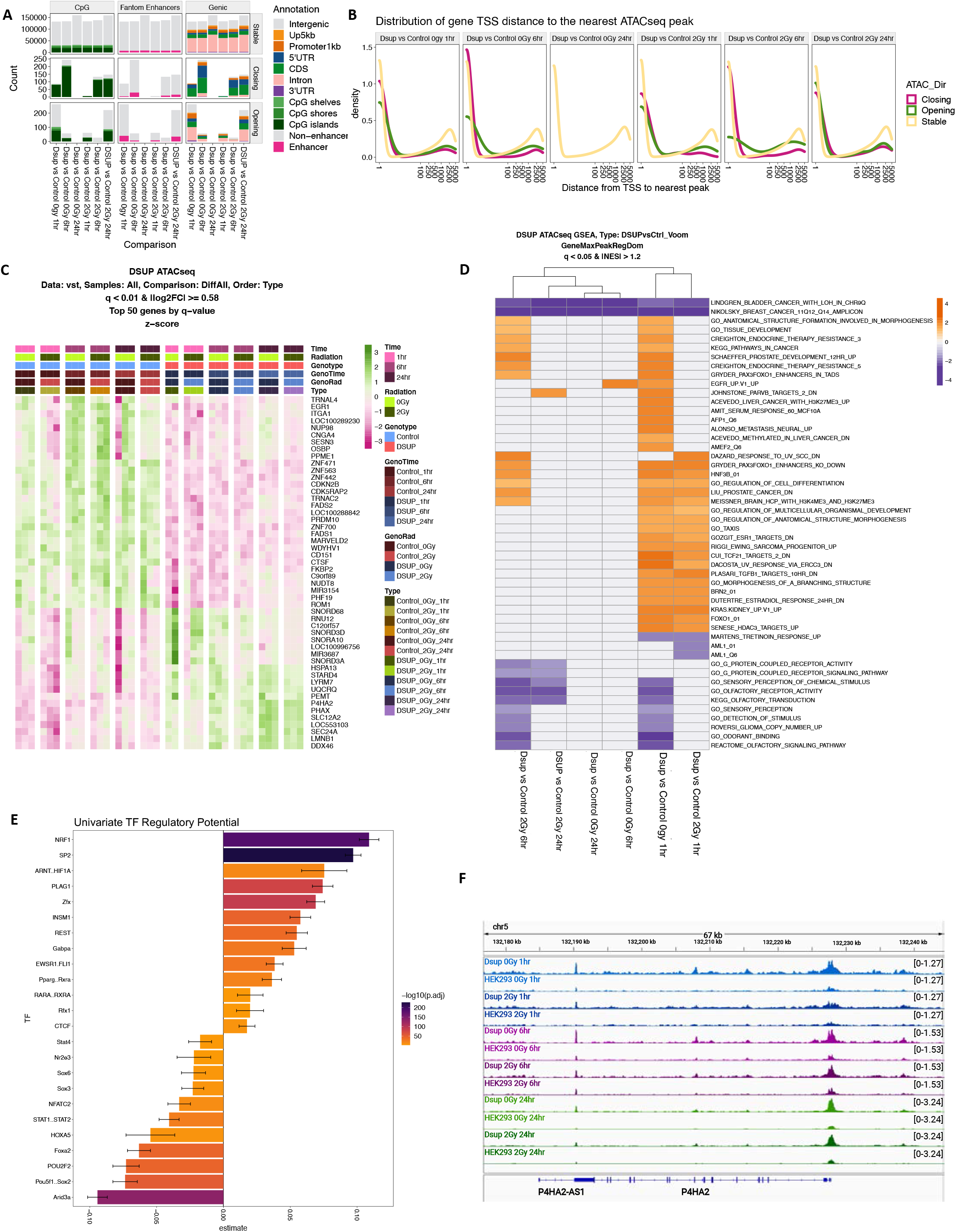
Selective open chromatin accessibility dynamics seen in Dsup expressing HEK293 cells across radiation dose and time. **(A)** Peak count distributions for each dose and time point for HEK293 Dsup vs WT control were annotated via CpG, enhancer, and genic regions using hg38 ENSEMBL BioMart annotations after filtering differential peaks with Log2FC ≥ 0.58 and q-value < 0.01 to obtain stable, opening, and closing annotated peaks. **(B)** Gene transcription start sites were annotated to the nearest differential peak within 5kb for each dose and time point to match one gene per peak and relative densities of opening, closing, or stable peaks were plotted along this distance Log_2_FC ≥ 0.58 and q-value < 0.01. **(C)** Hierarchical clustering for time, genotype, radiation dose and multivariate models including genotype interaction with radiation and time were visualized by Heatmap revealing top 50 differentially annotated peaks to associated nearest gene promoter Log_2_FC ≥ 0.58 and q-value < 0.01. Gradient represents degree of differential expression with green representing upregulated peaks and purple representing downregulated peaks. **(D)** Heatmap representing enriched pathways via Fgsea analysis for differentially enriched peaks annotated to the nearest gene regulatory domain, q-value<0.05 and absolute value of NES > 1.2. Gradient represents degree of normalized enrichment scores with orange representing upregulated pathways and purple representing downregulated pathways.**(E)** Bar plot shows top differentially enriched transcription factor Jaspar Motifs binding sites via univariate modeling, Log_2_FC ≥ 0.56 and q-value cutoff < 0.05 for 200bp peak width. Gradient represents –log10 p-adjusted value with top values ranging from dark purple to light orange. **(F)** IGV tracks shows a representative locus on chromosome 5 for top differential peak P4HA2 across time and radiation dose where Dsup expressing cells have an enrichment for more open chromatin accessibility around this region.

Genes were then annotated to the nearest peak interval and the top 50 associated peaks were clustered by q-value and viewed via heatmap with time, radiation, and genotype taken into account along with other multivariate models such as the intersection of time with genotype and radiation with genotype. Notably, clustering revealed an upregulation of P4HA2, a Polycomb target involved in collagen synthesis^114^. As our data showed a clear bimodal peak distribution and differences in opening and closing dynamics at promoter regions and regions 5kb away from the TSS we further explored these areas on the basis that chromatin modifications involved in increased transcriptional output can occur both at promoters and distal regulatory regions^101^. In order to understand some of the functional consequences of these differential peak dynamics, pathway analysis was performed by running Fgsea on peaks annotated to their nearest regulatory domain. We were able to elucidate the top differential pathways based on the groupings of their associated peaks; moreover, many of these pathways agreed with our RNAseq gene set enrichment analysis. For an overall assessment of the Fgsea results for each dose and time point, HEK293 Dsup vs WT comparisons were visualized by heatmap and an enrichment for an upregulation of peaks associated with genes involved in selective induction of a more open chromatin state such as MEISSNER_BRAIN_HCP_WITH_H3K4ME3_AND_H3K27ME3 was more open at 0Gy and 2Gys at 1hr (Figure 3D). The same data was visualized for each dose and time point using lollipop plots where leading edge size and adjusted p-values could also be visualized. At 1hr and 0Gy, BENPORATH_SUZ12_TARGETS were also found to be in opening peaks while GO_CHROMOSOME_CONDENSATION associated peaks were found to be more closed in Dsup expressing cells. At 2Gy and 1hr, BENPORATH_NANOG_TARGETS and BENPORATH_SOX2_TARGETS were in more accessible chromatin, while peaks associated with GPCR signaling and cell adhesion were in closing peaks (Supplemental Figures S17A & S17B). At baseline and 2Gys of radiation at 6hrs, peaks associated with Nikolsky breast cancer genes were in closed peaks in HEK293 Dsup cells (Supplemental Figures S18A & S18B). Again at 24hrs and 0Gy, annotated peaks associated to Nikolosky breast cancer were downregulated, while at 2Gy annotated peaks associated with biological adhesion were downregulated while differentiation and development associated pathways as well as MEISSNER_BRAIN_HCP_WITH_H3K4ME3_AND_H3K27me3 were found in open accessible regions (Supplemental Figures S19A & S19B).

We next looked at transcription factor binding sites within our differential peak dataset by running a motif univariate model and annotating these TF footprint sites with the JASPAR database, q-value < 0.05 and Log_2_FC ≥ 0.58. In the univariate model each transcription factor is checked individually without assuming if transcription factor A affects transcription factor B given that they may bind to similar regions. We observed NRF1 being the most statistically significant TF motif enriched for followed by SP2 for HEK293 Dsup while Arid3a and Pou5f1 were more enriched in WT HEK293 cells (Figure 3E). Multivariate models were also run where all TFs were checked at the same time and Bayesian probabilities were calculated for TF A given TF B. Here we observed at 0Gy and 2Gy at 1hr statistical significance of CTCF in Dsup expressing cells, while SP2 was more significant in HEK293 WT cells. Interestingly, at 6hrs CTCF motif was more enriched in HEK293 WT cells (Supplemental Figures S18C & S18D) but then became more significantly enriched for again at 24hrs 0Gy (Supplemental Figure S19C). At 2Gy 24hr the top significant TF motifs were Foxq1 and Foxd3 (Supplemental Figure S20C).

### Greater nucleosome occupancy and larger fragment size in HEK293 Dsup

As we’ve demonstrated that Dsup induces changes in chromatin architecture we next looked at how Dsup may affect nucleosome organization as this also plays a role in regulating gene expression. In embryonic stem cells, HMGN1 plays a role in the organization of nucleosomes at active promoters and those containing CpG Islands^109^. Nucleosomes themselves have also been implicated in shielding DNA from radiation induced damage in in-vitro models and when HMGB1, a member of the High Mobility Group proteins that also influences nucleosome occupancy, is knocked out cells accumulate more DSB damage^108^.

We began by comparing nucleosome fragment length distribution in HEK293 Dsup cells vs WT. We found that HEK293 Dsup cells had a greater frequency in larger enriched nucleosome fragment sizes than WT at 0Gy and post-irradiation with sub-nucleosome fragments making up a small portion of the peaks at 0Gy called by Nucleoatac as seen violin plots indicating more protected nucleosome occupied DNA, p<0 and p<.0024 respectively, Wilcoxon signed rank test (Figure 4D). This trend continued into 6hrs and 24hrs with HEK293 Dsup at 2Gy and 24hrs having the largest distribution of larger nucleosome enriched DNA fragments compared to WT (p<0, Wilcoxon signed rank test). While fragment sizes were markedly different between the cell types, spacing between neighboring nucleosomes were not. For the most part inter-dyad distance was typically the same between the cell types with only slight significant differences at 0Gy 1hr (p=0.036, Wilcoxon signed rank test), and both baseline and 2Gy at 6hr, where HEK293 cells had a slightly higher mean compared to HEK293 Dsup (p=0.015 & p=0.025, respectively, Wilcoxon signed rank test). (Figure 4).

**Figure 4.**
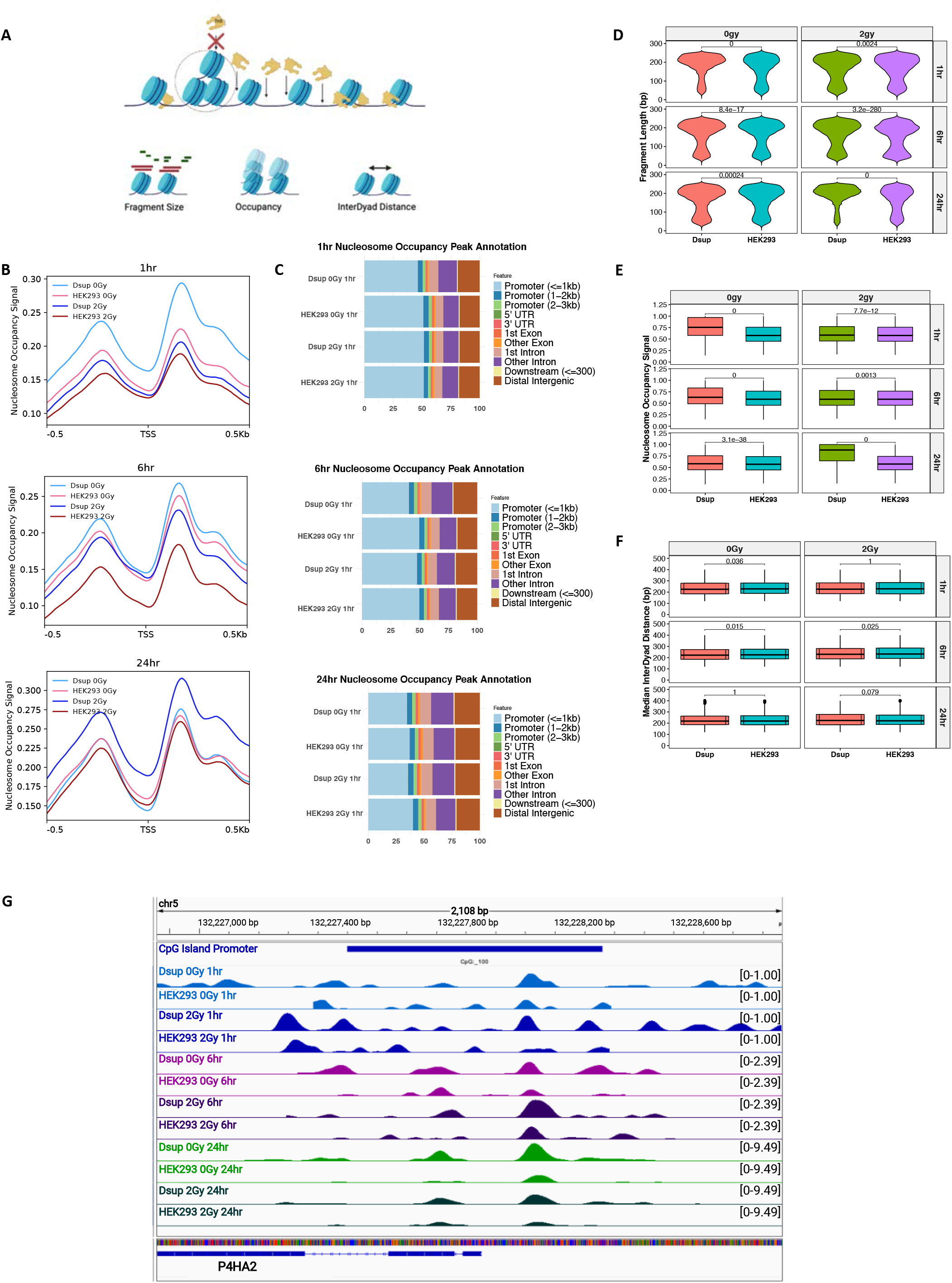
Nucleosome occupancy and fragment length far greater in HEK293 Dsup cells. **(A)** Schematic representation of definitions for fragment size, nucleosome occupancy, and nucleosome inter-dyad distance. Tn5 transposase cuts around nucleosomes in open chromatin regions while blocked from densely packed chromatin. Fragment size refers to the length of DNA wrapped around a nucleosome while sub-nucleosomal fragments <147bp are enriched for in nucleosome-free regions. Occupancy refers to the read pileup producing signal for mono-nucleosomes at dyad centers. Inter-dyad distance is the length between neighboring nucleosome dyad centers. **(B)** Nucleosome occupancy signal as calculated by NucleoATAC after MACS2 broad peak calling and normalization were plotted in metagene plots 500bp upstream and downstream of GRCh38 transcription start sites for Dsup vs WT control at 0Gy and 2Gy of radiation across time at 1hr, 6hrs, and 24hrs. **(C)** Bar plots show percent of annotated feature for nucleosome occupancy peaks for each cell type and dose of radiation across time. Peaks were annotated ± 3000bp of TSS regions for annotated hg38 promoters. **(D)** Violin plots show nucleosome fragment length as computed by NucleoATAC for all broad peaks called from MACS2 peak caller. **(E)** Nucleosomes from occpeak Nucleoatac output was used to generate boxplots for nucleosome occupancy signal for all broad peaks called by MACS2 **(F)** Boxplots for median inter-dyad distance was plotted for all neighboring nucleosomes belonging to all genes from GRCh38 called by NucleoATAC. All Statistical analysis was performed using Wilcoxon signed rank test with significant values plotted over bars. **(G)** IGV tracks shows a representative locus on chromosome 5 for top differential peak P4HA2 across time and radiation dose using combined NucleoATAC occupancy and positioning signal with signal plotted primarily over its associated CpG promoter.

We next analyzed nucleosome occupancy surrounding the TSS of genes belonging to GRCh38. Here nucleosome occupancy is defined as the pileup of reads producing signal for mono-nucleosomes at dyad centers^102^ (Figure 4A). We observed greater occupancy signal in Dsup expressing cells at 1hr and 0Gy as well as when comparing irradiated HEK293 Dsup to WT. This trend also continued into 6hrs, but at 24hrs irradiated HEK293 Dsup had the greater occupancy signal (Figure 4B). Here nucleosome occupancy is defined as the pileup of reads producing signal for mono-nucleosomes at dyad centers^102^ (Figure 4A). When annotating these nucleosome occupancy peaks to annotated features ± 3000bp from the nearest TSS of hg38 promoters, the majority of nucleosome occupancy could be seen as occurring mostly in promoter regions 1kb away from the nearest TSS with a slightly greater percentage in HEK293 cells (Figure 4C). Nucleosome occupancy for all open chromatin regions were then assessed and boxplots were generated to show that post-irradiation occupancy signal was lower in Dsup expressing cells than non-irradiated HEK293 Dsup but higher than irradiated WT (p<7.7e-12).

This trend continued at 6hrs post irradiation (p<0 for 0Gy and p <0.0013 for 2Gy), but at 24hrs HEK293 Dsup at 2Gy had the highest signal followed by Dsup 0Gy (p<3.1e-38 for 0Gy and p<0 for 2Gy, Wilcoxon signed rank test)(Figure 4E).

As there was generally more nucleosome occupancy around promoter regions and our ATAC-seq data showed an enrichment for PRC2 targets regions, we next looked at nucleosome occupancy in CpG island promoters as many of these regions contain targets of Polycomb group repressor proteins^110^. Studies have also shown HMGN1 binding patterns to these regions, particularly during transcriptional activation^109^, and so we tested the hypothesis that there would be greater nucleosome occupancy enrichment at CpG island promoters in HEK293 Dsup. We began by plotting nucleosome occupancy signal over all TSS of CpG Island promoters belonging to hg38 and observed lower signal than the aggregate of identifying occupancy at all TSS belonging to GRCh38. However, similar trends were identified where HEK293 Dsup tended to have greater signal than WT except at 0Gy 24hrs later (Supplemental Figures SA). Further annotation using KEGG and GO analysis of the aggregate of nucleosome occupied CpG Island Promoter peaks revealed an enrichment for several pathways related to cell cycle, axon guidance, and cell adhesion molecule binding (Supplemental Figures S20B & S20C). Nucleosome occupancy signal was also plotted over hg38 active Cage promoters and Fantom enhancers.

Overall, more signal was enriched over Cage promoters than enhancers, but similar trends were found with nucleosome occupancy signal where HEK293 Dsup had more signal at 0Gy and 2Gy for both promoters and enhancers compared to WT HEK293 cells. This trend continued into 0Gy and 2Gy at 6hrs. At 24hrs HEK293 Dsup vs WT signals were roughly the same for promoters and enhancers at 0Gy, but at 2Gy HEK293 Dsup had greater occupancy signal than WT HEK293 (Supplemental Figure S20D).

We then calculated nucleosome inter-dyad distance defined as the distance between neighboring nucleosome dyad centers for all genes belonging to hg38 (Figure 4A). At 0Gy and 1hr HEK293 cells had slightly greater distance between dyads and at 2Gy there was no significant difference (p=0.036 & p=1, Wilcoxon signed rank test). HEK293 cells exhibited slightly larger nucleosome inter-dyad distances at both baseline and irradiation (p=0.015 & p=0.025, Wilcoxon signed rank test). At 24hrs there was no significant difference between the cell types for both 0Gy and 2Gys (p=1 & p=0.079, Wilcoxon signed rank test).

Last we plotted nucleosome signals over the CpG Island promoter region of P4HA2 as this was an enriched peak from our ATAC-seq data. Here we observed increased nucleosome occupancy over this promoter for HEK293 Dsup at all time points and doses of radiation where Dsup 0Gy and 1hr had more nucleosome peaks along a greater length of the gene. Spacing between neighboring nucleosomes was also greater for Dsup expressing cells at 0Gy and 1hr and became spatially closer after irradiation (Figure 4G).

### Cut&Run profiling reveals global increases in transcriptionally active histone post translational modifications and decreases in repressive marks in HEK293 Dsup

While previous reports have shown that short term induction of HMGN1 over expression did not significantly alter chromatin accessibility^27^, our own data suggests that Dsup does modulate chromatin state; albeit, that HEK293 Dsup chromatin state dynamics does appear mostly stable across time with variation in specific annotated genic locations. Global and locus specific variations in histone post translational modifications on the other hand have been shown to be more prevalent in epigenetic variations among HMGN1 over expressing cells vs WT cells; in particular, increases in the active histone PTMs H3K27ac and H3K4me3 and decreases in the repressive histone mark H3K27me3^27, 88^. As these marks are typically associated with bivalently poised promoters such as those involved in differentiation and are either silenced or have low transcriptional activity as a result of PRC2 repression^27, 101, 102^, we next set out to probe whether Dsup induction would promote similar epigenetic changes. In addition, we also tracked changes in these epigenetic marks in response to irradiation as fluctuations in opening and closing of chromatin regions following irradiation and DNA damage have been demonstrated such as H3K4me3 removal and increased H3K27me3 deposition via EZH2 around double stranded breaks^103,104^.

After scaling to 1x normalization of reads per genomic content, profile plots and heatmaps of H3K4me3, H3K27ac, H3K27me3, and EZH2 signals were generated over the transcription start sites and centers of all genes belonging to hg38 genome assembly and revealed global differences in peaks for Dsup vs WT at baseline and 1hr post irradiation. Here we observed a global increase in H3K4me3 for both baseline and 2Gy of radiation (p<0.0001) with a greater drop in H3K4me3 signal post-irradiation in WT cells as expected (Figure 5A). After differential peak analysis and feature annotation of H3K4me3, most of these sites were those that gained and lost H3K4me3 marks compared to WT occurred in promoter regions (Supplemental Figure S21A). This finding is consistent with our increased transcriptional output, upregulation of bivalently marked PRC2 target genes, and slight increase of globally active promoter regions found in our RNAseq and ATACseq analyses. Annotating these peaks to within +/- 3000bp of their nearest genes and identifying associated KEGG enriched pathways revealed gains in Wnt signaling at both 0Gy and 2Gy with a loss in peaks associated with apoptosis at 2Gy (Figure 5E). Global increases of H3K27ac were also observed in Dsup expressing cells vs WT for baseline and after irradiation (p<0.0001) with slight decreases in signal following irradiation (Figure 4B). Annotating these peaks regions revealed slight deviations in promoter regions ≤1Kb from the TSS where there was a slight gain in H3K27ac in these areas (Supplemental Figure S21B). Wnt signaling, Hippo signaling, and focal adhesion were among the top significant KEGG pathways marked by H3K27ac in Dsup expressing cells at baseline and post irradiation (Figure 5F).

**Figure 5.**
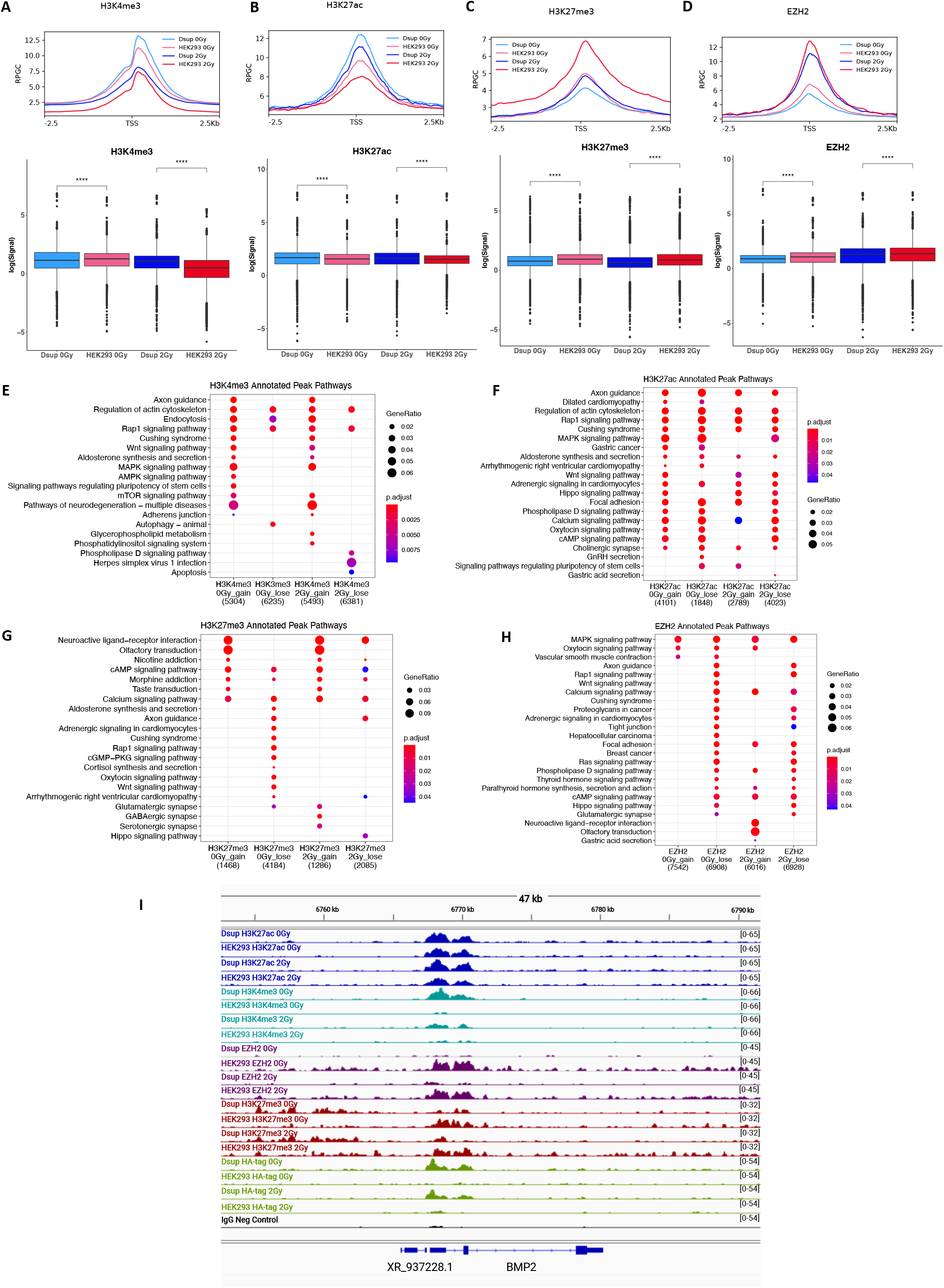
Cut&Run analysis reveals global effects of histone PTM and EZH2 enrichment modulated by expression of Dsup in HEK293 cells. **(A)** Metagene plots (top) and boxplot with log signal (bottom) shows H3K4me3 signal enrichment greater in Dsup expressing cells at baseline and post-irradiation compared to WT after normalizing to 1x genomic coverage. H3K4me3 signal decreases post-irradiation for both HEK293 Dsup and HEK293 WT. **(B)** Metaplot (top) and boxplot with log signal (bottom) shows H3K27ac signal enrichment greater in Dsup expressing cells at baseline and post-irradiation compared to WT after normalizing to 1x genomic coverage. H3K27ac signal decreases post-irradiation for both HEK293 Dsup and WT HEK293 **(C)** Metaplot (top) and boxplot with log signal (bottom) shows H3k27me3 signal enrichment lower in Dsup expressing cells at baseline and post-irradiation compared to WT after normalizing to 1x genomic coverage. EZH2 signal increases post-irradiation for both HEK293 Dsup and HEK293 WT. **(D)** Metaplot (top) and boxplot with log signal (bottom) shows EZH2 signal enrichment lower in Dsup expressing cells at baseline and post-irradiation compared to WT after normalizing to 1x genomic coverage. EZH2 signal increases post-irradiation for both HEK293 Dsup and HEK293 WT. All Statistical analysis was performed using Wilcoxon signed rank test with significant values plotted over bars, p****=p>0.001. **(E)** Differential peak analysis was conducted using RGT-THOR and resulting gains and losses in peaks were annotated using ±3000bp of TSS regions for annotated hg38 promoters. Gene annotations were then used to generate enriched terms using KEGG analysis with p-adjusted cutoff < 0.05 with Bonferroni correction. Dot plots show an enrichment for gains in H3K4me3 peaks associated with Wnt signaling in HEK293 Dsup while peaks associated with apoptosis were lost post-irradiation. **(F)** dot plot for H3K27ac annotated pathways show gains and loses for Wnt and Hippo signaling. **(G)** H3K27me3 annotated KEGG pathways show a loss in Wnt signaling at 0Gy and a loss in Hippo signaling pathways at 2Gy. **(H)** EZH2 annotated KEGG pathways show gains and losses in MAPK signaling pathways while Wnt signaling pathways are lost in HEK293 Dsup and EZH2 marks for focal adhesion pathways are lost at 0Gy and 2Gy, but some associated focal adhesion peaks are gain at 2Gy. Size of dots for dot plots represent the ratio of genes for each pathway in comparison to the entire set while gradient color from red to blue represents p-adjusted values with blue being more significant. **(I)** IGV tracks shows a representative locus on chromosome 20 for top differential peak BMP2. Signal for each histone PTM, EZH2, and Dsup with an HA-tag antibody is plotted for both HEK293 Dsup and HEK293 WT. Here we see increased H3K27ac and H3K4me3 signal for Dsup expressing cells while EZH2 and H3K27me3 are increased in in HEK293 WT cells. Dsup binding signal is also present over these active sites. IgG negative control is included at the bottom in black.

When evaluating repressive marks, H3K27me3 signal was lowest in Dsup expressing cells at baseline and post irradiation (P<0.0001). Both cell types followed a similar increase in H3K27me3 marks post-irradiation but it was far more pronounced in WT cells (Figure 5C). Annotated KEGG pathways revealed a repression of Neuroactive ligand-receptor interactions, cAMP pathways, and calcium signaling in HEK293 Dsup, while Wnt signaling and Hippo signaling were more repressed in the WT cells (Figure 5G). Most H3K27me3 signal was lost around promoter regions 1kb from the TSS (Supplemental Figure S21C). EZH2 followed a similar pattern as H3K27me3 with lower levels of EZH2 signal in Dsup expressing cells for baseline and post-irradiation (P<0.0001) (Figure 4D). Differential peak analysis and annotation revealed about a 30% loss in EZH2 in HEK293 Dsup for both conditions at promoter regions ≤1Kb from the TSS (Supplemental Figure S21D). Consistent with H3K27me3 marks lost in HEK293 Dsup, Hippo signaling and Wnt signaling were not present in HEK293 Dsup but in WT cells and repression of focal adhesion pathways only occurred at 2Gy of radiation (Figure 5H).

When looking at histone PTMs and EZH2 signals plotted over the aggregate of the centers of Dsup peaks, most of the overlap occurs within H3K4me3 signal followed by H3K27ac followed by a slight decrease in signal post-irradiation. There was very little overlap with Dsup binding sites and EZH2 and H3K27me3 signal (Supplemental Figure S22A, S22B). Annotating Dsup peaks with KEGG pathways revealed a preference for pathways associated with collagen synthesis, SUMOylation of DNA replication proteins, and VEGA-VEGFR2 signaling pathways at 0Gy of radiation. At 2Gy of radiation there was an enrichment for VEGF signaling and MAPK signaling (Supplemental Figure S22C). Enrichment analysis with Reactome related terms showed increased laminin interactions, collagen formation, and SUMOylation of DNA replication proteins pathways at 0Gy baseline Dsup binding. Post-irradiation Dsup binding continued to have an enrichment for Receptor Tyrosine Kinase signaling and VEGA-VEGFR2 pathways while RAF/MAP kinase cascade, potassium channel, and MAPK signaling cascade related pathways were uniquely enriched at 2Gy (Supplemental Figure S22C). Enrichment analysis using annotated gene sets from MSigDB revealed a Dsup binding preference to genes belonging to targets of bivalently marked genes and those with just the repressive H3K27me3 marks (Supplemental Figure S22D). When looking at enriched MsigDB pathways for each dose of radiation individually at 0Gy and 2Gy there was a Dsup binding preference to gene sets involved in cell growth, Tp53 targets, and cell differentiation as well as genes that would be downregulated during a UV response (Supplemental Figure S22E & S22F). Annotating Dsup peaks to different genic and intronic features showed a slight preference for promoter regions post-irradiation compared to Dsup binding at baseline (Supplemental Figure S22G).

These findings are in accordance with our transcriptomic and chromatin accessibility data that suggests that bivalently marked PRC2 targets are upregulated in Dsup expressing cells and as such we generated histone PTM and EZH2 signal tracks for the PRC2 target BMP2 for baseline and 2Gy of irradiation (Figure 4E). In addition, we also looked at Dsup binding to this gene and discovered increased Dsup signals overlapping with increased H3K4me3 peaks in HEK293 Dsup while H3K27me3 and EZH2 were decreased in the same area at baseline suggesting a Dsup binding preference for open chromatin marks at PRC2 targets (Figure 5I).

### Direct comparison of HEK293 Dsup with HEK293 HMGN1 over expression and HEK293 Dsup with HMGN1 KO reveal distinct epigenetic and expression profiles

As Mowery et al. first demonstrated that induced HMGN1 over expression in Nalm6 cells led to an overall more transcriptionally active and modified epigenetic state^27^ similar to our own Dsup expressing cells we next sought out a direct comparison between Dsup and HMGN1 over expression in HEK293 cells to see if we could induce similar cell state profiles. Since HEK293 Dsup cells contained the same levels of HMGN1 and HMGN2 as inferred from our expression dataset across time and radiation doses (Supplemental Figure S23A & S23B), we wanted to see if the changed cellular state of HEK293 Dsup was due to introduction of what seems to be an extra copy of a chromosomal architectural protein. In addition, we also knocked out HMGN1 in HEK293 Dsup expressing cells to see if this would rescue the phenotype back to a more WT like cell. We confirmed overexpression of HMGN1 in HEK293 and KO of HMGN1 in HEK293 Dsup cells via western blot (Supplemental Figure S23C).

Our histone PTM data indicated that Dsup induced increases in H3K27ac, H3K4me3, and decreases in the repressive H3K27me3 globally and so we chose these epigenetic modifications to compare our new cell types to. After normalization to spike-in control we observed global increases in H3K27ac with HEK293 Dsup having the greatest increase followed by Dsup HMGN1-KO and then HEK293 HMGN1-OE compared to WT (Anova p<2.2e-16, Figure 6A). H3K4me3 on the other hand was highest in Dsup HMGN1-KO followed by HEK293 HMGN1-OE (Annova p<2.2e-16, Figure 6C). Overall decreases in H3K27me3 were seen in HMGN1-OE, followed by HEK293 Dsup, and then Dsup HMGN1-KO with the largest decrease in this histone PTM compared to WT (Annova p<2.2e-16, Figure 6E).

**Figure 6.**
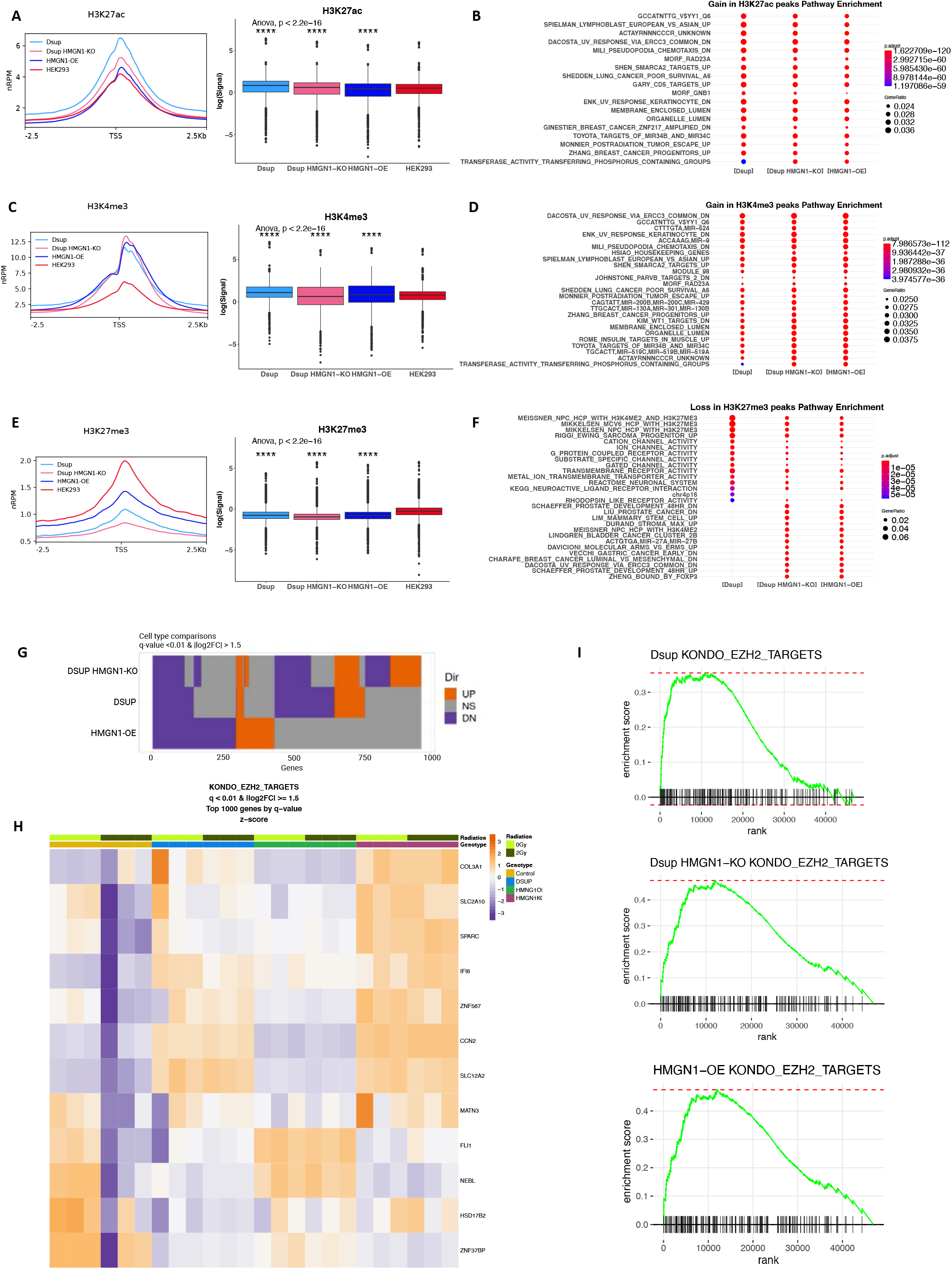
Comparison of HEK293 Dsup, HEK293 HMGN1-OE, and HEK293 Dsup HMGN1-KO reveals similar global modulations of transcriptomic and epigenetic landscape. **(A) (C) (E)** Metagene plots center histone PTM Cut&Run normalized reads per million 2.5kb downstream and upstream of GRCh38 transcription start sites (left). Boxplots represent the distribution of the log of Cut&Run signal at each gene. All Statistical analysis was performed using Wilcoxon signed rank test with significant values plotted over bars, ns=p>0.05, **p<0.05, ***p<0.01, ***p<0.001, ****p<0.0001. Global Significance was also calculated by ANOVA analysis followed by Tukey’s HSD post hoc test for comparisons of all cell types to control HEK293 cells. **(B) (D) (F)** Dotplots were used for visualization of functional annotation of enriched terms after RGT-THOR differential peak analysis. Top differential peak pathways were obtained for HEK293 Dsup, HEK293 Dsup HMGN1-KO, and HEK293 HMGN1-OE were obtained via Enricher with pathways obtained from msigdb.v7.1. with q-value cutoff of 0.05 and Bonferroni correction. Color gradient legend displays p-adjusted values and size of dots represents the ratio of genes for each pathway belonging to total gene set. **(G)** Altuna plot reveals comparisons of differentially expressed genes for each cell type with x-axis representing total gene counts with approximate counts visualized for each up/ns/down groups by their relative sizes, q-value < 0.01 & absolute value of Log_2_FC ≥ 1.5. **(H)** Heatmap clustering for radiation dose and genotype shows expression of top leading edge differentially expressed genes belonging to KONDO_EZH2_TARGETS as assessed by transcriptomic Fgsea analysis, q-value <0.01 & Log_2_FC ≥ 1.5. **(I)** Running score for preranked gene lists assessed via Fgsea enrichment distribution plots of transcriptomic data for each cell type for KONDO_EZH2_TARGETS.

Differential peak analysis was performed for each cell type and histone PTM using RGT-THOR. Permutation tests were performed for differential peaks called and then compared to one another to test the null hypothesis that peak distributions for HEK293 Dsup vs HEK293 HMGN1-OE, HEK293 Dsup vs HEK293 Dsup HMGN1-KO, and HEK293 HMGN1-OE vs HEK293 Dsup HMGN1-KO were the same. There was overlap among H3K27ac and H3K4me3 differential peaks for the three cell types (Supplemental Figures S24A & S24B); however, only HMGN1-OE vs Dsup H3K27me3 had overlap while Dsup HMGN1-KO vs Dsup H3K27me3 had different peak distributions (p=0.001). The same was seen for the comparison of HMGN1-OE vs Dsup HMGN1-KO H3K27me3 (p=0.001) (Supplemental Figure S24C). After performing differential peak analysis, functional annotation of gains in H3K27ac showed shared chromatin remodeling pathways such as those belonging to SHEN_SMARCA2_TARGETS_UP and radiation response pathways such as MORF_RAD23A, ENK_UV_RESPONSE_KERATINOCYTE_DN, and MONNIER_POSTRADIATION_TUMOR_ESCAPE_UP (Figure 6B, Supplemental Figure S25). KEGG annotation of peaks called by MACS2 highlighted some of the differences among the cell types with H3K27ac peaks belonging to increased focal adhesion seen in only the Dsup expressing cells. Among shared pathways not seen in WT cells, Wnt, Ras, and MAPK signaling were some of the top hits (Supplemental Figure S28A). Peak annotations for each cell type also revealed a slightly greater percentage of H3K27ac peaks around promoter regions in HEK293 HMGN1-OE cells (Supplemental Figure S28B). Further analysis into associated focal adhesion PRC2 target histone PTM marks revealed a lack of transcriptionally permissive H3K27ac and H3K4me3 marks in HEK293 HMGN1-OE cells for the collagen synthesis gene P4HA2 while HEK293 Dsup and HEK293 Dsup HMGN1KO had similar amounts of H3K27ac but Dsup expressing cells with HMGN1-KO had higher levels of H3K4me3 after normalization.

Interestingly, H3K27me3 levels remained low in HMGN1-OE cells compared to WT (Supplemental Figure S28G). In comparison, BMP2, a PRC2 target not related to focal adhesion showed strong H3K27ac and H3K4me3 signals for Dsup expressing cells and HMGN1-OE cells while WT HEK293 had repressive H3K27me3 marks. In both cases Dsup expressing cells with HMGN1-KO had the highest H3K4me3 signal, while HEK293 Dsup with HMGN1 intact had the stronger H3K27ac signal (Supplemental Figure S28H).

Similar shared gene set enrichments were also seen in H3K4me3 differential peak pathways for the different cell types (Figure 6D, Supplemental Figure S26). Among MACS2 peak calls for the individual cell types, KEGG annotations revealed a similar cell type distribution where only Dsup expressing cells had increased H3K4me3 peaks related to focal adhesion and Dsup expressing cells had the greater percentage of H3K4me3 around promoter regions (Supplemental Figures S28C & S28D). Among H3K27me3 peaks gene set enrichments, losses in pathways belonging to histone core proteins associated with bivalent H3K4me2 and H3K27me3 marks were lost in Dsup expressing cells and HMGN1 over expressing cells as well as pathways enriched for receptor and channel activities (Figure 6G, Supplemental Figure S27) H3K27me3 KEGG annotated pathways were mainly enriched for gene sets belonging to Neuroactive ligand-receptor interactions for all cell types, while Focal adhesion, Rap 1 signaling, and regulation of actin cytoskeleton H3K27me3 peaks were enriched only in HEK293 cells (Supplemental Figure S28E & S28F).

As we have demonstrated a more transcriptionally active state particularly at bivalently marked genes in HEK293 Dsup cells and evidence from the literature has demonstrated increased transcriptional activity in HMGN1 over expressing cells at similar regions^27, 88, 90^, we next wanted to compare the transcriptional states of HEK293 Dsup, HEK293 HMGN1-OE, and HEK293 Dsup HMGN1-KO. Here we performed whole transcriptome sequencing on the different cell types for both baseline and 2Gy of radiation since we wanted to see if the radioprotective properties of HEK293 Dsup with HMGN1 Knocked out was perturbed and if HMGN1 over expression in HEK293 cells could potentially confer increased radio-tolerance to HEK293 cells. We began first by clustering the transcriptional states of each cell type via PCA and t-SNE hierarchical clustering. HEK293 Dsup and HEK293 Dsup HMGN1-KO both clustered together for both baseline and 2Gy of radiation while HEK293 HMGN1-OE clustered tightly together for both 0Gy and 2Gy of radiation. WT HEK293 at baseline formed its own cluster while WT HEK293 at 2Gy of radiation formed its own cluster. 54.59% of variance was explained along PC1 and 23.88% of variance along PC2 (Supplemental Figures S29A & S29B).

Next, we plotted overall up vs down regulation of genes via Altuna plot and discovered HEK293 Dsup with HMGN1 KO had the greatest amount of upregulation of genes followed by HEK293 Dsup and last HEK293 HMGN1-OE with the majority of gene sets shared between HEK293 Dsup HMGN1-KO and HEK293 Dsup (q-value <0.01 & absolute value Log_2_FC ≥ 1.5) (Figure 6H). There was greater overlap of downregulated genes among the three cell types with 115 inclusive gene counts while inclusive gene counts for upregulated genes was just 23. (Supplemental Figure S29C). FGSEA was then performed on differentially expressed genes for each cell type and dose of radiation and top upregulated and downregulated pathways were obtained (Supplemental Figure S30A). KONDO_EZH2_TARGETS was among the shared pathways that was upregulated in Dsup expressing cells and HEK293 HMGN1-OE with HEK293 Dsup HMGN1-KO and HEK293 HMGN1-OE having the greater enrichment scores (Figure 6I). Plotting top genes via heatmap revealed another upregulation of collagen forming Polycomb targets such as COL3A1 only present in Dsup expressing cells with stronger enrichment signal in HEK293 Dsup HMGN1-KO. HEK293 HMGN1-OE clustering looked more similar to WT cells for this pathway except after radiation where HMGN1-OE cells had stronger signal for FLI1 and NEBL, which were downregulated in HEK293 at 2Gys and in Dsup expressing cells at baseline and radiation (Figure 6H). Analyzing Fgsea pathways with with radiation normalized revealed upregulated shared pathways among Dsup expressing cells and HEK293 HMGN1-OE cells included MIKKELSEN_MEF_HCP_WITH_H3K27ME3, BENPORATH_SUZ12_TARGETS, and BENPORATH_EDD_TARGETS. Top shared downregulated pathways between HEK293 Dsup HMGN1-KO and HEK293 GNF2_APEX1 (Supplemental Figures S30B & 30C). When analyzing these same gene sets with REACTOME enrichment terms WNT related signaling and histone demethylase activity pathways were upregulated among the three cell types (Supplemental Figures S30D, S30E & S30F).

## Discussion

Chromosomal aberrations due to GCR-induced DSBs can be detrimental to future long-duration space flight missions. In the NASA Twins Study data, a greater rate of chromosomal inversions was found during space flight, and after, indicating genomic instability and rearrangement^8^.

Since the tardigrade Dsup protein has been shown to suppress DNA damage we tested if a stable expression of Dsup in a human cell line could also sustain such protective phenotypes, and if there would be other changes to human gene expression networks. Overall, the results validate the functional radioprotective capacity of Dsup in human cells, to establish the stable line for use by others, and provide new molecular details about the functional genomic impact of Dsup integration into the human genome.

We integrated Dsup under a CMV promoter into the HEK293 genome and then validated its radioprotective properties through a previously unexplored DNA damage biomarker assay in the context of Dsup expressing cells. Here we looked at how well Dsup could reduce 8-OHdg bulky adducts as this is a common and accessible biomarker of radiation-induced ROS stress. As we were able to measure a significant increase in 8-OHdg in the NASA Twin Study during space flight^8, 52^, our aim was to then to see if Dsup could reliably mitigate some of this damage for perhaps future space exploration missions. Here we observed significant reductions up to 4Gy of radiation at 1hr post irradiation and at 24hrs later a good portion of the damage was cleared.

While baseline 8-OHdg is suppressed at 0Gy of radiation, the utility of baseline 8OHdG in the genome has yet to be explored. It has been proposed that endogenous 8OHdG may play a role in antioxidant and anti-inflammatory mechanisms^56^. Oxidative stress plays a complex role in cell signaling and when in excess can be highly damaging to macromolecules. However, when expressed at normal levels are important for activation of tyrosine kinases, MAPK, and Ras proteins^54, 55^. While Dsup thus far has been shown to reduce damage typically repaired by nonhomologous end joining and homologous recombination^20^, we hypothesized that whether Dsup has an exclusive shielding effect from ROS damage or also a DNA repair enhancing effect like HMGN1, it would be likely that Dsup could also prevent the accumulation of 8-OHdg in cells. Indeed, it has been shown that HMGN1 also has a direct effect on regulating BER through stimulating PARP-1 activity^94^

Continuing off the work of Chavez and colleagues where they first discovered the similar core RRSARLA amino acid sequence shared between Dsup and the vertebrate HMGN1 nucleosome binding domains^19^; we tested if Dsup may behave in a similar manner as HMGN1 has been demonstrated to confer radioprotective properties^92^. We began with an exploratory analysis of the HEK293 Dsup transcriptome and discovered perturbations in gene expression networks similar to that of an HMGN1 over expression genotype^27^. We opted to include time and radiation dose in our experimental design to gain a more in-depth multivariate understanding of the impact of introducing a foreign gene into a whole new system. To our surprise, hierarchical clustering revealed gene expression changes were mainly driven by the presence of Dsup itself, far outweighing the impact of time and then radiation even at higher doses. In addition, clustering via Principal Component Analysis did not reveal particularly strong separation in variance for our gene expression data; unlike PCA for our chromatin accessibility data which did show a clearer delineation between Dsup expressing cells and WT. This implicates Dsup as having its strongest effect on altering the epigenome, which would not be unfeasible as Chavez et al. demonstrated Dsup’s proclivity towards nucleosome binding^19^, potentially similar to that of HMGN1’s role as a chromatin architectural protein that scans the epigenome and binds transiently on and off nucleosomes to affect a number of DNA dependent processes including transcription, recruitment of DNA repair factors, and chromatin decompaction which enables chromatin accessibility to histone modifying complexes^21, 22, 26^.

Our own analysis indicated an overall more transcriptionally active HEK293 Dsup cell type as compared to WT across time and radiation doses. Fast Gene Set Enrichment analyses of these significant differentially expressed genes revealed an upregulation of typically lowly expressed genes belonging to Polycomb Repressor Complex 2 (PRC2) protein targets. This key finding demonstrated that HEK293 Dsup cells may have a genotype similar to that of HMGN1 over-expressing cells such as that seen in B cell acute lymphoblastic leukemia where there is a global amplification of gene expression and modulation of chromatin compaction. In particular, Mowery et al. showed that within their own HMGN1-OE in-vitro models, genes that were silent in WT cells remained silent but those that were lowly expressed belonging to PRC2 target pathways were enriched for^27^. PRC2 targets are bivalently marked genes with the active H3K4me3 and repressive H3K27me3 histone PTMs at promoter regions typically associated with lineage defining genes and as such are poised for higher expression depending on cellular context such as chromatin decompaction^27, 101, 102^.

KEGG, ENRICHR, and GO enrichment analysis on these differentially expressed genes also revealed an upregulation of several transcriptionally permissive pathways at baseline and pathways conducive to chromatin remodeling; while pathways involved in kidney cell development were downregulated post-irradiation. Interestingly, when taking a closer look at the top upregulated genes belonging to pathways that overlapped with PRC2 targets, many were involved in neuronal development pathways and became more enriched for when normalizing for time and radiation. This finding points to a potential overlap in function with HMGN1 as loss of HMGN1 has been shown to enhance the rate of OSKM induced reprogramming of mouse embryonic fibroblasts into iPSCs due to destabilization of the epigenetic landscape while overexpression of HMGN1 has been shown to affect lineage and maturation specific transcriptional programs ^23, 27^. As HEK293 cells may have originated from adrenal precursor cells with a neural crest origin during development^106, 107^, it could be possible that epigenetic perturbations induced by Dsup may further relieve some of these typically suppressed PRC2 targets that could lead to this neuronal type differentiation.

With what looks to be a more transcriptionally active and relaxed chromatin state, we next sought out to probe the epigenetic landscape to confirm our findings. While no significant changes are known to occur in chromatin accessibility after induction of HMGN1 over expression in Nalm6 cells^27^, in HEK293 Dsup we were able to see overall chromatin stability when compared to WT but also were able to track opening and closing dynamics of ATACseq peaks over time and with a 2Gy dose of irradiation with some greater resolution. Here, we were able to observe a slightly more open chromatin state in gene TSS 2.5Kb away from the nearest enriched ATACseq peak while gene TSS closest to the nearest peak under 100Bp away were generally in closed regions. When plotting open chromatin signal over all TSS of genes belonging to GRCh38 we observed generally greater enrichment of peaks in Dsup expressing cells over time but with a drop off in signal post-irradiation and Dsup chromatin peaks appearing more closed at 6hrs. This pattern of opening and closing of peaks appears to be in general agreement with our analysis of gene expression dynamics over time with more downregulated genes appearing at the 6hr timepoint as well. Closing of chromatin post-irradiation has also been well documented as mammalian cells will modulate histone PTMs around the sites of damage to facilitate a repressive chromatin state that limits local transcription so that the steps necessary to begin DNA damage repair can proceed first^104^. Although chromatin accessibility in our dataset were generally stable like that of Nalm6 HMGN1-OE cells, the nuances we observed in opening and closing chromatin state dynamics could be due to our constitutive expression of Dsup over several passages while Mowery et al. looked at chromatin state dynamics 6hrs after induction of HMGN1 over expression. It would then be worthwhile to see at some point if long term constitutive over expression of HMGN1 would lead to a similar chromatin landscape, or if perhaps short-term induction of Dsup would lead to less chromatin state perturbations which could provide for temporally controlled radiotolerance while mitigating the risks associated with epigenomic instability.

Our proposed mechanism of how Dsup improves radiotolerance is through modulation of the epigenetic landscape. While the prevailing hypothesis is that Dsup binds to nucleosomes to shield against ROS^18, 19^; our results indicate a more open and permissive chromatin state for less steric hindrance of repair machinery to be recruited to the sites of damage much in the same way HMGN1 acts to reduce the compaction of higher-order chromatin structure around sites of DNA damage^22^. However, it should be noted that these two mechanisms are not incompatible with each other, and that we are merely adding another level of nuance to how Dsup functions but in the context of an *in-vitro* model. It’s been demonstrated that nucleosomes themselves provide a shielding effect against ROS damage as opposed to nucleosome free regions near promoters of actively transcribed genes. This protective effect is most prevalent at sites where nucleosomes occupy for extended periods of time and when HMGB1 in mammalian cells is mutated or knocked out there is a reduction in nucleosome occupancy that is correlated with higher rates of double stranded breaks in MCF-7 cells^108^. Our own data suggests greater nucleosome occupancy and longer nucleosome fragment length surrounding the TSS of active promoters in Dsup expressing cells and a decrease in occupancy following 2Gy of radiation. Longer fragment length distributions in HEK293 Dsup could support the shielding effect hypothesis as Dsup is larger than HMGN1 or any of the other High Mobility Group proteins and so perhaps greater portions of nucleosome associated DNA could be protected from ROS damage. Increases in nucleosome occupancy could be counterintuitive to the theory that Dsup promotes a more relaxed chromatin landscape; however, HMGN1 has also been shown to be involved in affecting the stability and positioning of nucleosomes, particularly around the TSS of nucleosomes residing in transcriptionally active genes with promoters within CGI regions^109^. In addition, there is a correlation in ESCs where HMGN1 levels influences nucleosome occupancy, and such transcriptional output is also upregulated at these nucleosomes^109^. HMGN1 occupied nucleosomes are more susceptible to micrococcal nuclease digestion as opposed to nucleosomes occupied with histone H1 as they make up more compact chromatin regions and thus would take longer to digest^109^. Interestingly, CGI promoters typically are associated with ubiquitously expressed genes and developmental genes marked by Polycomb group repressor proteins^110^. As such, HMGN1 occupied nucleosomes within these regions have enhanced levels of transcription as nucleosomes at the TSS are destabilized but occupancy is increased at the +1 site. When plotting nucleosome occupancy signal over CGI promoters we observed a similar phenomenon with stronger signal within these regions for Dsup expressing cells at baseline but decreased signal post-irradiation. During initial repair processes, chromatin remodeling complexes are recruited to DSBs and can remove the steric hinderance that nucleosomes may impose^103^ and hence why we see a drop in nucleosome enrichment post irradiation initially.

As HMGN1 preferentially localize to chromatin regulatory regions with permissive histone post translational modifications^23^ and consequently impacts higher order chromatin structure^25^, gene expression networks^23, 27^, and modulate DNA damage repair machinery accessibility^22, 29, 35, 38^, the next question was to address how Dsup might shape the HEK293 global histone PTM landscape given our current data. HMGN1 is known to colocalize to active chromatin marks such as H3K4me3 and H3K27ac while being absent from genomic sites enriched in condensed and transcriptionally silent regions containing H3K27me3^23^. Interestingly, when HMGN1 is overexpressed as in the case with HMGN1 induction in Nalm6 cells there is a global increase in H3K27ac and promoter associated H3K4me3 while there was a significant decrease in H3K27me3 that led to an upregulation of lineage and leukemia specific pathways as well as EZH2 targets, the catalytic subunit of PRC2^27^. These results could be expected as HMGN1 is known to compete with the repressive linker histone H1 for nucleosome binding sites and thus recruit or repel enzymes that enact histone PTMs^25, 26^. In our own study we observed global increases in these active marks H3K4me3 and H3K27ac primarily occurring in promoter regions while there were overall decreases in H3K27me3 and EZH2. When plotting these histone PTM and EZH2 signals over peaks called where Dsup was bound we also observed a strong overlap in active histone PTMs and Dsup presence while Dsup was nearly absent wherever H3K27me3 and EZH2 was present. Further annotation revealed an enrichment for Dsup belonging to pathways associated with bivalently marked PRC2 targets. Altogether, this data supports our hypothesis that Dsup in HEK293 cells binds to nucleosomes with active histone PTM marks to promote a more open chromatin state much like HMGN1, but also in the context of a Dsup acting like an HMGN1 over expression, can also affect histone PTMs globally to perturb lineage specific networks. Although this proposed mechanism scratches the surface of how Dsup may interact with the chromatin landscape to induce a more relaxed chromatin state, another angle to consider is how exactly does Dsup interacts with histone H1 to bring about these changes. While biochemical assays demonstrated that Dsup and H1 can bind to nucleosomes simultaneously^19^, the same was observed in mobility shift assays where HMGN1 and H1 also bound simultaneously. It was discovered; however, that while H1 was not displaced from the nucleosome^19^, its c-terminal domain responsible for nucleosome dependent condensation and stabilizing higher order chromatin structures was altered by HMGN1 to enable a more permissive state^111^. To better understand the mechanism for which Dsup alters higher level chromatin structure and recruit active mark histone PTMs it would be worthwhile then to study the interactions of the highly basic H1 C-terminal domain with Dsup in an in-vitro model.

Last, we asked the question is the resulting perturbed epigenetic and transcriptomic states of the introduction of Dsup into the HEK293 genome a result of what seems to be an overexpression of chromosomal architectural proteins? If this is the case, what would be the resultant cellular state of knocking out HMGN1 in Dsup expressing cells, and perhaps could we see the same perturbed cellular state in HEK293 HMGN1 over expressing cells? Surprisingly, it seems that Dsup does indeed promote its own unique influence over cellular state irrespective of HMGN1 knock out. Rather than reverting to a more WT cellular state, KO of HMGN1 in some cases even produced a more transcriptionally permissive state with the loss of H3K27me3 peaks and increased global gains H3K4me3 peaks after normalization to spike-in control. HMGN1 over expression in WT cells led to similar epigenetic and transcriptomic profiles first documented by Mowery and colleagues in Nalm6 cells^27^; however, while some overlap existed among the three cell types, Dsup expressing cells had its own unique epigenetic and transcriptomic signatures. Among shared GSEA annotated enriched pathways after differential peak calling there was a gain in H3K27ac and H3K4me3 peaks related to radiation responses and chromatin remodeling while resultant loss in H3K27me3 peaks were mainly enriched at bivalently marked Polycomb target regions. KEGG annotation after MACS2 peak calling revealed shared pathways among the cell types such as Wnt and Hippo signaling but also highlighted enriched pathways unique to Dsup expressing cells such as increased focal adhesion. When viewing signal tracks for genes belonging to Polycomb targets, Dsup expressing cells and HEK293 HMGN1-OE had similar histone PTM signals for BMP2 with HEK293 Dsup HMGN1-OE having a higher H3K4me3 signal while HEK293 Dsup had a higher H3K27ac signal among the different cell types. When viewing histone PTM signal over P4HA2, a collagen synthesis gene related to focal adhesion and also a Polycomb target, only strong H3K27ac and H3K4me3 signal could be seen for Dsup expressing cells while very little signal was visible for HEK293 HMGN1-OE and WT, indicating that increased focal adhesion may be a novel unexplored consequence of Dsup introduction into the human genome. Further research could be done into exploring perhaps wound healing therapeutic benefits of increased focal adhesion upon Dsup induction^99, 117^.

Further analysis into the transcriptomic states of these cell types also revealed a similar transcriptionally permissive state among Dsup expressing cells and HEK293 HMGN1-OE with an increased overlap of upregulation of Polycomb targets such as those belonging to the KONDO_EZH2_targets first reported as upregulated in Nalm6 cells^27^. Here we saw an overall upregulation of genes with Dsup expressing cells with HMGN1-KO surprisingly having the greatest amount of upregulation of genes. While it’s been documented that HMGN2 may have similar redundant nucleosome binding properties and functionality as HMGN1 and may compensate for a loss of HMGN1^35, 118^, our own expression analysis showed relatively stable expression among the different cell types for HMGN2 with no increased expression in HEK293 Dsup with HMGN1-KO, thus ruling out a potential chromosomal architectural protein over expression in this cell type. If there are no potential compensatory mechanisms at play here, then perhaps a KO of HMGN1 may allow for more Dsup binding to nucleosomes as its only been demonstrated thus far that Dsup and histone linker H1 can bind to the same sites on nucleosomes simultaneously, but competition for nucleosome binding sites has still yet to be explored with HMGN1 vs Dsup, despite having similar nucleosome binding domains^19^. In contrast to the hypothesis that introduction of Dsup may perturb the cellular state as an over expression of a chromosomal architectural protein, it may be likely that Dsup just has stronger binding affinity to nucleosomes than HMGN proteins. A recent computational model explored the strong electrostatic interactions and flexibility of Dsup binding with nucleosomal DNA due to its intrinsically disordered structure that allows for tight aggregates to form between Dsup’s highly abundant positive charges in its c-terminus and the negative charges of DNA phosphate backbones^119^. Further *in situ* or *in vitro* experiments could elucidate if there is competition for binding sites between these two proteins, and if so the rate of residence on nucleosomes can also be distinguished if in fact Dsup binds stronger of for longer periods of time.

In the first results section we revisited some of these pathways of interest that appeared often in our multi-omic analyses and confirmed these unique phenotypes as many of these gene sets also overlapped with bivalently marked PRC2 targets. Here we tested a series of assays to validate the enhanced proliferative, anti-apoptotic properties, and adhesive properties of HEK293 Dsup cells. As we saw an enrichment of Hippo and Wnt signaling pathways we thought that this might explain some of the increased survival of Dsup expressing HEK293 cells first observed by Hashimoto et al^10^ as these pathways are often involved in increased cell proliferation and anti-apoptosis^46, 49, 57, 60^. In addition, Hippo and Wnt signaling dysregulation have also been shown to increase radiotolerance of cancer cells through their role in promoting DNA repair and inhibiting apoptosis. Although it’s likely that HEK293 Dsup’s reduction in apoptotic signaling could be due in part to the shielding hypothesis and that apoptotic signals are reduced because damage was prevented, it is a worthwhile research question to explore whether Wnt and Hippo signaling may play a role in this phenomenon and if these perturbations in these signaling pathways are brought about via chromatin remodeling^95, 115^.

Collagen formation and focal adhesion in general were some of the top upregulated pathways revealed by enrichment analysis. Although not a widely reported phenotype, elevated levels of HMGN2, a high mobility group protein shown to have redundant properties of HMGN1 when HMGN1 is knocked out^35^, has been shown to exhibit upregulation of cell adhesion pathways in MEF^112^. In an *in-vivo* model, increased collagen formation was found to be correlated with increased HMGN1 expression in mouse kidney cells and was suggested to play a role in renal fibrosis^113^. Increased focal adhesion in the context of HEK293 HMGN1-OE cells in our own multi-omic data was not observed. However, we confirmed the increased adhesive properties of HEK293 Dsup on Collagen I, Collagen IV, and fibronectin substrates at baseline and post-irradiation. The exact mechanism of how the HMGN1 proteins regulate these adhesive properties remains to be elucidated but given the context of how HEK293 Dsup cells have an altered chromatin landscape and an upregulation of collagen synthesis PRC2 targets, such as P4HA2^114^, there could be a link between chromatin architectural proteins and the extracellular matrix that remains to be further investigated.

Given these broad results, one can imagine Dsup and related genes helping in the creation of a temporal, radiation, or molecule controlled, inducible CRISPR-dCas9-Dsup system that is activated in response to intense periods of galactic cosmic radiation. Indeed, such work has already begun for atomic energy workers and in other trials of engineered cancer therapies that leverage CRISPR^85^. Even if not directly used as a therapeutic, the gene expression perturbation data here, as well as the stably expressing cell line, can serve as resources to guide future research on radioresistance and baseline levels of transcriptional disruption after the stable integration of a radioresistance gene. Together, such tools and data sets will help keep astronauts safer as missions move farther from Earth, become riskier, and are laden with ever-increasing levels of GCR and irradiation.

### Experimental Procedures Plasmids

The complete coding sequence for DSUP from *Ramazzottius varieornatus* as described in Hashimoto et. Al was obtained from The Kumashi Genome Project database. This sequence was cloned into a custom pLVX Puro plasmid lentiviral vector with a constitutive CMV promoter.

Dsup was ligated into the 5’ EcoR1 and 3’ XbaI in the multiple cloning site of pLVX Puro. GeneWiz^®^ Gene synthesis services was used to codon optimize DSUP for expression in mammalian systems. An influenza Hemagglutinin tag was synthesized on the N-terminal region of DSUP to minimize interference of c-terminal binding of DSUP to nucleosomes and for downstream antibody detection. A HMGN1 overexpressing cell line was generated by synthesizing the coding sequence for HMGN1 with a HA tag into the pLVX puro vector. The empty vector plasmid was generated by cutting Dsup out of pLVX puro at the EcoRI and XbaI and ligating in a new mcs site. HMGN1 KO plasmid was purchased from GenScript^®^ with the gRNA target sequence CGCACTCACCTCTTCCTTGG and cloned into a pLentiCRISPR v2 vector.

### Cell lines

The plasmids were transformed into DH5α competent cells, colonies were selected, restriction enzyme digested for confirmation of proper cut sites, and then purified plasmid was obtained from maxiprep. HEK293 and HEK293T cells donated by Dr. Levitz at Weill Cornell were grown and incubated at standard incubation conditions at 37°C with 5% CO_2_ in complete DMEM media supplemented with 10% FBS, and 1% Pen/Strep. Mirus TransIt^®^ Lentivirus System was used to transfect HEK293T cells. TransIT^®^ Lenti Reagent was warmed to room temperature and vortexed gently before using. 200ul of Opti-Mem™ media was added to sterile 1.5ml Eppendorf microcentrifuge tubes while 1μg of transfer plasmid was combined with 5μg of lentivirus packaging mix. The plasmid mixture was then transferred to the tube containing Opti-MEM.

30ul of TransIT^®^-Lenti Reagent was added to the diluted plasmid DNA mixture and incubated for 10mins at room temperature. The mixture was then added dropwise to different areas of the plate and gently rocked back and forth to ensure even distribution. Plates were stored at 37°C with 5% CO_2_ for 48hrs before packaged lentivirus particles were harvested. Viral particles were filtered through a 10ml syringe with a 0.45um PVDF filter and separated into 1ml aliquots to be flash frozen and stored in -80°C. TransduceIT™ Reagent was added to HEK293 cells along with 1ml of viral particles distributed evenly over the plate dropwise. Cells were incubated for 48hrs before being selected with 3ug/ml of puromycin for all cells transduced with pLVX puro plasmids. After 1 week of selection the cells were expanded on T-75 flasks.

Confirmation of DSUP expression was assessed via western blot with primary R&D Systems recombinant rabbit HA Tag primary antibody (MAB0601) at a 0.1ug/ml dilution. Bio-Rad Mouse anti Human Vinculin antibody (V284) was used as a control at a 1:500 dilution.

Invitrogen Goat anti-Rabbit IgG (H+L) Cross-Absorbed Secondary Antibody DyLight 800 (SAS-10036) and Rabbit anti-Mouse IgG (H+L) Cross-adsorbed Secondary Antibody Alexa Fluor Plus 680 (A32729) diluted to working concentrations of 1:5000 dilution and 0.1ug/ml respectively in 1% milk in 1X TBST was used to image fluorescence on a Li-Cor Odyssey^®^ CLx imagine system using 700nm and 800nm channels.

Dsup integration site into the HEK293 genome was identified via Oxford Nanopore long read sequencing. Briefly, high molecular weight DNA was obtained from HEK293 Dsup and WT control using Trizol extraction and prepped using Ligation Sequencing Kit (LSK-109). Adapter ligated gDNA was loaded on a PromethION flow cell and base called in real time using Guppy 3.4. The data was analyzed using FindGenomics digital karyotyping service. Genome coverage for HEK293 was 2.8x and for HEK293 Dsup was 1.1x.

### Cell Irradiation

Cells were incubated overnight post seeding at 37°C with 5% CO_2_ before irradiation. The following day cells were taken out of the incubator and irradiated with X-rays in a Rad Source Technologies RS 2000 Biological research irradiator at 0.5 Gy, 2Gy, and 4Gy while the 0Gy plate was kept outside the incubator as a no treatment control.

### Cell Proliferation and Growth arrest

Abcam’s ab126556 BrdU Cell Proliferation ELISA Kit was used to determine growth rate after exposure to 2Gy of irradiation compared to 0Gy controls. Each cell type was seeded at 2 x 10^5^ cells/ml in 96 well plates and 100 µl of complete DMEM media were added to empty wells as controls along with cells that would receive no BrdU reagent in order to assess assay background. 20 µl of diluted 1X BrdU label were added to sample wells and the cells were incubated for 24 hrs. Media was aspirated and cells were then fixed and DNA denatured by incubating in fixing solution at room temperature for 30 mins. Fixing solution was then aspirated and the wells were washed three times with 1X wash buffer before 100 µl of anti-BrdU monoclonal Detector Antibody was added to appropriate wells and incubated for 1hr at room temperature. 100 µl of 1X Peroxidase Goat Anti-Mouse IgG conjugate was then added to the plate and incubated for 30 mins at room temperature. A final wash step was performed three times before addition of 100 µl of TMB Peroxidase Substrate and incubated for 30 mins at room temperature. 100 µl of stop solution was added and the wells and the plate was immediately assayed on a Promega™ Glowmax® Plate Reader with an asborbance set to 450 nm.

### Apoptosis

Live cell real time monitoring of apoptosis via annexin V binding was conducted using Promega RealTime-Glo^™^ Annexin V Apoptosis Assay. Cells were seeded in 96 Well White Polystyrene microplates at a density of 1,000 cells/well and incubated at 37°C with 5% CO_2_ overnight. 100ul of complete medium were added to wells not containing cells as no cell and no compound controls. The following day cells were subjected to X-ray irradiation as indicated above.

Apoptosis was assessed at 1hr, 6hrs, and 24hrs post irradiation. To prepare the 2X detection reagent, 1000X Annexin NanoBiT^®^ substrate was added to prewarmed DMEM complete medium at a 500-fold dilution. 1000X CaCl_2_ was then added to combined DMEM complete medium with diluted NanoBiT^®^ substrate at a 500-fold dilution. 1000X Annexin V-SmBiT and 1,000X Annexin V-LgBiT was added at a 500-fold dilution to the combined substrate, CaCl_2,_ complete medium and was inverted gently a few times to mix. 100ul of the 2X detection reagent was then added to each 96 well plate using a multichannel pipette and mixed at 700rpm for 30 seconds on a plate shaker. Luminescent output were measured on a Promega™ Glowmax® Plate Reader. Significance was calculated by ANOVA analysis followed by Tukey’s HSD post hoc test for comparisons of all cell types to control HEK293 cells, ns=p>0.05, *p<0.05, p<0.01, p<0.001.

Apoptosis as assessed by caspase-3 signal was obtained by resuspending cells in FluoroBrite DMEM supplemented with 10% FBS and 1% 100x glutamax. Cells were counted in a TC20™ Automated Cell Counter. Based on the count of live cells, each cell line was diluted to 75 cells/µL. 200 µL of this mix was added to each well of a Corning 3603 96 well microscopy grade plate for a total of 15,000 cells per well. The plate was put in a humidified 37°C incubator with 5% CO_2_ for 4 hours to allow cell adherence to the bottom of the well.

After cell adherence, plates were removed from the incubator and inoculated with 10 µL of a mix containing 105 µM Hoechst 33342 (diluted from Thermo H3570), 26.25 µM Caspase dye (Diluted from Sartorius 4440), and 277.2 µM Camptothecin (diluted from Millipore Sigma C9911) in water. This inoculation amounts to 5 µM Hoechst 33342, 1.25 µM Caspase dye, 13.2 µM Camptothecin in the well. The plate was sealed with a Breathe-Easy® sealing membrane before incubating for 30 min in the same incubator described above. After incubation, cells were placed in a humidified 37°C ImageXpress Micro Confocal High-Content Imaging System with 5% CO_2_. All wells of the plate were imaged with a 10x Plan Apo Lambda lenses in wide field mode. 9 fields from each well were collected with a 4 ms DAPI exposure and a 93 ms FITC exposure. This process was repeated every 45 min for 22.5 hours (30 time points).

Images were analyzed using MetaXpress. Briefly nuclei were segmented using the built-in find round objects function applied to the DAPI channel. The local background of these objects was defined by using the expand objects without touching function (with the ultimate option checked). The difference between the expanded objects and the nuclei was used to define the background area. The average FITC intensity was calculated for the background and for the nuclear regions. The difference between the nuclear and background FITC intensity was calculated and used as the Caspase signal for each cell at each time point. Upon inspection of the distribution, a cutoff value of 650 caspase signal units was determined to be an appropriate cutoff for cell death as there was a bimodal break around that signal intensity. Cells with lower than 650 units of caspase signal were called non-apoptotic, cells with greater than 650 units of caspase signal were considered apoptotic. The percentage of apoptotic cells was calculated for every well at every time point. These points were plotted for all wells tested for each cell line. A bar graph was made with error bars at 1 standard deviation of the apoptotic rate of the cells at the final time point (22.5 h post inoculation). A heteroscedastic two tailed student t-test was performed to test the null hypothesis that the Empty Vector and Dsup containing cell lines have equal apoptotic rates.

### Cell Adhesion

Assessment of cell adhesive properties was done using Cell BioLabs CytoSelect™ Cell Adhesion Assay kit. First, cells were spun down and resuspended at 1x10^5^ cells/ml in serum free media.

150μl of cell suspension was added in triplicates to each of the well in an ECM adhesion plate included in the kit. The ECM adhesion plate contains rows of wells coated with either Fibronectin, Collagen I, Collagen IV, Laminin I, Fibrogen, and BSA as a negative control. One set of plates was set aside to be incubated at standard culturing conditions at 37°C and 5% CO_2_ and left unirradiated as a second set of matched plates was irradiated at 2Gy and then incubated along with the 0Gy control plates for 1hr. Media was then aspirated and wells were washed gently 5 times with 1X PBS. 200μl of 1X lysis Buffer with 1:300 diluted CyQuant® GR dye solution was added to each well and then incubated at room temperature for 20 mins while shaking. 150μl lysed solution was then transferred over to a 96 Well White Polystyrene microplate and fluorescence was measured at 480nm_Ex_ /520nm_Em_ emission on a Promega™ Glowmax® Plate Reader. Relative fluorescent units (RFU) were plotted and a heteroscedastic two tailed student t-test was performed to test the null hypothesis that the HEK293 control and each of the other compared cell lines have equal RFUs for each of the conditions.

### 8-hydroxy 2 deoxoguanosine

Cells were seeded in complete DMEM media in 100mm dishes at a density of 2.2 x 10^6^ cells/well in triplicates for each respective cell line and treatment condition. Each cell type was harvested 1hr and 24 hrs post treatment and DNA was extracted using Qiagen MagAttract^®^ HMW DNA kit. After extraction DNA was quantified using Qubit™ 1x dsDNA HS Assay kit and samples were normalized to 10μg of total DNA. DNA was then digested using 10units of NEB nuclease P1 with 10X nuclease p1 extraction buffer and incubated at 37°C for 30mins followed by inactivation at 75°C for 10mins. Digested DNA was then de-phosphorylated with 1unit of NEB shrimp alkaline phosphatase with 10X Cutsmart buffer and incubated at 37°C for 30 mins followed by a 65°C heat inactivation step for 5 mins.

Meanwhile, reagents were prepared for Abcam 8-hydroxy 2 deoxoguanosine ELISA kit. 1X 8-hydroxy-2-deoxoguanosine HRP conjugate monoclonal antibody was diluted at a 1:100 concentration using supplied antibody diluent. Standards were then prepared in serial dilutions ranging from 60ng/ml to 0.94ng/ml using supplied stock 8-hydroxy 2 deoxoguanosine standard with 0ng/ml being diluent. 50ul of samples and standards were added in triplicate to 8-hydroxy 2 deoxoguanosine BSA coated plate followed by the addition of 50ul of antibody. 50ul of sample diluent were added to blank wells. The plate was then covered and incubated for 1hr at room temperature before plate contents were aspirated and wells were washed 4 times with 1X wash buffer using a multichannel pipette. 100ul of supplied TMB substrate was then added into each well incubated at room temperature in the dark for 30mins. The enzymatic color change was then stopped using 100ul of provided stop solution and absorbance at 450nm was immediately measured using a Promega™ Glowmax® Plate Reader. Standard curves were generated and nM was calculated accordingly from absorbance reads. Significance was calculated via paired students t-test comparisons of all cell types to control HEK293 cells, ns=p>0.05, *=p<0.05, **=p<0.01, ***=p<0.001.

### RNA Library Prep and Sequencing

HEK293 Dsup, HEK293 empty vector, and WT HEK293 cells were seeded onto 6 well round bottom plates in triplicates for each cell type, treatment, and time point at a density of 3 x 10^6^ cells/well. Cells were irradiated the following day post seeding at 0.5Gy, 1Gy, 2Gy, and 4Gy of X-rays while 0gy of no treatment controls were kept outside the incubator for the duration of radiation treatment. The cells were then incubated at standard cell culture conditions and harvested at 1hr, 6hrs, and 24hrs post treatment. A total of 93 samples were selected for NEBNext® Ultra™ II RNA library prep consisting of 3 HEK293 DSUP replicates x 3 time points x 4 radiation doses + 3 0gy Controls. The same was done for HEK293 cells and 3 HEK293 empty vectors at 2Gy and 6hrs were included. Trizol extractions were performed as and RNA was isolated carefully from the upper aqueous layer and pelleted before being eluted in 30ul of RNAse free water. RNA concentration was measured using Qubit™ HS RNA assay and RIN and fragment distribution was obtained using Agilent High Sensitivity RNA ScreenTape System for TapeStation 2200. RIN was above 9 for all samples and 250ng of RNA sample was brought forward for rRNA ribosomal depletion using NEBNext^®^ rRNA Depletion Kit for human/mouse/rat samples. Following depletion, RNA samples were library prepped and adapter ligated with single Index Primer set 1 NEBNext^®^ Multiplex Oligos for Illumina^®^. Libraries were then sequenced at 40million single end 100bp reads on Novaseq 6000. Library preps for HEK293 Dsup, HEK293, HEK293 HMGN1-OE, and HEK293 Dsup HMGN1 KO were also performed using the same protocol but with paired end 100x100 reads and for 0 and 2Gys of radiation 1hr after treatment and including ERCC RNA spike in control.

### RNAseq Analysis

GTF and FASTA files for the human reference genome (GRCh38) were downloaded from ENSEMBL. Both GTF and FASTA files were edited to contain the coding sequence for Dsup. After a reference index was created using STAR (ver 2.7)^76^, the reads were then run through the nf-core/rnaseq pipeline. Briefly, read quality metrics were generated using FastQC and TrimGalore was used to remove adapters and low quality regions. Reads were then aligned to the reference genome using STAR and BAM files were generated. RSeQC was used to further analyze the quality of the reads and plots were generated in a MultiQC report that includes BAM stat, junction saturation, RPKM saturation, read duplication, inner distance for paired end reads, read distribution over genomic features, and junction annotations. Qualimap was used to calculate assignments of read alignments, transcript coverage, genomic origin for reads, junction analysis, and 3’-5’ bias. Duplication rates against expression RPKM was plotted using dupRadar and Preseq was then used to estimate the complexity of the library. Afterwards, the read distribution over genomic features was summarized using featureCounts from the subread package and the output .txt file was loaded into R Studio to generate plots, assess differential expression, and correct for batch effects. Reads that did not align to the reference genome were run through metagenomic k-mer based alignment and classification via Kraken and was included in the design matrix as a mycoplasm covariate for limma batch effect removal. A design matrix was generated for different comparisons and covariants that included genotype, radiation dose, and time. For the dataset comparing Dsup, HMGN1 KO, and HMGN1 OE cell types only genotype and dose was included in the design matrix and batch correction was not necessary.

Unsupervised clustering was performed using the log of TPM and plotting dendrograms, PCA, and tSNE plots for each coefficient in the design matrix. Differential expression gene analysis was performed using limma voom and DESeq2^79^. For limma voom the mean variance trend was estimated non parametrically using log CPM and a precision weight was generated for all individual observations^87^. In DESeq2, empirical Bayes shrinkage was used to estimate dispersion parameters within groups to assess for variability between replicates and for shrinkage of fold change estimation to then rank genes. To test for differential expression, DESeq2 used a Wald test to return a z-statistic and p-values from a subset of genes that were adjusted using multiple testing via Benjamini and Hochberg correction^79^. Volcano plots were generated for the resulting DEG with a q value cutoff of 0.01 and Log_2_FC of 1 and top 15 downregulated and upregulated genes were labeled. Fgsea was then used to obtain preranked expression to identify classes of overrepresented genes^64^. Altuna plots, venn diagrams, and heatmaps were also plotted using the same q value and Log_2_FC parameters. Predefined gene sets from MsigDB were grouped together by their involvement in the same biological pathway or by location on chromosome^80^. Features were ranked by differential expression analysis adjusted p value with corresponding Log_2_FC to get normalized enrichment scores^81^. Common genes among different time points for 0Gys of radiation were obtained as the intersection of all genes across the three timepoints and Enrichr was used to identify enriched gene sets using a hypergeometric model^74, 75^. KEGG pathways were also generated from the DEG dataset with a p-value cutoff of 0.05 and Log_2_FC cutoff of 1.5 and visualized on the web based integrative data mining system WebGestalt^82, 83^

### Omni-ATAC Library Preparation and Sequencing

HEK293 Dsup and WT cells were seeding in 12 well plates and incubated at standard conditions. The following day cells were irradiated at 2Gy and unirradiated controls were set aside. Cells were harvested, counted, and assessed for > 90% viability as mentioned above. 50,000 cells were used for Omni-ATAC library preparation as outlined by Corces et al. Briefly, cells were pelleted and resuspended in ATAC-Resuspension Buffer, incubated for 3 mins on ice, washed with ATAC-RSB containing no digitonin and pelleted again. The buffer was aspirated and the cells were resuspended in 50 µl of transposition mixture and incubated at 37°C for 30mins shaking at 1000 RPM. The reaction was cleaned up with a Zymo DNA Clean and Concentrator-5 kit and eluted in 21 µl of elution buffer before being amplified for 5 PCR cycles using NEBnext 2x MasterMix and NEBnext Multiplex Oligos. Additional cycles were determined by qPCR with 5 µl of the amplified sample. The final amplified products were cleaned up once with Ampure XP beads. The purified library was quantified and then fragment size distribution was assessed on a Fragment Analyzer before being sequenced on a Novaseq S4 flow cell at 50x50 PE reads for a read depth of 50 million.

### ATACseq Analysis

Fastq files were run through the nf-core/atacseq pipeline v1.0dev. First raw reads were quality controlled using FastQC. Adapter trimming was then performed with Trim Galore! Trimmed reads were then mapped to the reference genome using BWA and Samtools was used to generate indexes from the output BAM files. Libraries were merged and duplicates marked with picard and mitochondrial reads removed. Normalized bigwig files to 1 million mapped reads were then generated from the BAM files and were used subsequently for coverage visualization via IGV and to generate heatmaps and profile plots with deepTools after peaks were called with MACS2 using the –broad parameter to generate resulting bed files. HOMER was used to annotate peaks to genomic features and FRiP scores were generated and integrated into a MultiQC report.

Consensus peak sets were generated using BEDTools and an UpSetR plot was generated to illustrate peak intersection. Read counts relative to the consensus peaksets across all samples were generated using featureCounts and DESeq2 was used to perform differential accessibility analysis from which MA plots were generated. An ataqv .html file was generated to assess various QC parameters for comparison across samples. A merged replicate analysis was also generated using the same tools outlined above. Nucleosome positioning and occupancy were analyzed using NucleoATAC using broad peak files called by MACS2 and associated BAM files. Heatmaps showing enrichment around TSS were generated by using deepTools computeMatrix to generate a matrix containing normalized bigWig scores for GencodeV36 TSS bed file and profile plots and heatmaps were then generated using deepTools plotHeatmap and profileplyr. Nucleosome occupancy signals were plotted over CpG Island promoters obtained from GRCh38. Enrichment around promoters and enhancers were also evaluated as described above using FANTOM bed files. A design file for each of the time points and dose of radiation was created in R Studio from which count and annotation data generated by featureCounts and HOMER was loaded in order to perform unsupervised clustering analysis. Peaks were then annotated using annotatr for Hg38 CpGs, enhancers, promoters, gene bodies ranging from 1kb to 5kb, 5’UTRs, gene cds, introns, and 3’UTRs. Different peak annotation were plotted by either count or fraction and were assessed as either belong to stable, closing, or opening peaks for Dsup expressing cells vs WT cells. Peaks were then either annotated to gene promoters 5kb of the nearest TSS, genes annotated to the nearest peak within 5kb, or peaks were assigned to genes using GREAT regulatory domains for promoters and enhancers. Fgsea was then performed with a q value cutoff of 0.05 and NES of 1.2 on these annotated gene sets for pathway enrichment analysis and visualization with heatmaps and lollipop plots. Altuna and venn diagrams were plotted using a q value of 0.01 and a Log_2_FC of 0.58 for the different comparisons of opening and closing peaks. Next, HOMER and a multivariate model for JASPAR was used to call motifs in opening and closing regions with a q value cutoff of 0.05 and Log_2_FC of 0.58.

### Cut&Run Library Preparation and Sequencing

Cut&Run libraries for HEK293 Dsup, WT, HEK293 Dsup HMGN1 KO, and HEK293 HMGN1 OE cells were prepared following Epicypher^®^ Cutana^™^ Cut&Run protocol adapted for a 96 well plate format. All reagents and spin columns were obtained from CUTANA ChIC/CUT&RUN kit (SKU: 14-1048). Cells were harvested and 50,000 cells, washed, and bound with activated ConA beads. After magnetic separation, supernatant was removed and antibody buffer solution was added to each sample along with 1:100 dilutions of Epicypher antibodies for Histone H3K27ac (SKU: 13-0045), Histone H3K4me3 (SKU: 13-0041), Bethyl HA Tag antibody (A190-108A), Cell Signaling Technology Ezh2 XP^®^ (Cat: 5426) and Invitrogen Histone H3K27me3 monoclonal antibody (G.299.10). Cutana Rabbit IgG antibody (SKU: 13-0042) was used as a negative control. Samples were incubated overnight at 4°C rotating followed by magnetic separation and washing with 0.01% digitonin buffer. Cutana pAG-MNase were bound to each sample washed again twice with digitonin buffer and chromatin was digested and released with 100 mM CaCl_2_ and incubated for 2 hrs at 4°C. Stop buffer with 1ng of Cutana^™^ E.coli Spike-in DNA (SKU: 18-1401) was added to each sample followed by an incubation at 37°C for 10 mins. 10ul of purified DNA fragments were used for library prep following the NEBNext Ultra II for DNA protocol but with 16 amplification cycles. Samples were loaded on a Novaseq S4 flow cell and sequenced paired end 50x50 at 40 million reads.

### Cut&Run Analysis

Reads were preprocessed for quality control using FastQC followed by trimomatic adapter read trimming and Bowtie2 alignment to our STAR generated reference genome and e.coli ASM484v2 genome assembly for spike-in reference. Paired ends were aligned end to end for mapping of inserts 10-700bp in length. The spike-in controls were aligned and a scaling factor was calculated for each sample. Duplicates were marked using DeDup and samtools was used to sort and index resuting BAM files. Alignments were filtered into two classes of reads, fragments less than 120bp and all fragment sizes. Paired-end bedgraphs were then generated first using bedtools bamtobed and then bedtools genomecov to convert bed files into bedgraphs and normalized to spike-in control using the scaling factor. UCSC bedGraphToBigWig was used to generate bigwig tracks from the final alignments for visualization on IGV. DeepTools bamCoverage was also used to generate normalized bigwig files to reads per genome coverage at 1x depth using the effective genome size for comparison. MACS2 was used to call peaks from normalized bedgraph files using -f BAMPE, qvalue < 0.05, -B --SPRM, and --broad depending on what histone PTM was being analyzed. SEACR was also used for peak calling with both relaxed and stringent settings and with and without normalization to control IgG input bedgraph. The called peaks were then annotated with genomic features using HOMER. A MultiQC report was generated for the individual samples to assess various quality metrics before merging replicate BAM files and performing the above steps to generate bedgraph, bigWig, and called peaks. Matrices around a reference point were created using normalized bigWig files and either intersected peak called bed files or GencodeV36 TSS bed files were used to plot using deepTools plotHeatmap. BigWig tracks were also visualized on IGV. Rgt-THOR was used to call differential peaks with -merge parameter for histones, -pvalue set to 0.01 for broad peaks and 0.05 for narrow peaks, and -deadzones for blacklisted regions obtained from HG38 blacklist V2.

Chipseeker was used for further annotation of differentially expressed peaks and enrichment analysis via KEGG and Fgsea using gene set annotations from msigdb.v7.1.symbols.

Permutation tests for testing overlap of differential peaks for each cell type histone PTM was performed using the ChIPpeakAnno package. Briefly, peaks were randomized and shuffled 1000 times to generate random peak lists from which permutations were conducted testing the null hypothesis that there is no difference in overlap of peaks. A background reference peak pool for H3K27ac, H3K4me3, and H3K27me3 for HEK293 cells were obtained from Gencode bed files for random peak sampling.

## Acknowledgements

MTV, RLC, and PAS were sponsored by DARPA and the Army Research Office under Cooperative Agreement Number W911NF-19-2-0017. The views and conclusions contained in this document are those of the authors and should not be interpreted as representing the official policies, either expressed or implied, of any funding agency including DARPA, the Army Research Office, or the U.S. Government. The U.S. Government is authorized to reproduce and distribute reprints for Government purposes notwithstanding any copyright notation herein.

## Author Contributions

C.W., M.T.V, R.L.C, C.C., P.A.S., and C.E.M. helped with experimental design. C.W with the help of D.N. performed all assays, library prep, and experiments except for caspase-3 apoptosis assay. M.T.V and R.L.C performed caspase-3 apoptosis assay. D.B. and C.M. helped with library prep for RNAseq and oxford nanopore long read sequencing. C.W., C.M., and K.G. performed follow up bioinformatic analysis. C.W. wrote the manuscript. Special thanks to summer and medical students Sonia Iosim^1^ and Sherry Yang^1^ for participating in this project. Special thanks to Rafael Colon^1^ for uploading the data to GEO. C.W., D.N., R.L.C., C.C., E.A., and C.E.M took part in editing the manuscript.

---

**Supplemental Figure S1.**
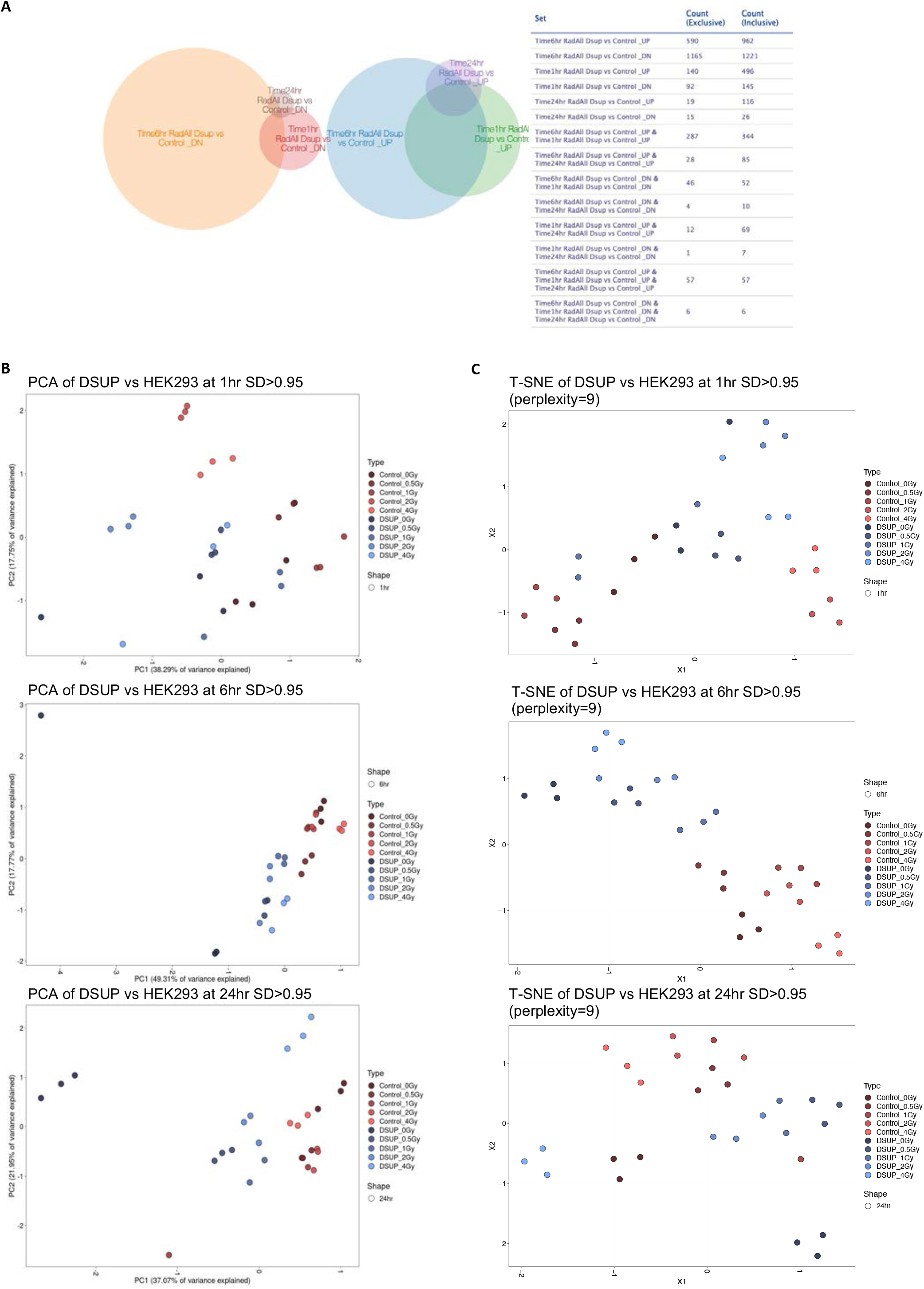
**(A)** Venn diagrams show exclusive and inclusive counts for differentially expressed upregulated and downregulated genes as assessed by Limma-Voom, q<0.01 and the absolute value of Log2FC 1.5. All radiation doses are included in aggregate for each comparison across time. **(B)** Principal Component Analysis Clustering for expression data reveals sample to sample distance as calculated by unsupervised variance stabilizing transformation for each dose of radiation represented by colored circles across time points with PC1 variance on x-axis and PC2 variance on y-axis, SD>0.95. **(C)** T-Distributed Stochastic Neighbor Embedding shows relative distances between the same expression data in a 2 dimensional space, SD>0.95 and perplexity=9.

**Supplemental Figure S2.**
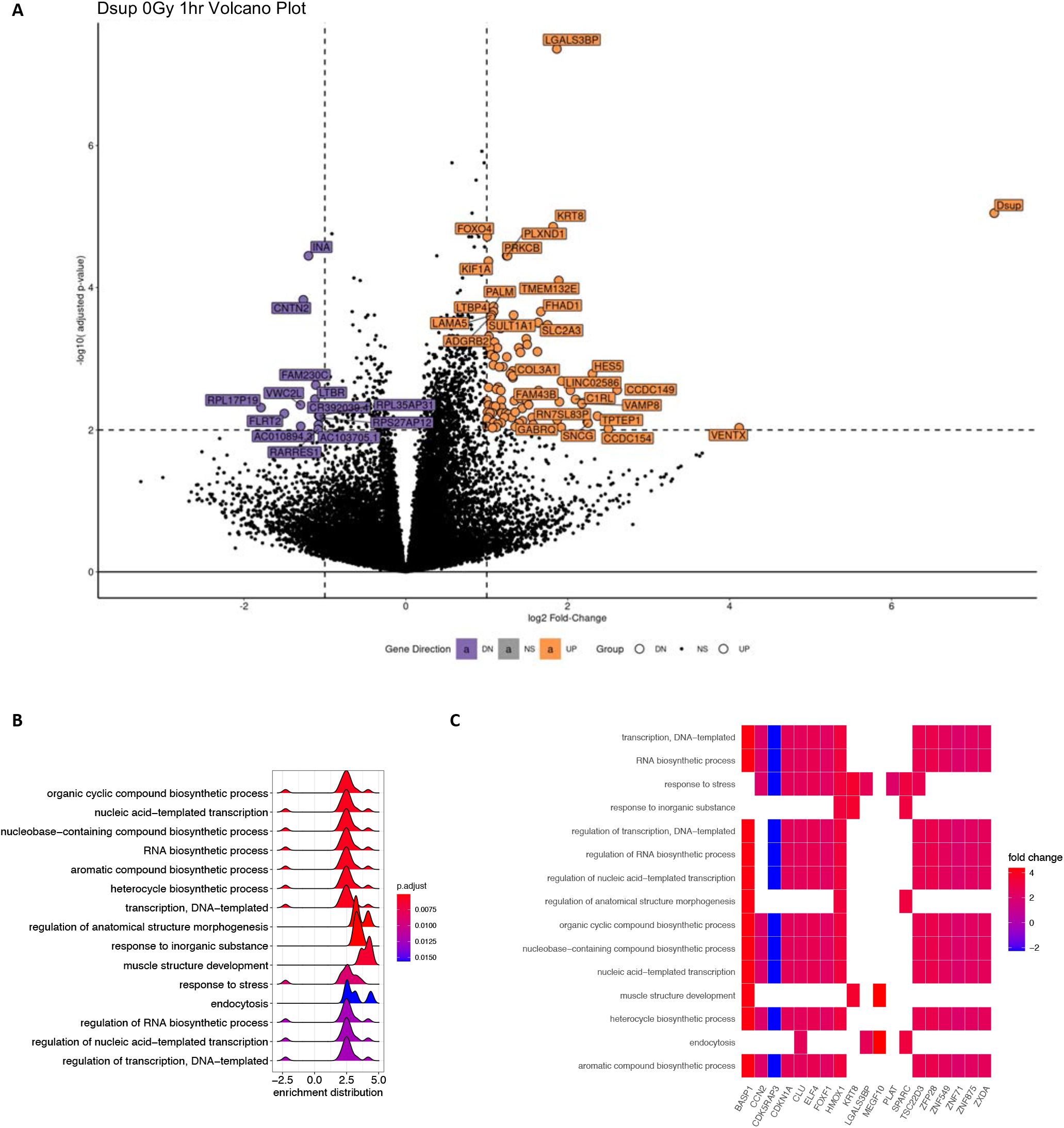
**(A)** Volcano plot depicting top differentially expressed genes for Dsup vs WT at 0Gy and 1hr as determined by Limma-Voom. The –log10 of the p-adjusted value is plotted along the y-axis with horizontal vertical lines representing the -log10 of p-adjusted cutoff <0.01 while Log_2_FC is on the x-axis with dashed vertical lines representing |Log_2_FC| 1. Purple dots represent downregulated differentially expressed genes while orange dots represent upregulated differentially expressed genes. **(B)** Ridge plot shows pathway enrichment analysis for the same set of differentially expressed genes for Dsup vs WT at 0Gy and 1hr using GO pathway terms with p-adjusted value cutoff=0.05 and Bonferroni p-adjusted method. Gradient shows p-adjusted values from red to blue going from smallest to largest with height of peaks representing density of enrichment distribution scores. Top pathways indicate a more transcriptionally permissive state. **(C)** Heatmap depicting the same enriched pathways as before but with associated differentially expressed genes along the x-axis. Color gradient shows fold change from upregulated (red) to downregulated (blue).

**Supplemental Figure S3.**
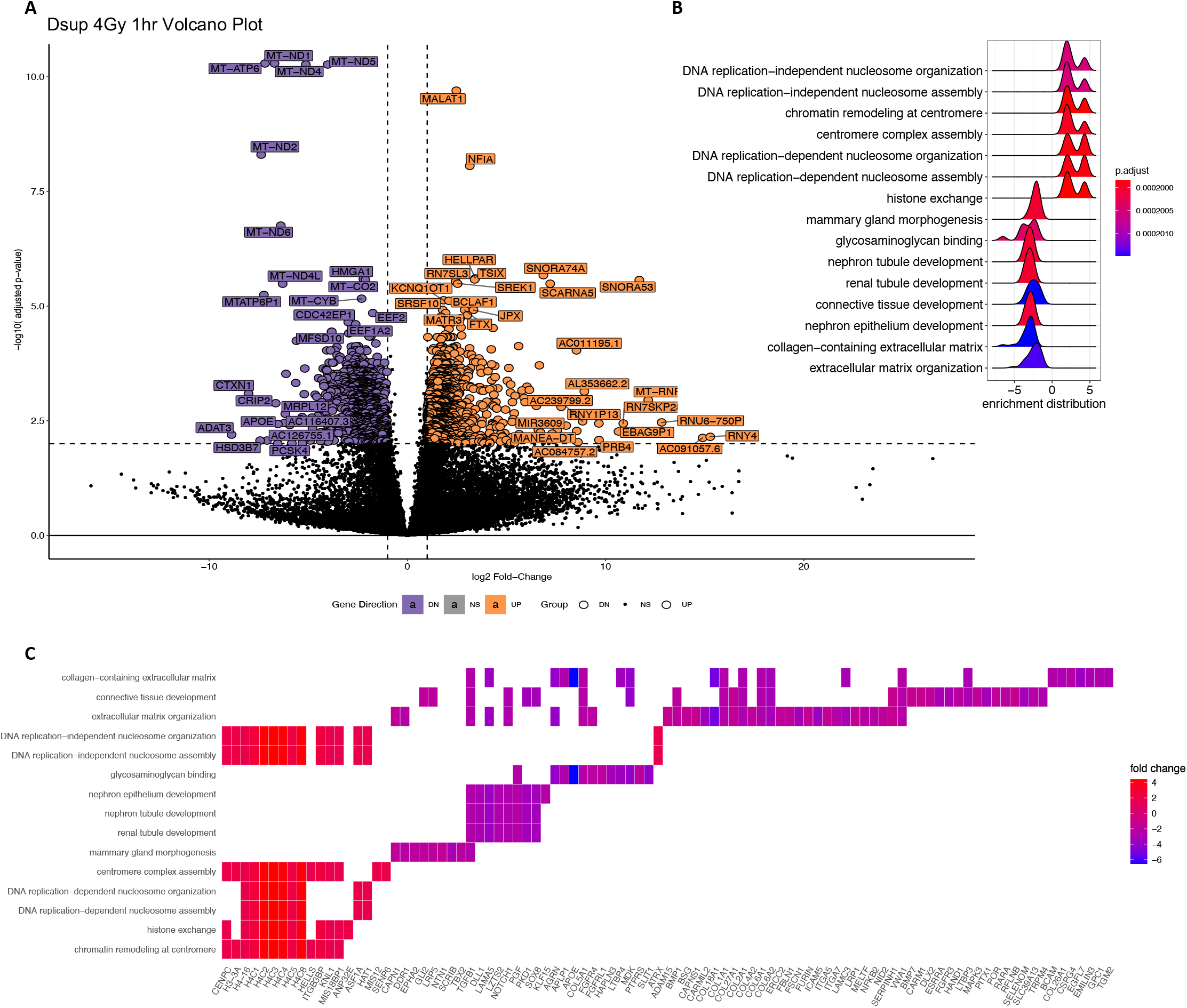
**(A)** Volcano plot depicting top differentially expressed genes for Dsup vs WT at 4Gy and 1hr as determined by Limma-Voom. The –log10 of the p-adjusted value is plotted along the y-axis with horizontal vertical lines representing the -log10 of p-adjusted cutoff <0.01 while Log_2_FC is on the x-axis with dashed vertical lines representing |Log_2_FC| 1. Purple dots represent downregulated differentially expressed genes while orange dots represent upregulated differentially expressed genes. **(B)** Ridge plot shows pathway enrichment analysis for the same set of differentially expressed genes for Dsup vs WT at 4Gy and 1hr using GO pathway terms with p-adjusted value cutoff=0.05 and Bonferroni p-adjusted method. Gradient shows p-adjusted values from red to blue going from smallest to largest with height of peaks representing density of enrichment distribution scores. Top pathways indicate an upregulation of chromatin organization related genes and a downregulation of developmental genes. **(C)** Heatmap depicting the same enriched pathways as before but with associated differentially expressed genes along the x-axis. Color gradient shows fold change from upregulated (red) to downregulated (blue).

**Supplemental Figure S4.**
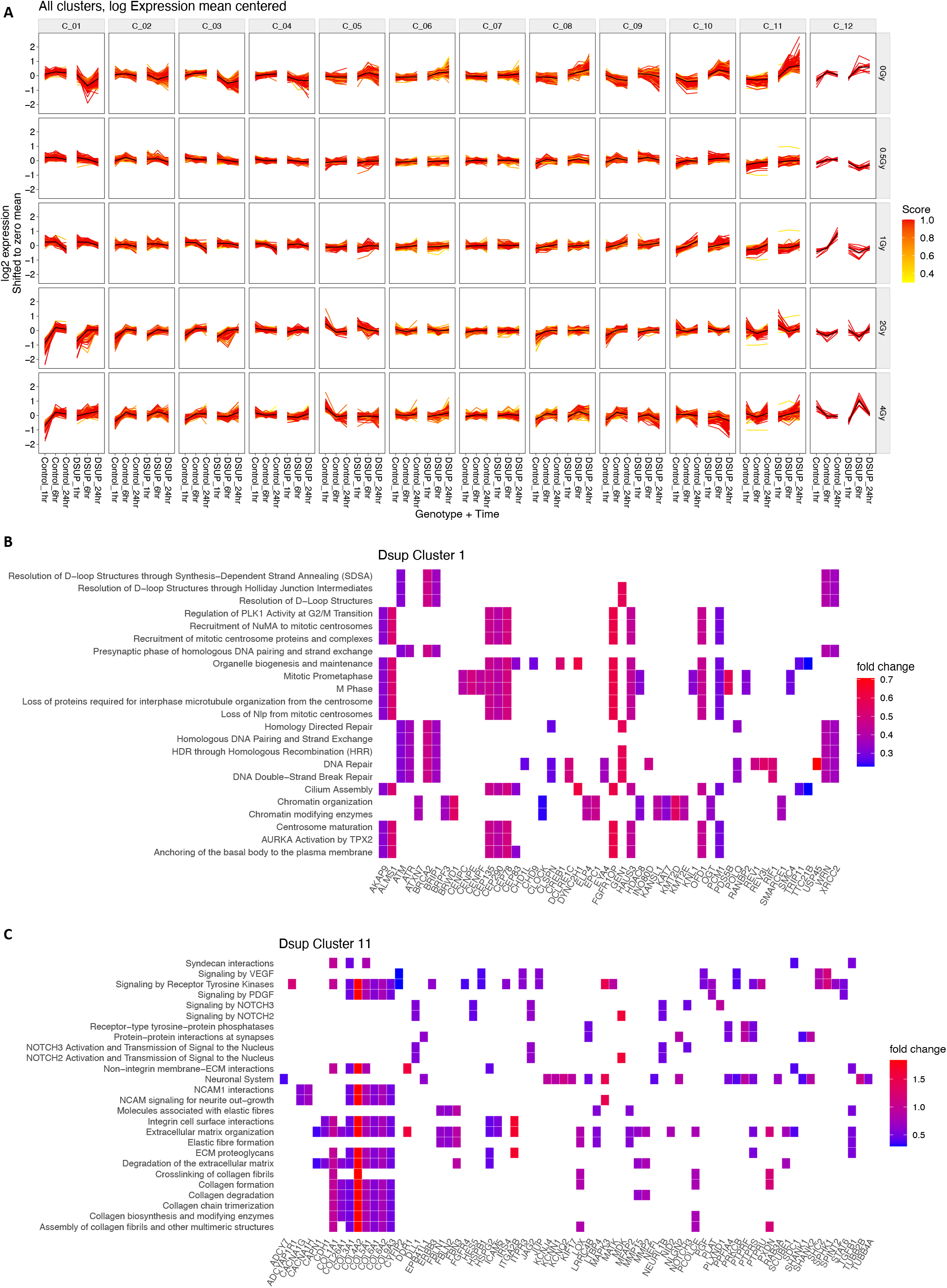
**(A)** Gene trajectory analysis for different clusters of genes show Log_2_FC change dynamics over time when grouped by radiation dose. Log2FC expression scores for all genes were assessed by comparing to baseline mean at 0Gy and 1hr. Genes were filtered first by using q-value < 0.01 and an absolute value Log_2_FC cutoff ≥ 1 and then normalized to baseline mean for purpose of visualization of both HEK293 Dsup and Control HEK293 trajectory dynamics. **(B) (C)** Gene from clusters 1 and 11 were ranked by Log_2_FC and further analyzed using Enricher REACTOME terms and resulting pathways and associated genes were plotted via heatmap with gradient from red to blue showing fold change. Gene sets belonging to cluster 1 were enriched for DNA repair and chromatin organization related pathways while genes belonging to cluster 11 were enriched for NOTCH signaling and collagen synthesis pathways.

**Supplemental Figure S5.**
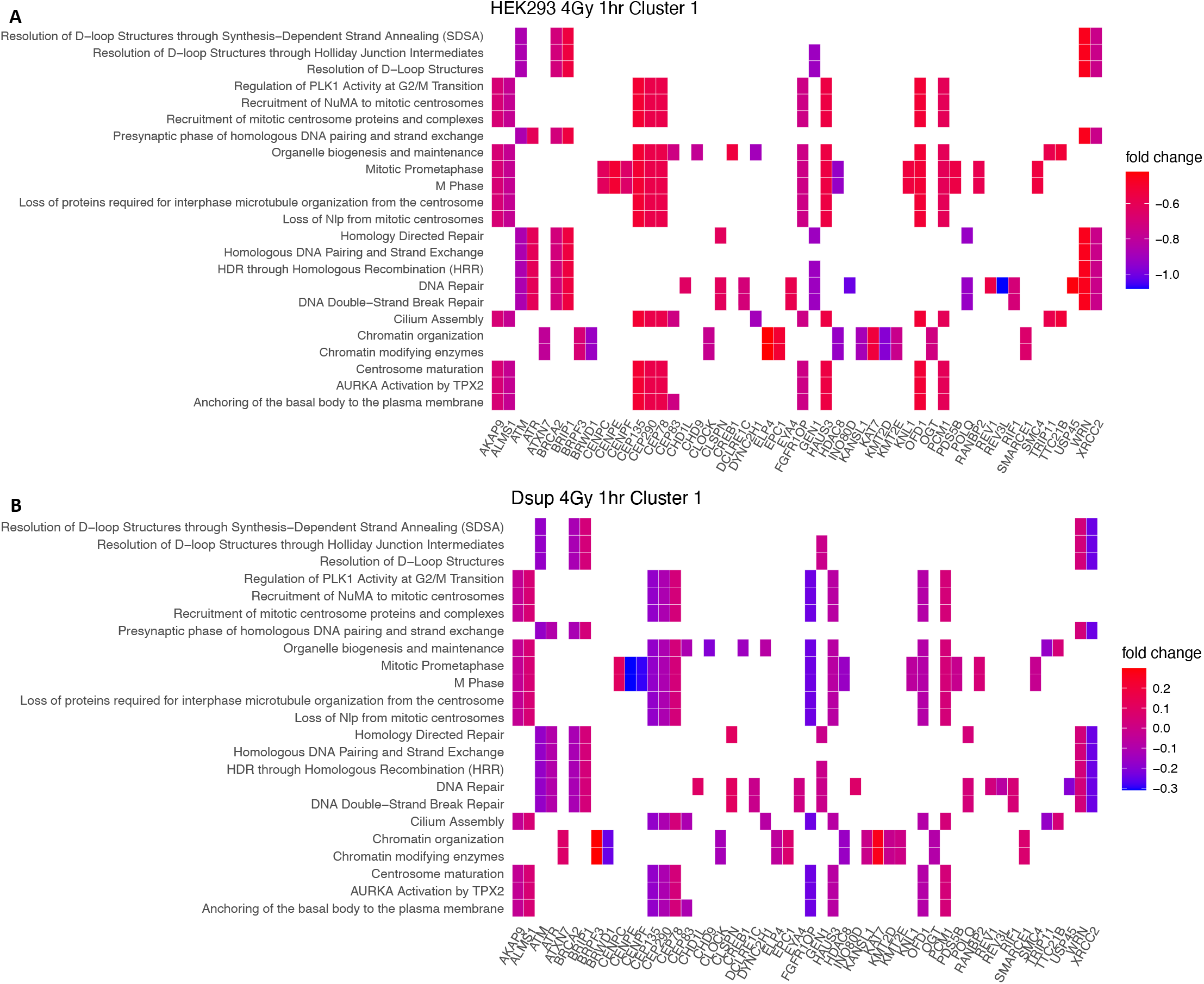
**(A)** Heatmaps show filtered and normalized to baseline Log2FC for genes from cluster 1 at 4Gy and 1hr for HEK293 Control cells and **(B)** HEK293 Dsup cells. Chromatin organization related genes such as BRPF3 remain upregulated Log2FC = 0.29 in HEK293 Dsup while downregulated -0.69 in HEK293 Control. DNA repair genes CHD1L and CLSPN are upregulated in HEK293 Dsup (Log_2_FC = 0.09 & Log_2_FC = 0.11, respectively) and downregulated in HEK293 Control (Log_2_FC = -0.64 & Log_2_FC = -0.62, respectively).

**Supplemental Figure S6.**
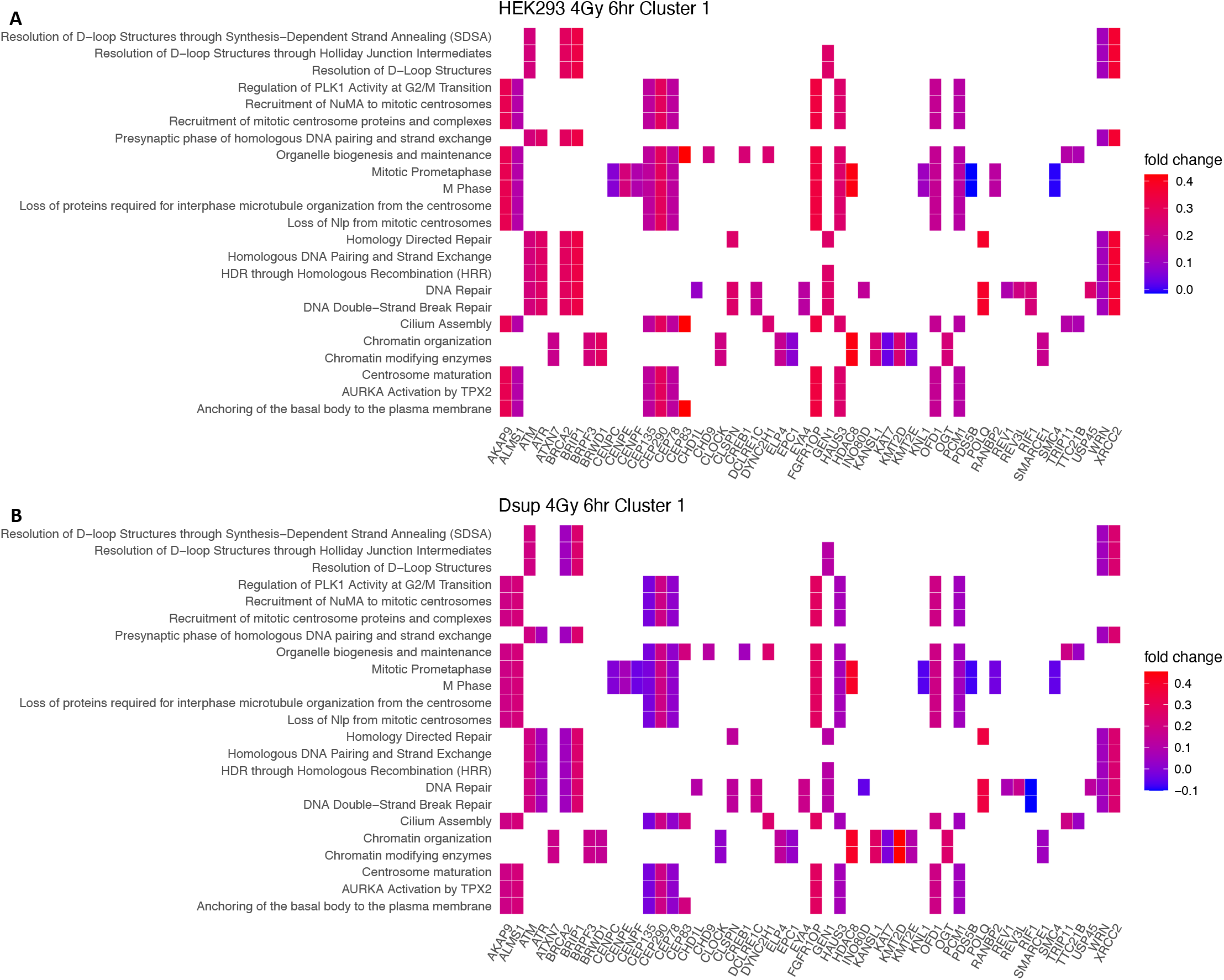
**(A)** Heatmaps show filtered and normalized to baseline Log2FC for genes from cluster 1 at 4Gy and 6hr for HEK293 Control cells and **(B)** HEK293 Dsup cells. DNA repair genes such as RLF1 and INDO80D are now downregulated in HEK293 Dsup (Log_2_FC = -0.10 & Log_2_FC = -0.05) while upregulated in Control HEK293 (Log_2_FC = 0.24 & Log_2_FC = 0.18).

**Supplemental Figure S7.**
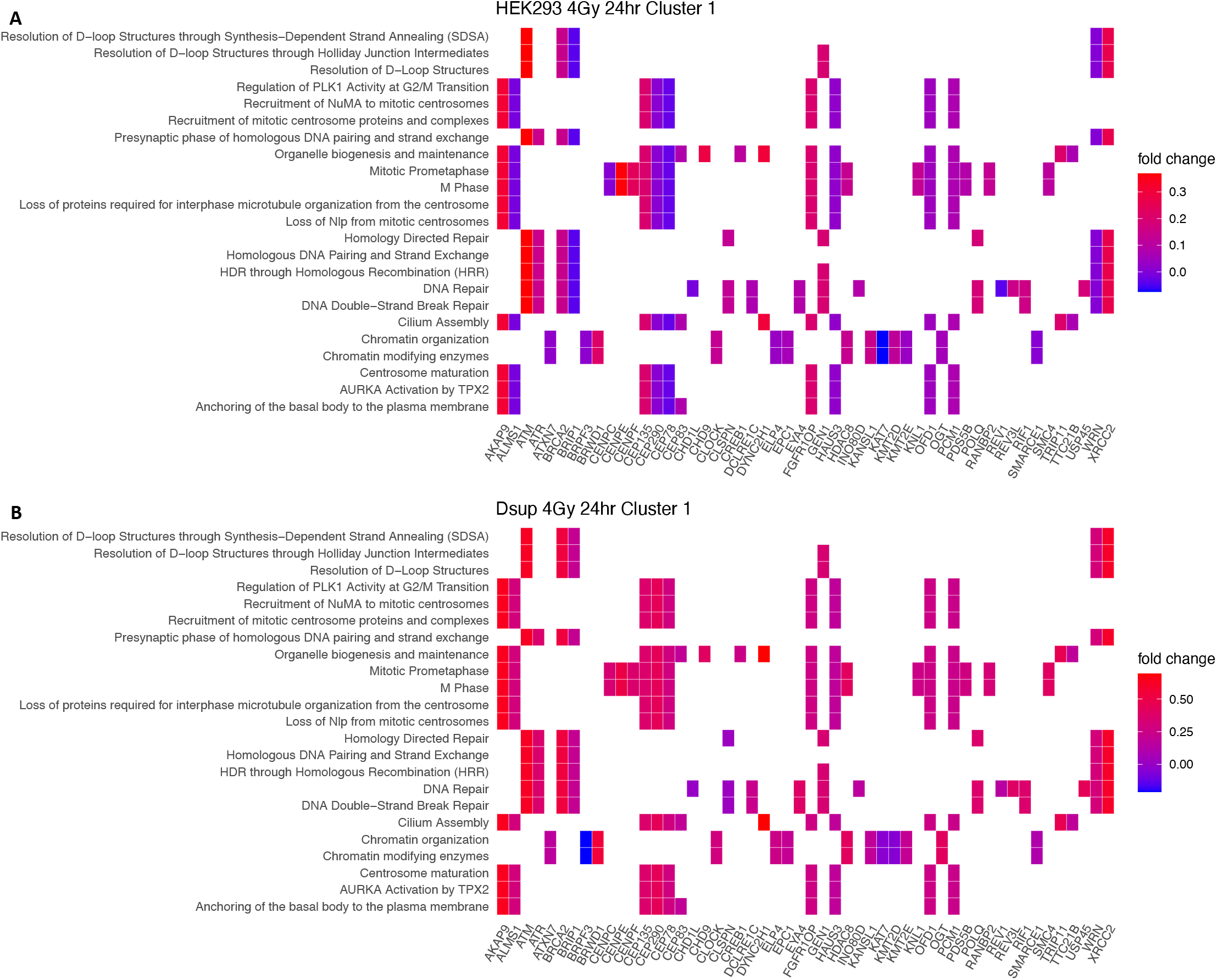
**(A)** Heatmaps show filtered and normalized to baseline Log2FC for genes from cluster 1 at 4Gy and 14hr for HEK293 Control cells and **(B)** HEK293 Dsup cells. Chromatin organization genes such as BRPF3 are now downregulated in HEK293 Dsup (Log_2_FC = -0.21) while upregulated in Control HEK293 (Log_2_FC = 0.02). Mitotic prometaphase and organelle biogenesis related gene CEP78 is upregulated in HEK293 Dsup (Log_2_FC = 0.30) and downregulated in HEK293 Control cells (Log_2_FC = -0.03).

**Supplemental Figure S8.**
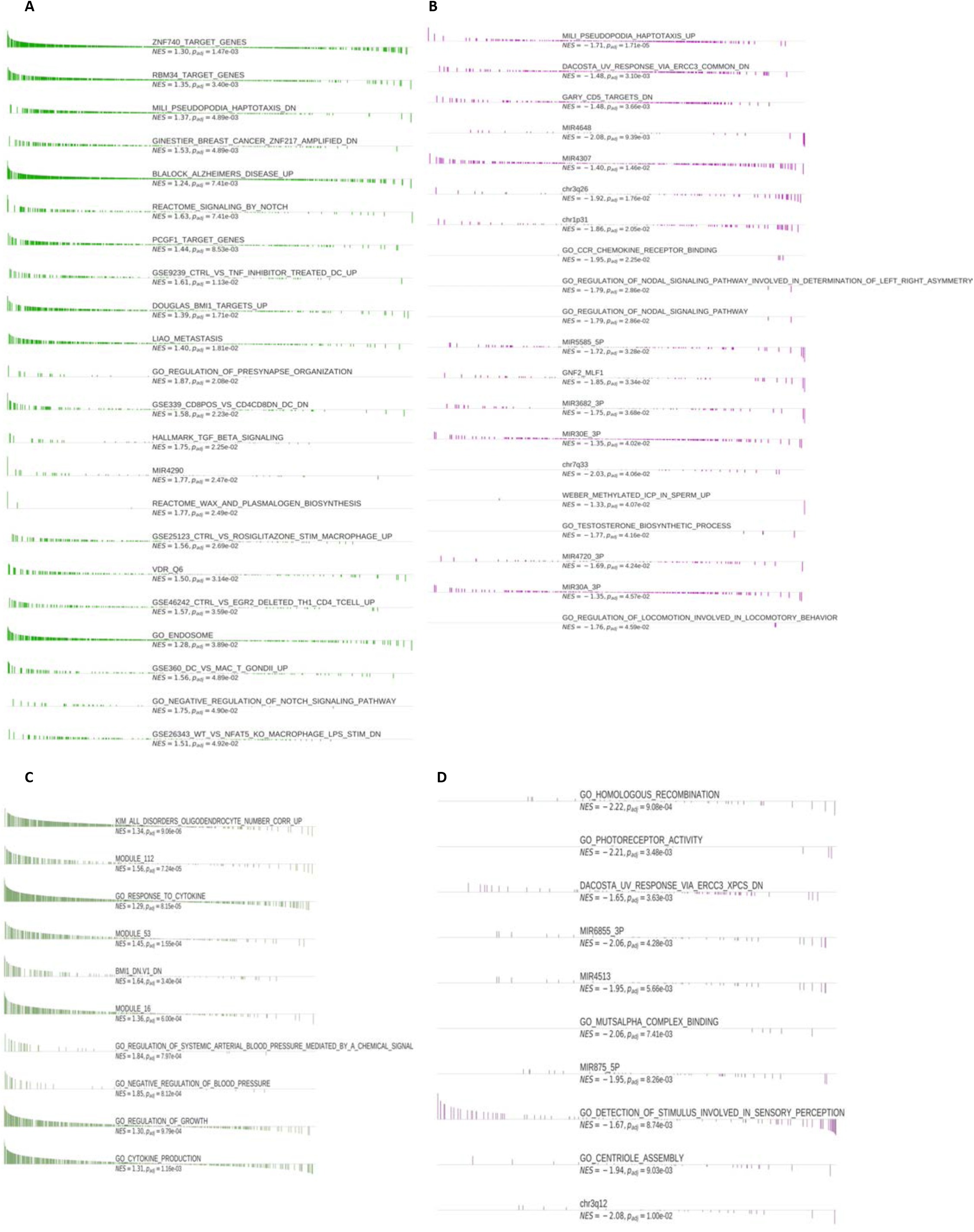
**(A)** Enrichment plot for Fgsea results shows top upregulated pathways for HEK293 Dsup vs Control at 1hr and the aggregate of all doses of radiation combined to see overall effect of Dsup introduction into the human genome. Leading edge genes are ranked and plotted along the x-axis for each associated pathway with normalized enrichment score and p-adjusted value. **(B)** Top downregulated pathways at 1hr. **(C)** Top Upregulated pathways at 6hrs. **(D)** Top downregulated pathways at 6hrs. Differential expressed genes giving rise to all pathways seen here were calculated using Limma-voom with q-value < 0.01 and absolute value of NES > 1.2.

**Supplemental Figure S9.**
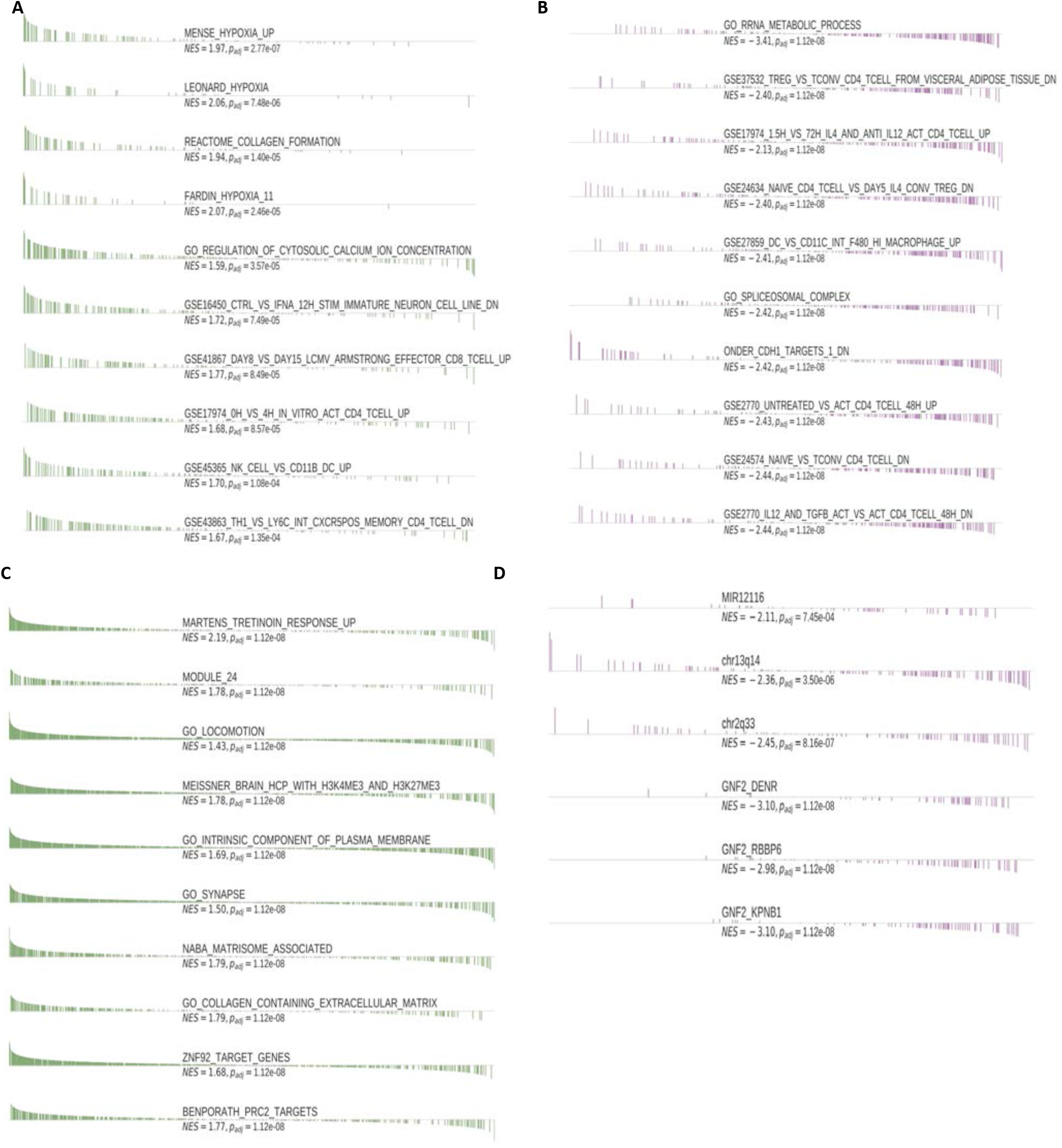
**(A)** Enrichment plot for Fgsea results shows top upregulated pathways for HEK293 Dsup vs Control at 24hr and the aggregate of all doses of radiation combined to see overall effect of Dsup introduction into the human genome at each time point. Leading edge genes are ranked and plotted along the x-axis for each associated pathway with normalized enrichment score and p-adjusted value. **(B)** Top downregulated pathways at 24hr. **(C)** Top Upregulated pathways for HEK293 Dsup with all doses and time points combined. **(D)** Top downregulated pathways with all doses and time points combined. Differential expressed genes giving rise to all pathways seen here were calculated using Limma-voom with q-value < 0.01 and absolute value of NES > 1.2.

**Supplemental Figure S10.**
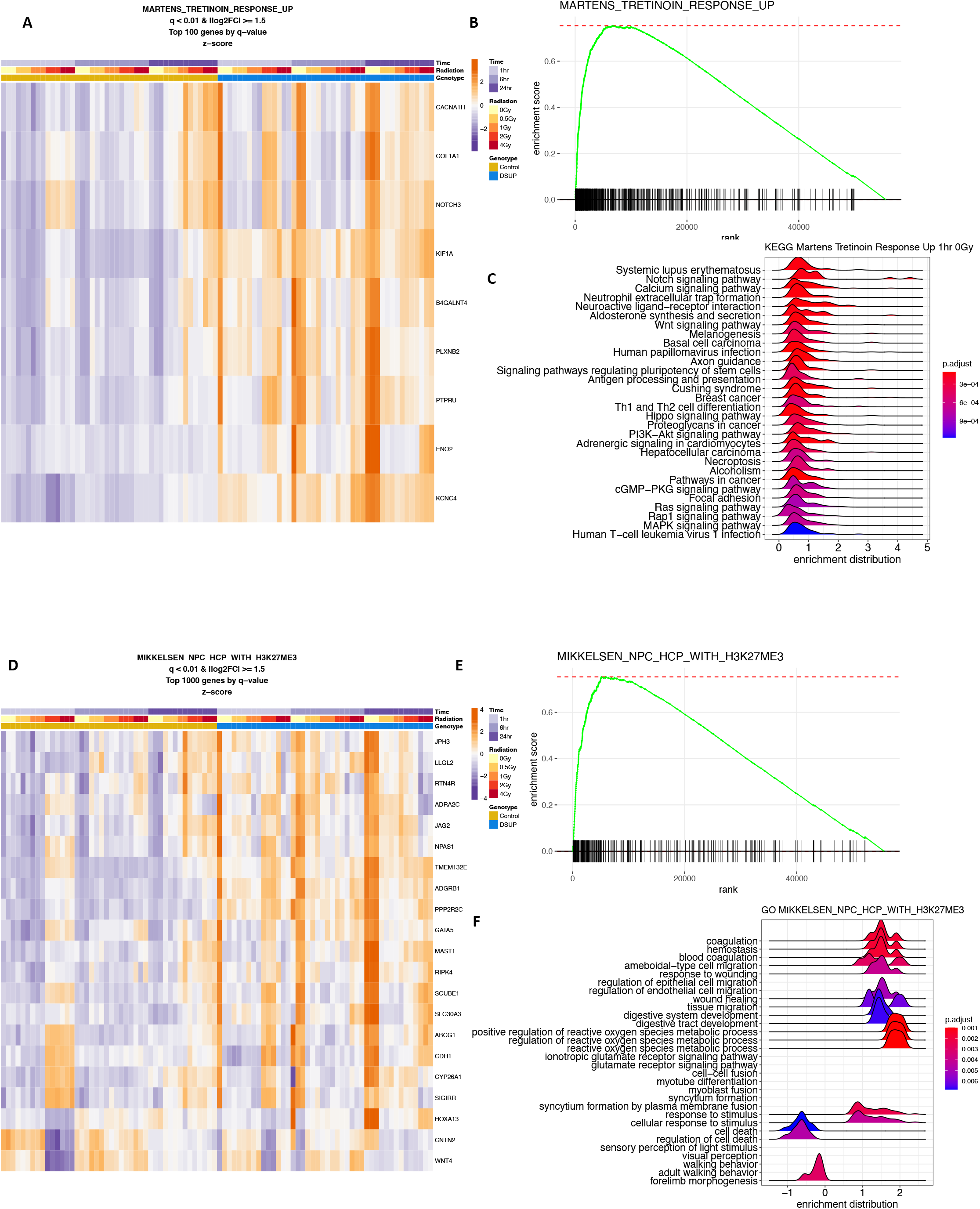
**(A)** Heatmap shows top differentially expressed genes belonging to MARTENS_TRETINOIN_RESPONSE_UP filtered by q-value < 0.01 and absolute value of Log_2_FC 1.5. **(B)** Enrichment plot shows overall upregulation of pre-ranked genes with enrichment score plotted on the y-axis and leading-edge gene rank on the x-axis. **(C)** Ridge plot shows KEGG analysis of the same set of genes with p-adjusted <0.05 with Bonferroni correction. Of note are upregulated pathways enriched for Wnt and Hippo signaling. **(D)** Heatmap shows top differentially expressed genes belonging to MIKKELSEN_NCP_HCP_WITH_H3K27me3 filtered by q-value < 0.01 and absolute value of Log_2_FC 1.5. **(E)** Enrichment plot shows overall upregulation of pre-ranked genes with enrichment score plotted on the y-axis and leading-edge gene rank on the x-axis. **(F)** Ridge plot shows KEGG analysis of the same set of genes with p-adjusted <0.05 with Bonferroni correction. Of note are upregulated pathways enriched for reactive oxygen species metabolism and a downregulation of cell death related pathways.

**Supplemental Figure S11.**
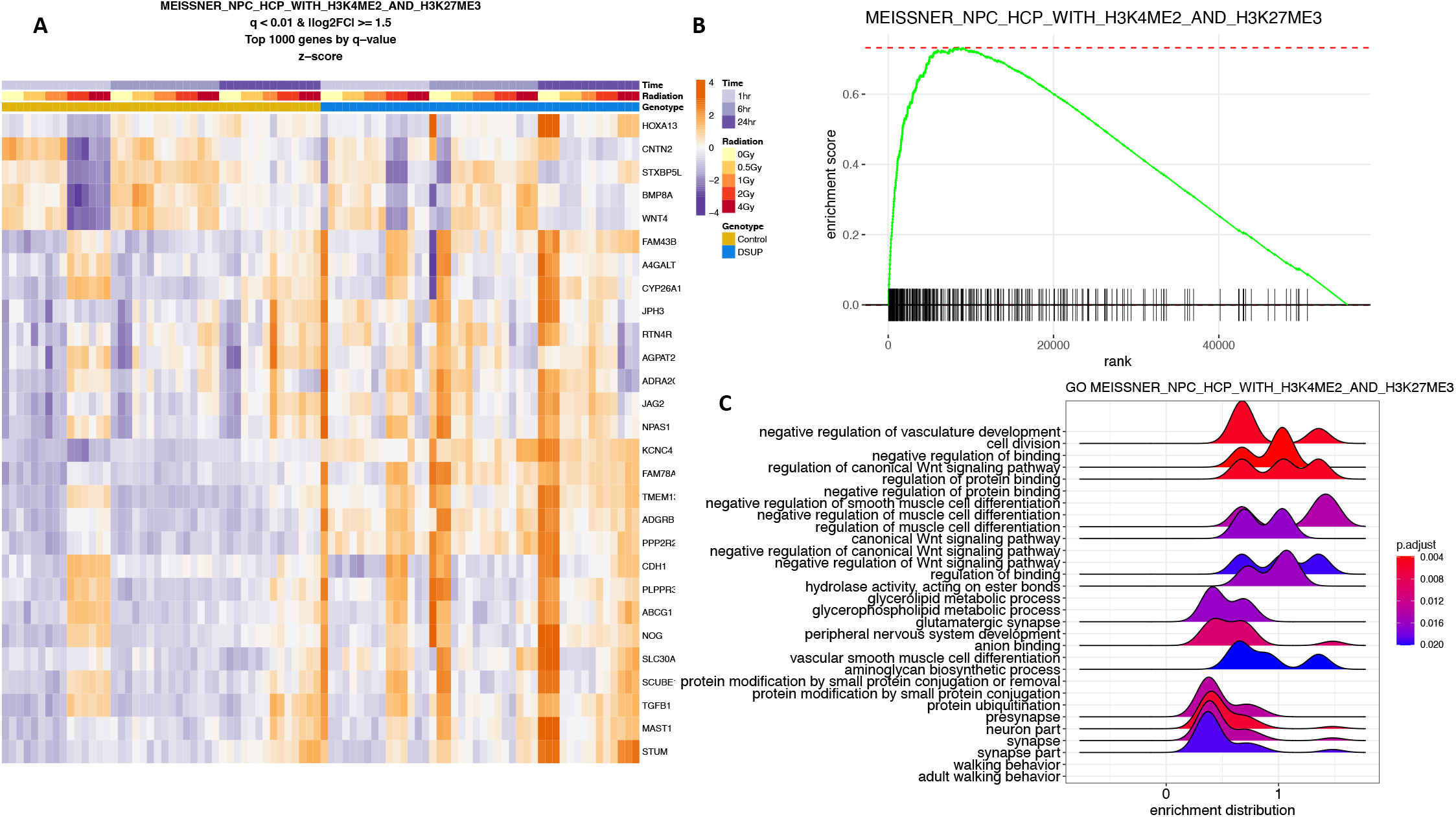
**(A)** Heatmap shows top differentially expressed genes belonging to MEISSNER_NCP_HCP_WITH_H3K4me2_AND_H3K27ME3 filtered by q-value < 0.01 and absolute value of Log_2_FC 1.5. **(B)** Enrichment plot shows overall upregulation of pre-ranked genes with enrichment score plotted on the y-axis and leading-edge gene rank on the x-axis. **(C)** Ridge plot shows KEGG analysis of the same set of genes with p-adjusted <0.05 with Bonferroni correction. Of note are upregulated pathways enriched for Wnt signaling and developmental pathways.

**Supplemental Figure S12.**
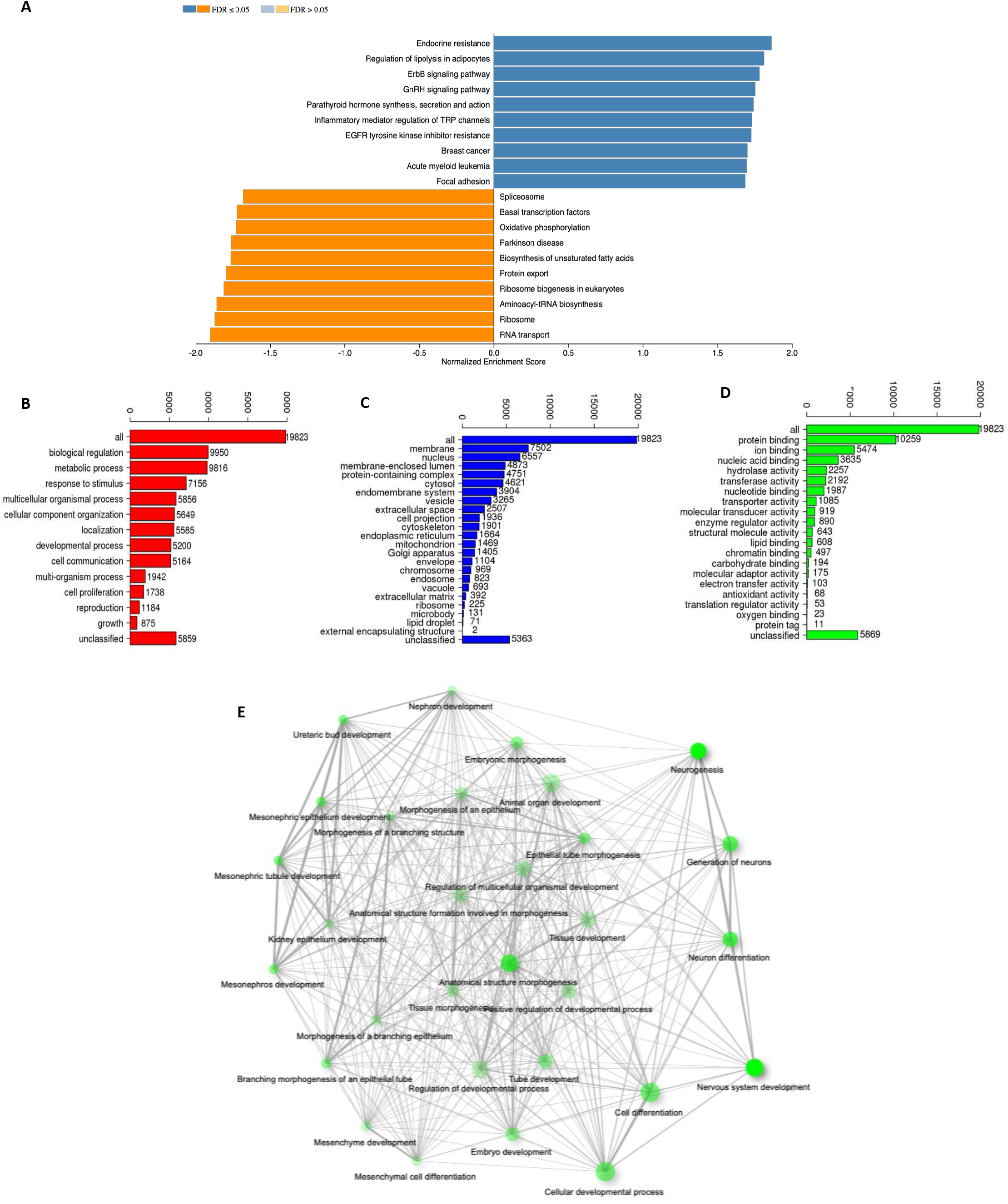
**(A)** WebGestalt results show top KEGG enriched pathways ranked by –sign(Log2FC) x log10(p-adjusted) from Limma-voom DEG results for the interaction of Dsup with all radiation doses and time points. Normalized enrichment score is plotted along the x-axis and significant FDR ≤ 0.05 is displayed with dark blue for upregulation and dark orange for downregulation. **(B) (C) (D)** GO enrichment results for biological process, cellular component, and molecular function from same list of pre-ranked DEG results. **(E)** Shiny GO network visualization of baseline 0Gy and 1hr HEK293 Dsup upregulated DEG displays nodes with developmental processes and shared genes between them using hypergeometric distribution analysis, p-value cutoff < 0.05.

**Supplemental Figure S13.**
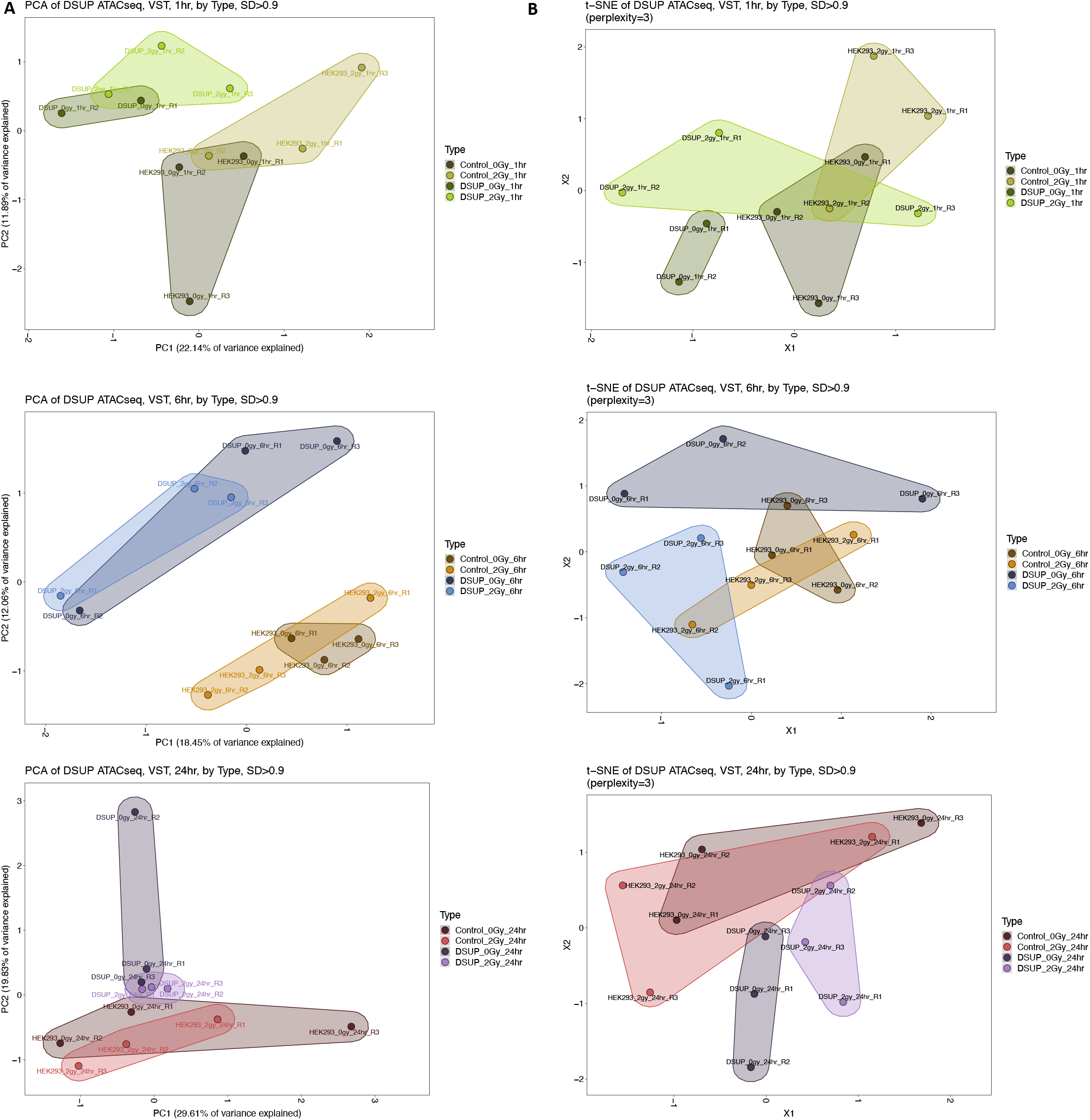
**(A)** Principal Component Analysis Clustering for expression data reveals sample to sample distance as calculated by unsupervised variance stabilizing transformation for each dose of radiation represented by colored circles across time points with PC1 variance on x-axis and PC2 variance on y-axis, SD>0.9. **(B)** T-Distributed Stochastic Neighbor Embedding shows relative distances between the same chromatin accessibility dataset in a 2 dimensional space, SD>0.9 and perplexity=3.

**Supplemental Figure S14.**
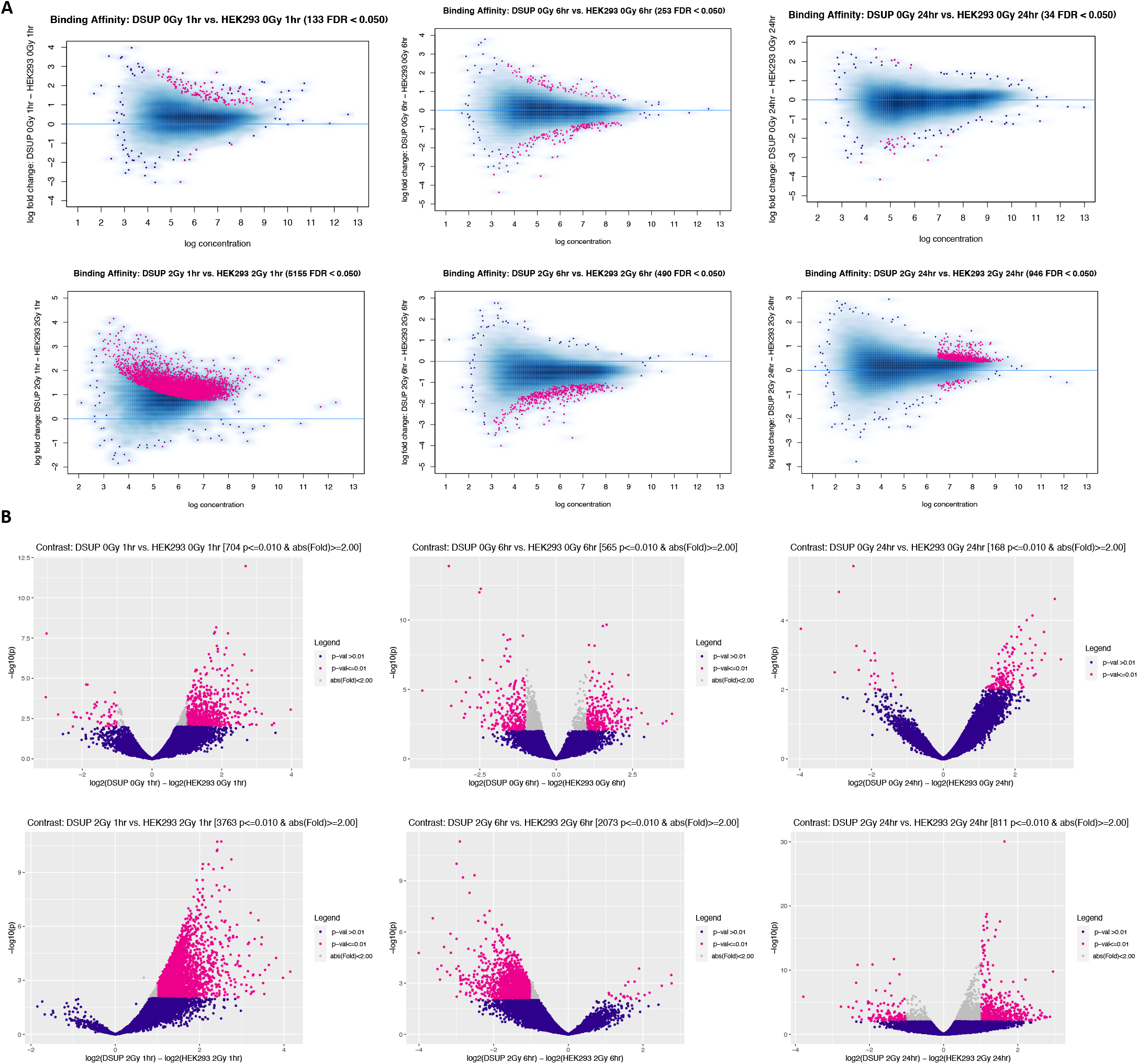
**(A)** MA plots show show differential peak analysis results conducted using DESeq2 and edgeR for each dose and time point for HEK293 Dsup vs. Control HEK293 chromatin accessibility, statistically significant differences are shown in pink FDR<0.05. **(B)** Volcano plots show the same differential peak analysis results but with –log10 p-value plotted on the y-axis and Log_2_FC on x-axis with statically significant results plotted as pink dots, absolute value Log_2_FC 2 & p-value < 0.01.

**Supplemental Figure S15.**
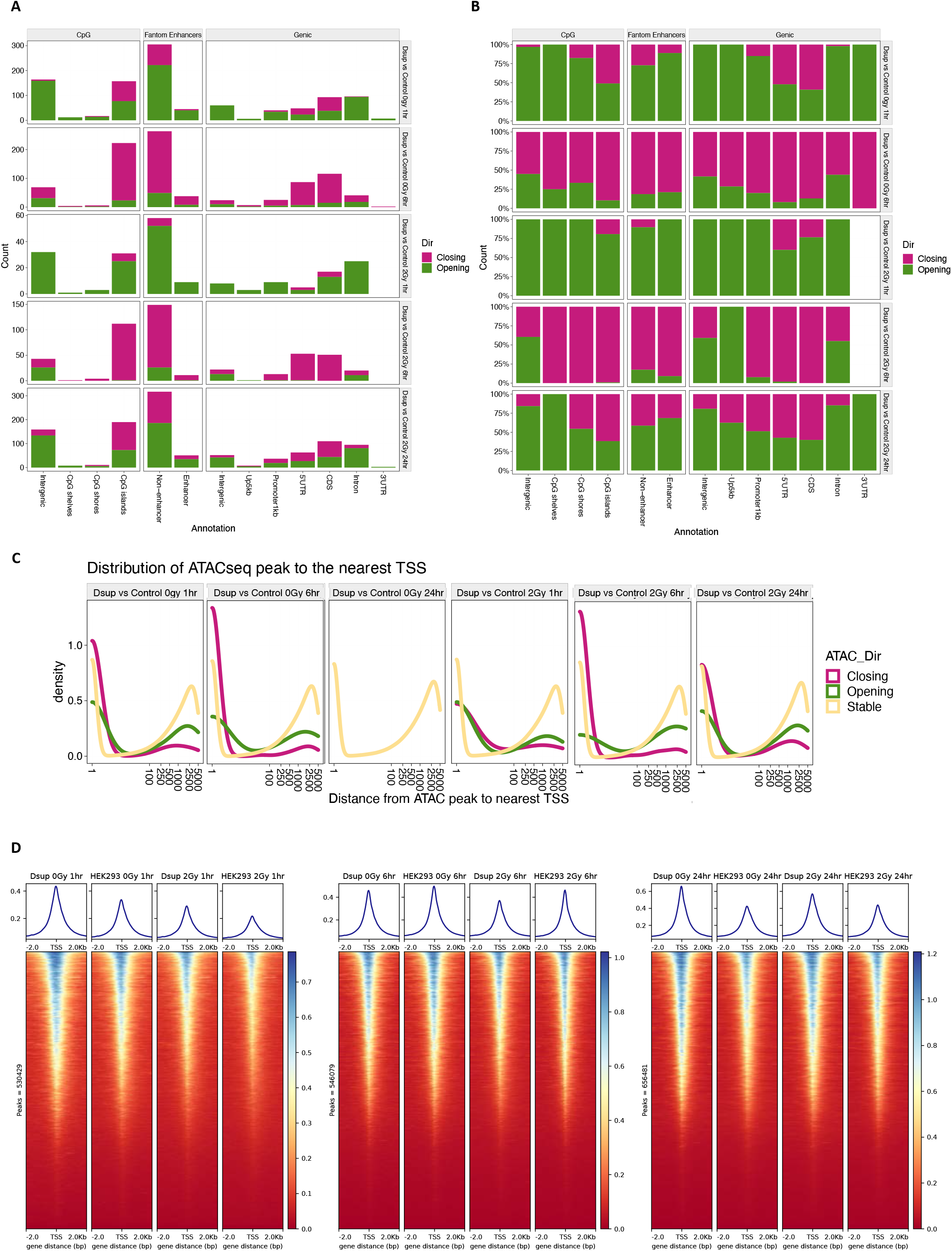
**(A)** Peak count distributions for each dose and time point for HEK293 Dsup vs Control HEK293 control were annotated via CpG, enhancer, and genic regions using hg38 ENSEMBL BioMart annotations after filtering differential peaks with Log_2_FC ≥ 0.58 and q-value < 0.01 to obtain just opening and closing annotated peaks. Peak counts are plotted along the y-axis for each condition. **(B)** The same peak annotation plot but separated by fraction rather than count. Overall Dsup at 0Gy and 2Gys for 1hr has the greater fraction of opening peaks in most areas with 5’ UTRs and coding sequences becoming more open post-irradiation. While 6hrs has the greater fraction of closing peaks. 24hrs at 0Gy is not represented as peaks were overall stable as assessed by differential peak analysis. **(C)** ATACseq peaks were annotated to the nearest gene TSS within 5kb for each dose and time point and relative densities of opening, closing, or stable peaks were plotted along this distance, Log_2_FC ≥ 0.58 and q-value < 0.01. **(D)** Heatmaps were generated for HEK293 Dsup and HEK293 WT Control for each dose and time point by plotting signal over the 2kb of TSS of all genes belonging to GRCh38. Of note, HEK293 Dsup has greater opening of peaks at 0Gy and 1hr compared to HEK293 WT, while signal drops at 2Gy but remains more open for HEK293 Dsup compared to WT.

**Supplemental Figure S16.**
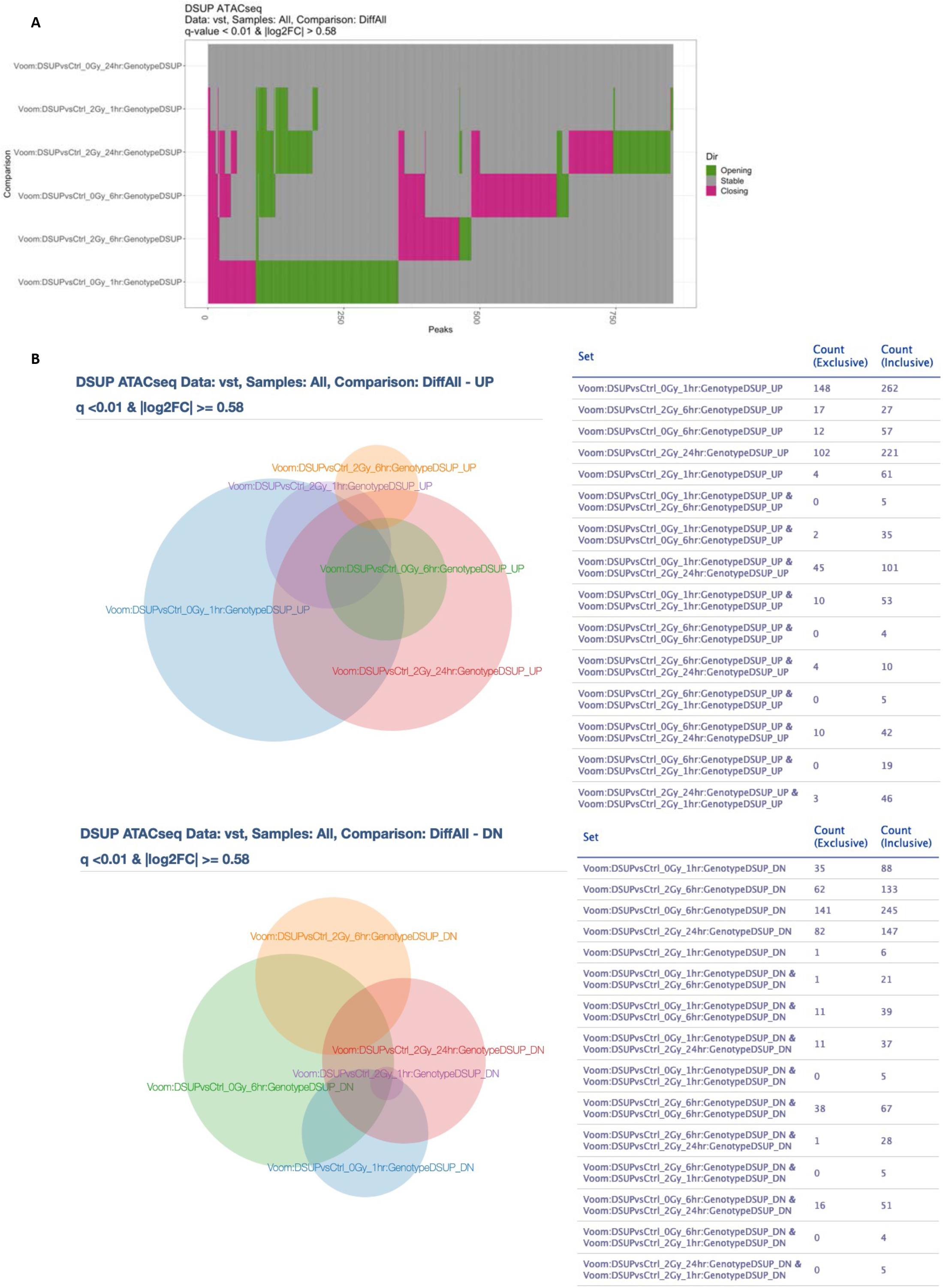
**(A)** Altuna plot reveals comparisons of differentially called peaks for each cell type with x-axis representing total peak count with approximate counts visualized for each opening/stable/closing groups by their relative sizes, q-value < 0.01 & Log_2_FC ≥ 0.58. **(B)** Venn diagrams show exclusive and inclusive differential peak counts for opening and closing peaks as assessed by Limma-Voom, q<0.01 and the absolute value of Log_2_FC ≥ 1.5. Individual time points and doses of radiation are plotted.

**Supplemental Figure S17.**
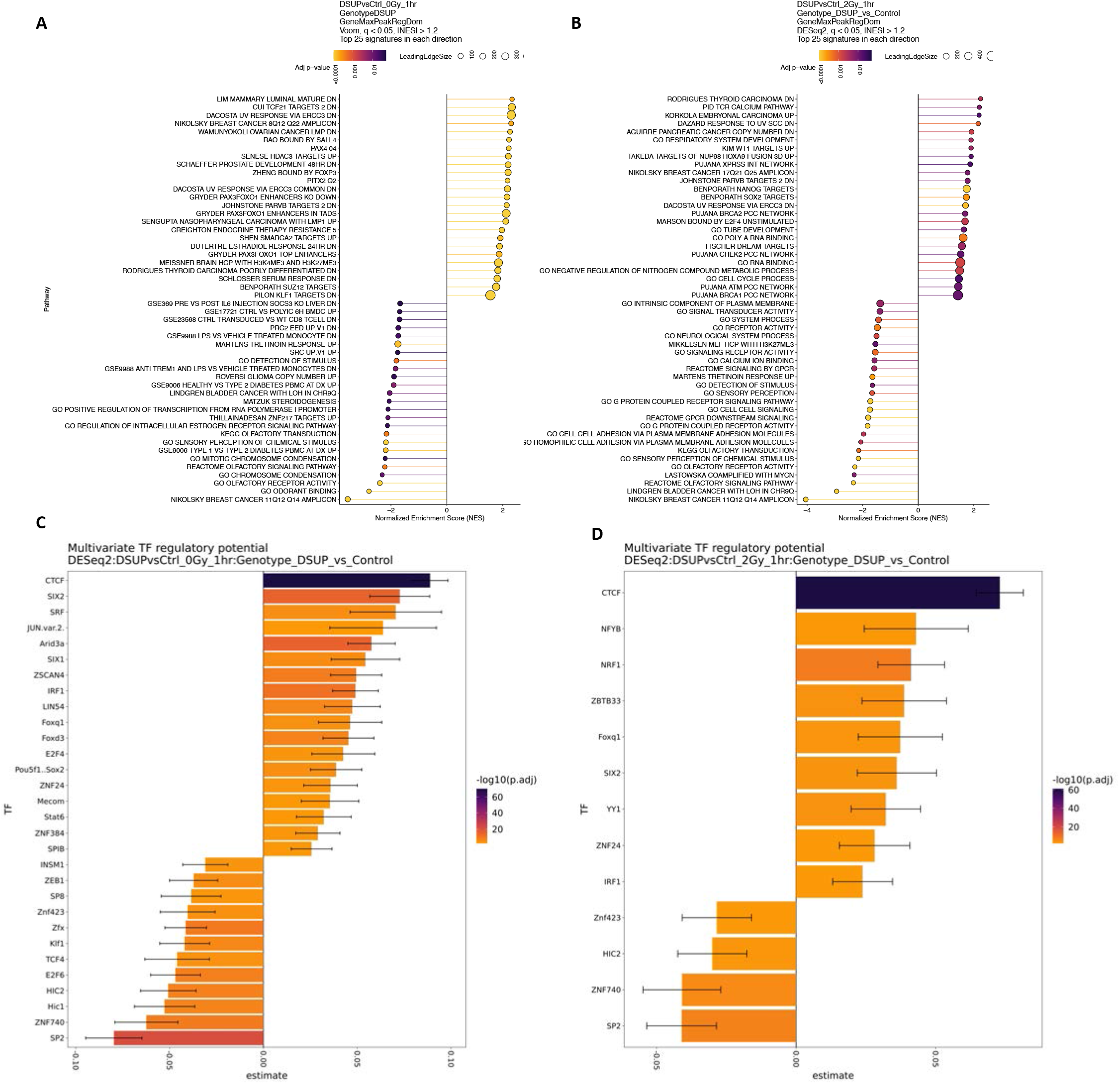
**(A)** Lollipop plot shows top differential peaks for Dsup vs Control at 0Gy and 1hr as determined by DESeq2 and then peaks are annotated to nearest gene regulatory domain using GREAT promoters and enhancers in order to obtain Fgsea associated pathways using q-value < 0.05 and absolute value of NES > 1.2. Gradient from yellow to purple shows p-adjusted values from smallest to largest with dot size representing number of leading-edge genes belonging to a particular pathway. **(B)** Lollipop plot for Dsup vs Control at 2Gy and 1hr. **(C)** Bar plot shows results of multivariate model of transcription factor regulatory potential on identified TF footprint motifs from ATACseq data for Dsup vs Control at 0Gy and 1hr. Gradient from purple to yellow represents –log10 p-adjusted value with purple being the most significant. The binding score estimate of enriched TF motifs is plotted along the x-axis as calculated by motif score and factor binding variance. **(D)** Bar Plot shows results of multivariate model for Dsup vs Control at 2Gy and 1hr.

**Supplemental Figure S18.**
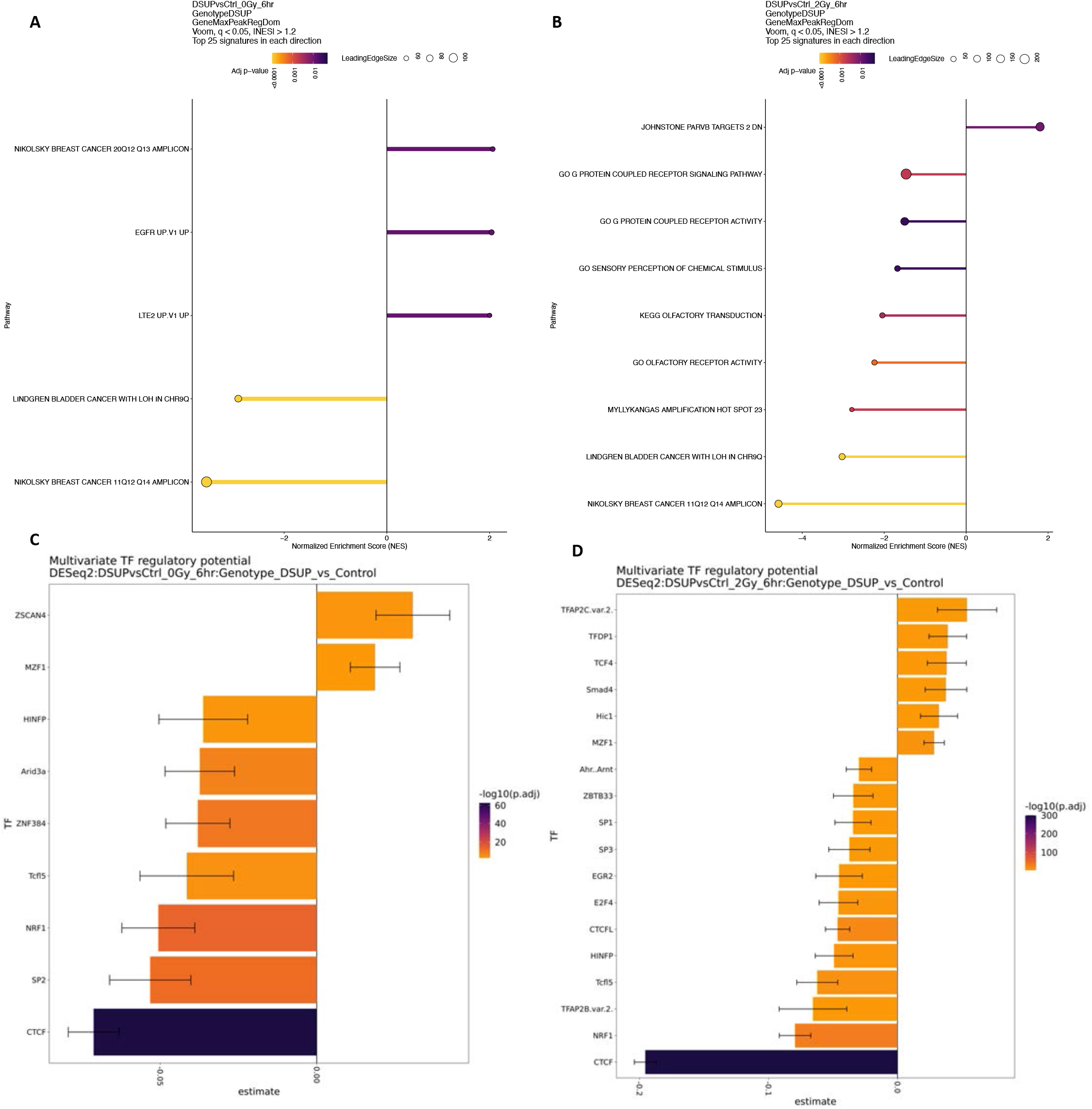
**(A)** Lollipop plot shows top differential peaks for Dsup vs Control at 0Gy and 6hr as determined by DESeq2 and then peaks are annotated to nearest gene regulatory domain using GREAT promoters and enhancers in order to obtain Fgsea associated pathways using q-value < 0.05 and absolute value of NES > 1.2. Gradient from yellow to purple shows p-adjusted values from smallest to largest with dot size representing number of leading edge genes belonging to a particular pathway. **(B)** Lollipop plot for Dsup vs Control at 2Gy and 6hr. **(C)** Bar plot shows results of multivariate model of transcription factor regulatory potential on identified TF footprint motifs from ATACseq data for Dsup vs Control at 0Gy and 6hr. Gradient from purple to yellow represents –log10 p-adjusted value with purple being the most significant. The binding score estimate of enriched TF motifs is plotted along the x-axis as calculated by motif score and factor binding variance. **(D)** Bar Plot shows results of multivariate model for Dsup vs Control at 2Gy and 6hr.

**Supplemental Figure S19.**
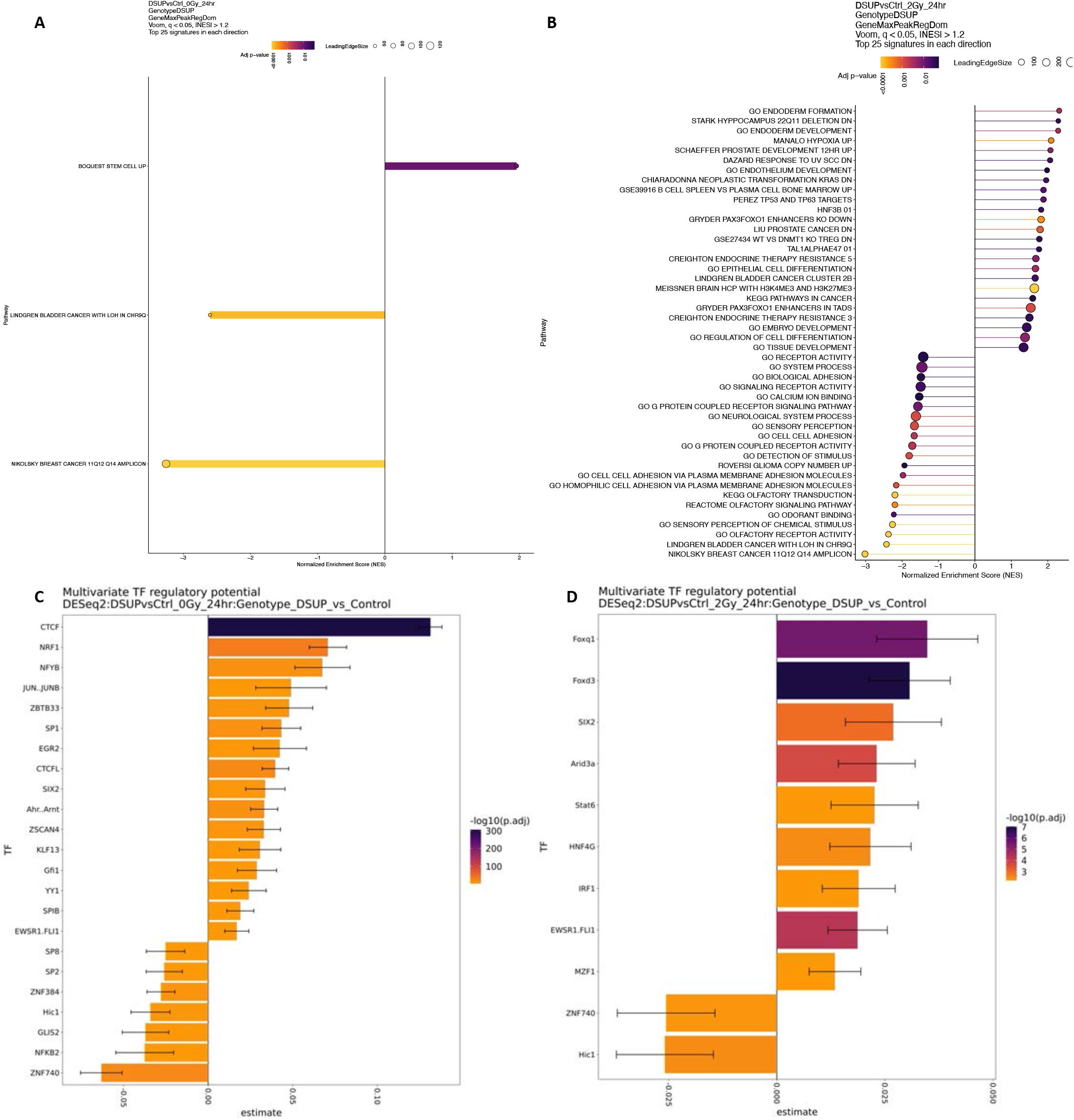
**(A)** Lollipop plot shows top differential peaks for Dsup vs Control at 0Gy and 24hr as determined by DESeq2 and then peaks are annotated to nearest gene regulatory domain using GREAT promoters and enhancers in order to obtain Fgsea associated pathways using q-value < 0.05 and absolute value of NES > 1.2. Gradient from yellow to purple shows p-adjusted values from smallest to largest with dot size representing number of leading edge genes belonging to a particular pathway. **(B)** Lollipop plot for Dsup vs Control at 2Gy and 24hr. **(C)** Bar plot shows results of multivariate model of transcription factor regulatory potential on identified TF footprint motifs from ATACseq data for Dsup vs Control at 0Gy and 24hr. Gradient from purple to yellow represents –log10 p-adjusted value with purple being the most significant. The binding score estimate of enriched TF motifs is plotted along the x-axis as calculated by motif score and factor binding variance. **(D)** Bar Plot shows results of multivariate model for Dsup vs Control at 2Gy and 24hr.

**Supplemental Figure S20.**
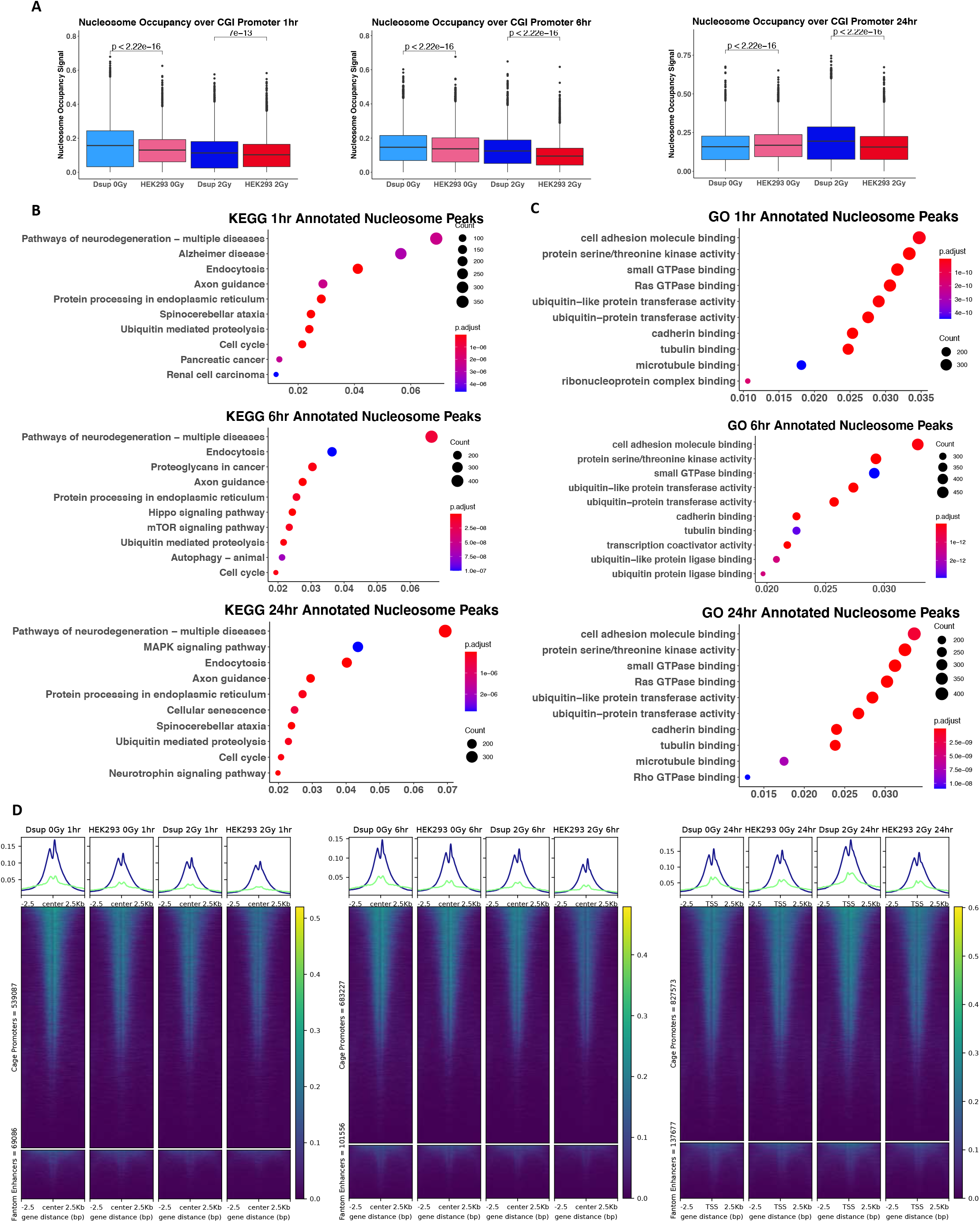
Greater nucleosome occupancy at CpG Island promoters in Dsup expressing cells. **(A)** Boxplots show comparisons of HEK293 Dsup vs HEK293 WT nucleosome occupancy signal plotted over hg38 CpG Island promoters for each time point and dose of radiation. All Statistical analysis was performed using Wilcoxon signed rank test with significant values plotted over bars with comparison to HEK293 as the reference group for each dose of radiation. **(B) (C)** Dot plots show enriched KEGG and GO pathways for annotated peaks using UCSC hg38 known genes within a ±3000bp range from the nearest TSS, p-adjusted value cutoff <0.01 with Bonferroni correction. Dot size represents number of peak counts with x-axis representing the ratio of all annotated peaks within the entire peak set. Gradient from red to blue represents the p-adjusted values with red being the highest p-adjusted value. **(D)** Heatmaps show HEK293 Dsup and HEK293 WT nucleosome occupancy signals plotted over ±2.5kb over the center of hg38 active CAGE promoters and Fantom enhancers.

**Supplemental Figure S21.**
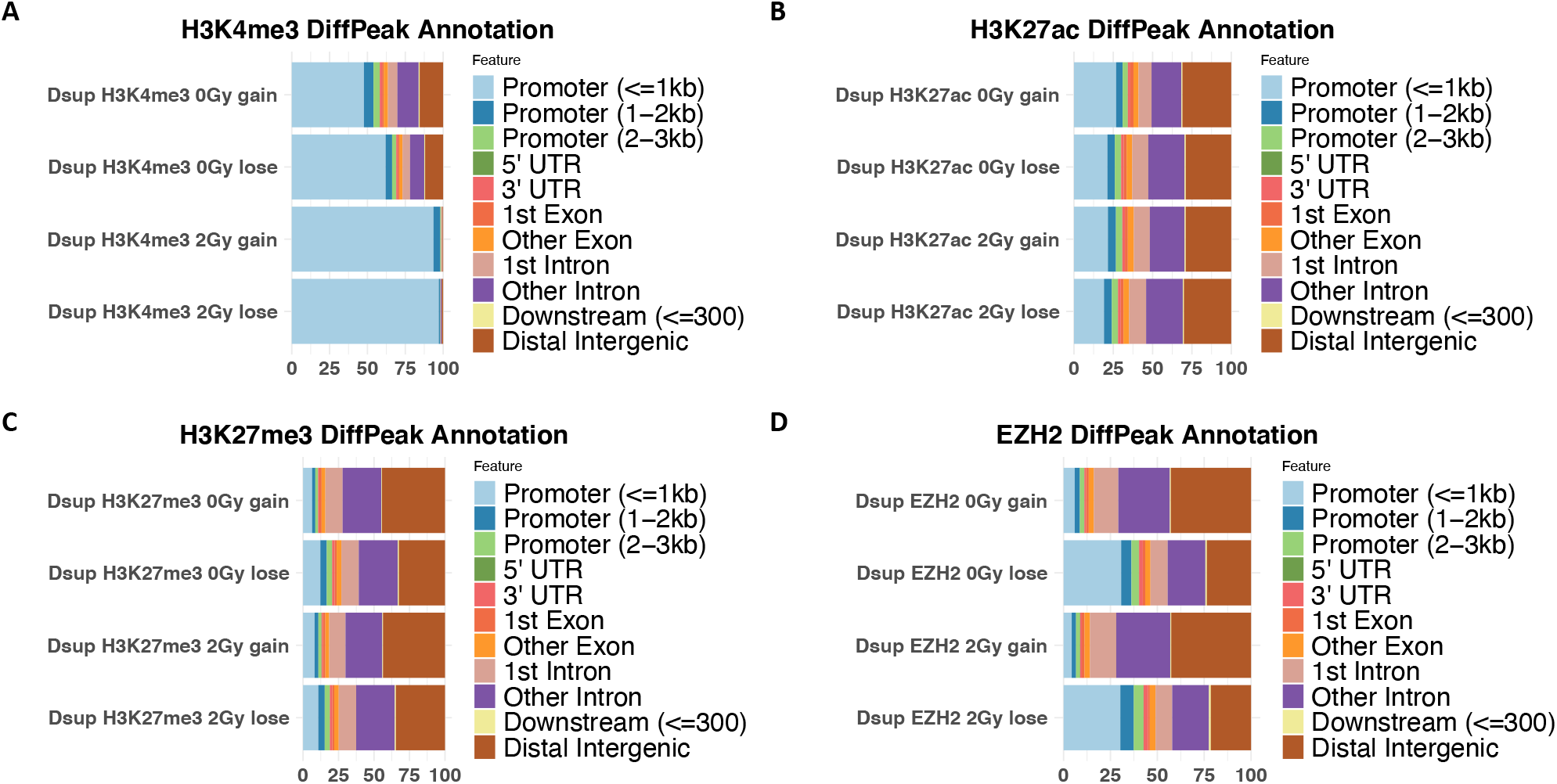
**(A)** Differential Peak analysis was performed using RGT-THOR with absolute value of Log_2_FC ≥ 1 and p-adjusted value cutoff p<0.05 for comparison of H3K4me3 peaks in HEK293 Dsup to HEK293 WT cells. Bar plot shows percent of annotated feature for H3K4me3 peaks for gains and loses in peaks at each dose of radiation. Peaks were annotated ±3000bp of TSS regions for annotated hg38 promoters. **(B)** Bar plot shows percent of annotated feature for H3K27ac peaks for gains and loses in peaks at each dose of radiation after differential peak analysis. **(C)** Bar plot shows percent of annotated feature for H3K27me3 peaks for gains and loses in peaks at each dose of radiation after differential peak analysis. **(D)** Bar plot shows percent of annotated feature for EZH2 peaks for gains and loses in peaks at each dose of radiation after differential peak analysis.

**Supplemental Figure S22.**
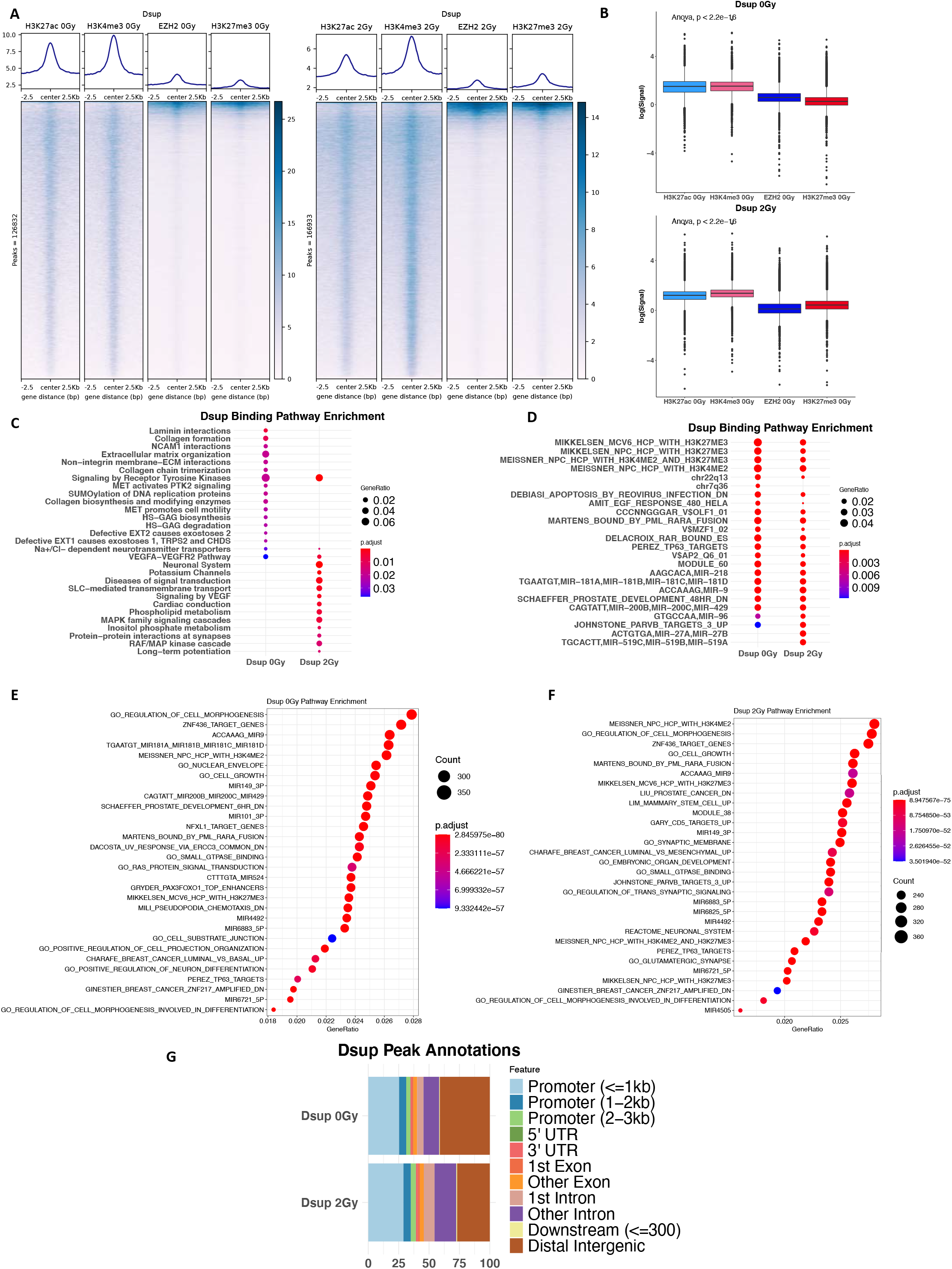
**(A)** Heatmap shows Dsup histone PTM and EZH2 signal plotted over the ±2.5kb of the center of all Dsup peaks called by MACS2 for 0Gy (left) and 2Gy (right). Active histone PTMs overlap more with Dsup peaks rather than repressive marks. **(B)** Boxplot representations of heatmap plots where log(signal) is plotted on the y-axis. Anova analysis was conducted to test the null hypothesis that the log(signal) was the same for all histone PTM and EZH2 signal. **(C)** Dot plot shows gene ratio displayed as dot size and p-adjusted values displayed as color gradient from red to blue with most significant value as blue. Here Dsup peaks are annotated ±3kb of UCSC hg38 known genes and enricher analysis with Reactome terms are displayed for each dose of radiation, p-adjusted value <0.05 with Bonferroni correction. **(D)** Dot plot shows Dsup 0Gy and Dsup 2Gy clusters compared and annotated with MsigDB v. 7.1 pathways, p-adjusted value <0.05 with Bonferroni correction. **(E)** Dot plot shows enriched MsigDB v. 7.1 pathways for just Dsup 0Gy with dot size representing associated gene count while gene ratio is plotted on the x-axis. **(F)** Dot plot shows enriched MsigDB v. 7.1 pathways for just Dsup 2Gy with dot size representing associated gene count while gene ratio is plotted on the x-axis. **(G)** Bar plots show percent of annotated feature for Dsup peaks for 0Gy and 2Gy.

**Supplemental Figure S23.**
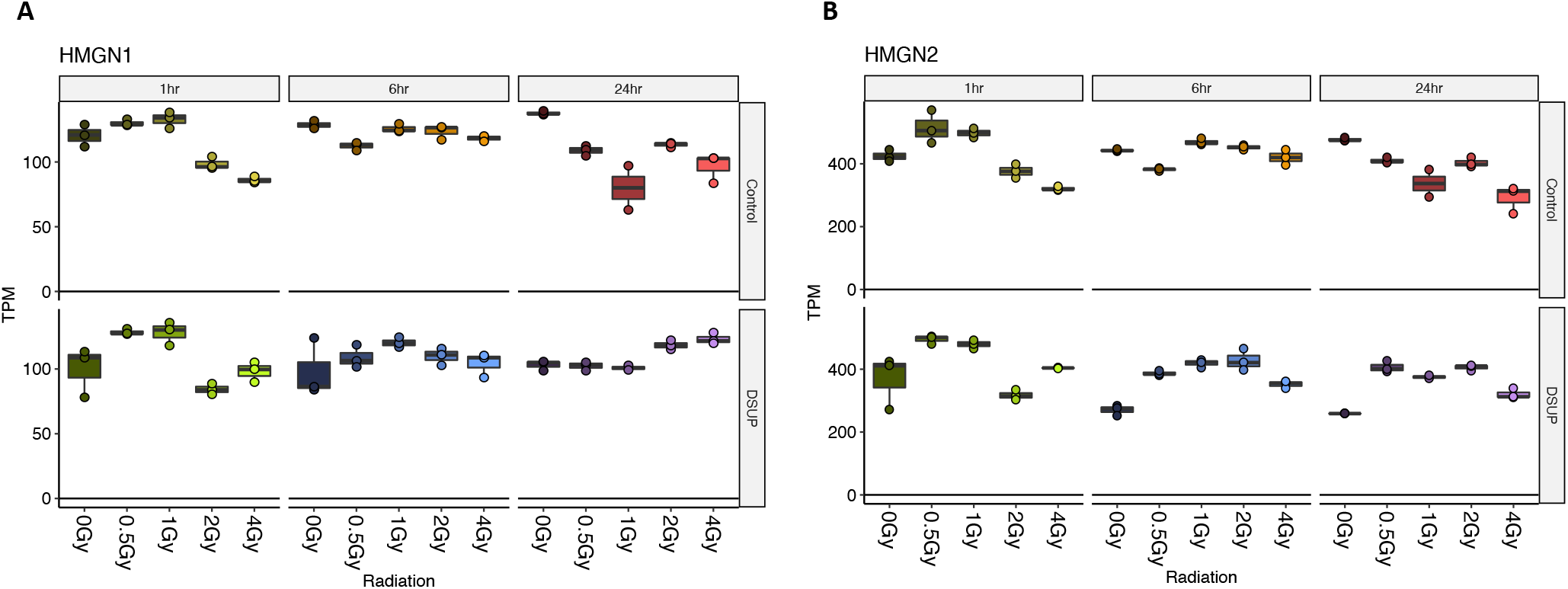
**(A)** Gene expression box plot shows transcripts per million plotted on y-axis for HMGN1 in HEK293 WT Control and HEK293 Dsup cells for each dose of radiation and timepoint. **(B)** The same RNAseq dataset but with gene expression box plot for HMGN2 across each dose and timepoint. Overall HEK293 Dsup and HEK293 WT cells have similar levels of HMGN1 and HMGN2 expression.

**Supplemental Figure S24.**
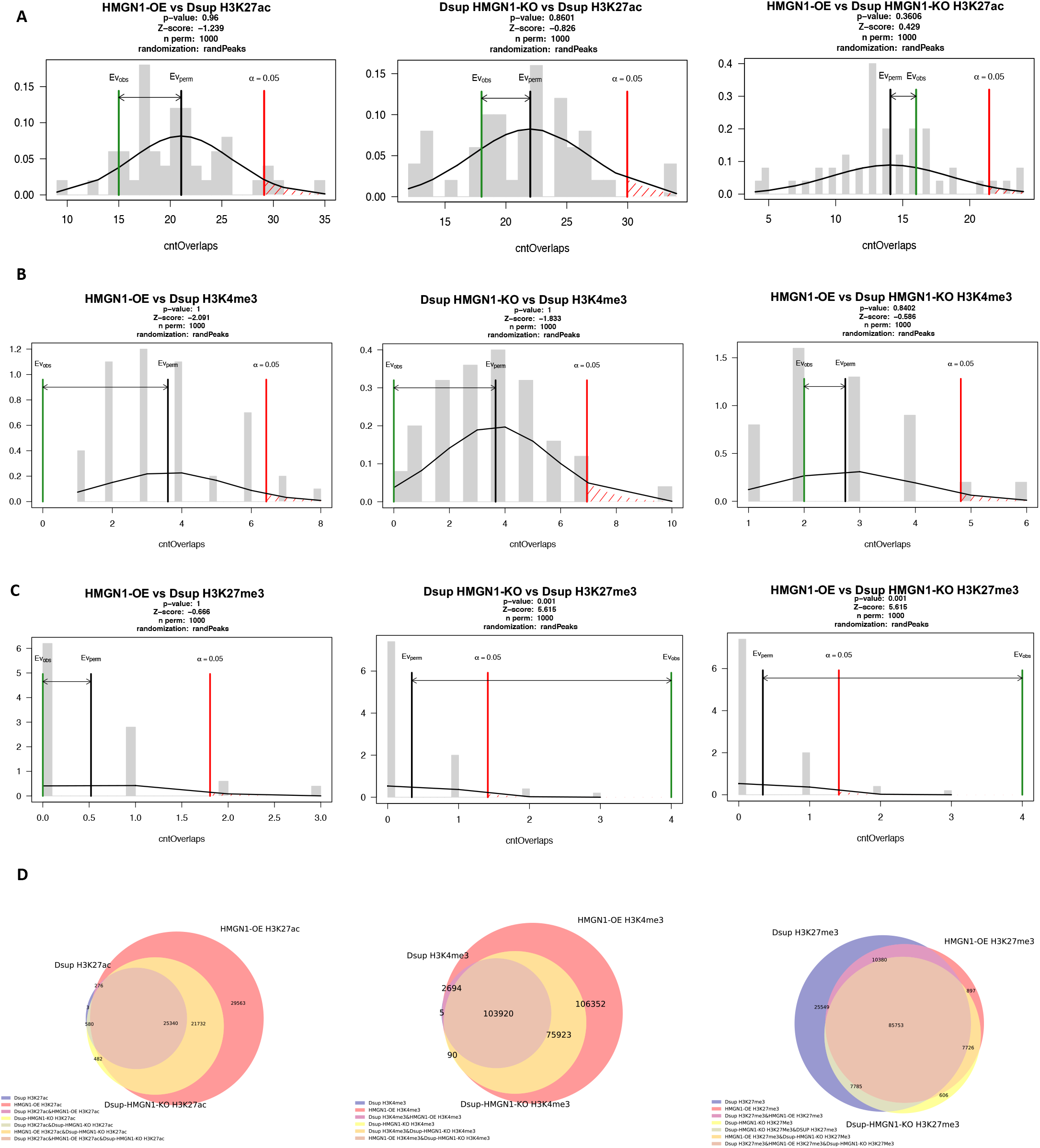
**(A)** Histograms show permutation tests for testing the null hypothesis ⍰ = 0.05 that HEK293 HMGN1-OE, HEK293 Dsup HMGN1-KO, and HEK293 Dsup have the same H3K27ac peaks after differential peaks were called using RGT-THOR. Peak overlaps were generated and and a random peak list was generated by shuffling each peak list 1000 times before permutation test was conducted using a HEK293 H3K27ac background reference for random peak sampling. X-axis shows count overlaps and y-axis shows frequency. **(B)** Permutation test results for H3K4me3 for comparison of the three cell types. **(C)** Permutation test results for H3K27me3 for comparison of the three cell types. **(D)** Venn diagrams were generated to display overlap of differential peaks among the three cell types for H3K27ac, H3K4me3, and H3K27me3.

**Supplemental Figure S25.**
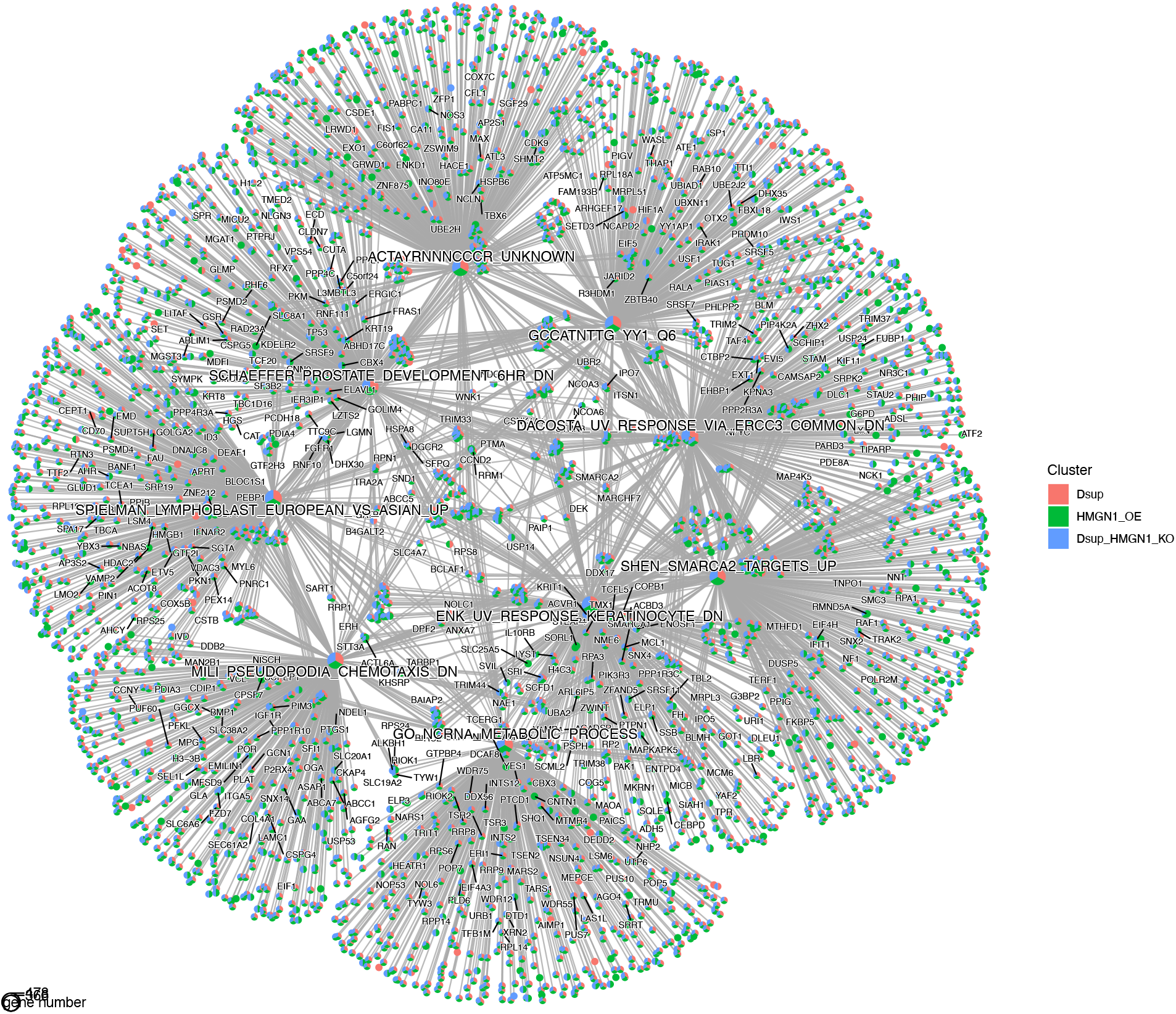
Gene concept network plot shows annotated H3K27ac peaks gained in differential peak analysis assessed by RGT-THOR. Peaks were annotated to nearest gene using UCSC hg38 known genes within a ±3000bp range from the nearest TSS, p-adjusted value cutoff <0.01 with Bonferroni correction and pathway enrichments were obtained using MsigDB terms. Pie charts clusters show breakdown of HEK293 Dsup, HEK293 HMGN1-OE, and HEK293 Dsup HMGN1-KO and are displayed over each associated pathway and genes belonging to that pathway with segments connecting related nodes.

**Supplemental Figure S26.**
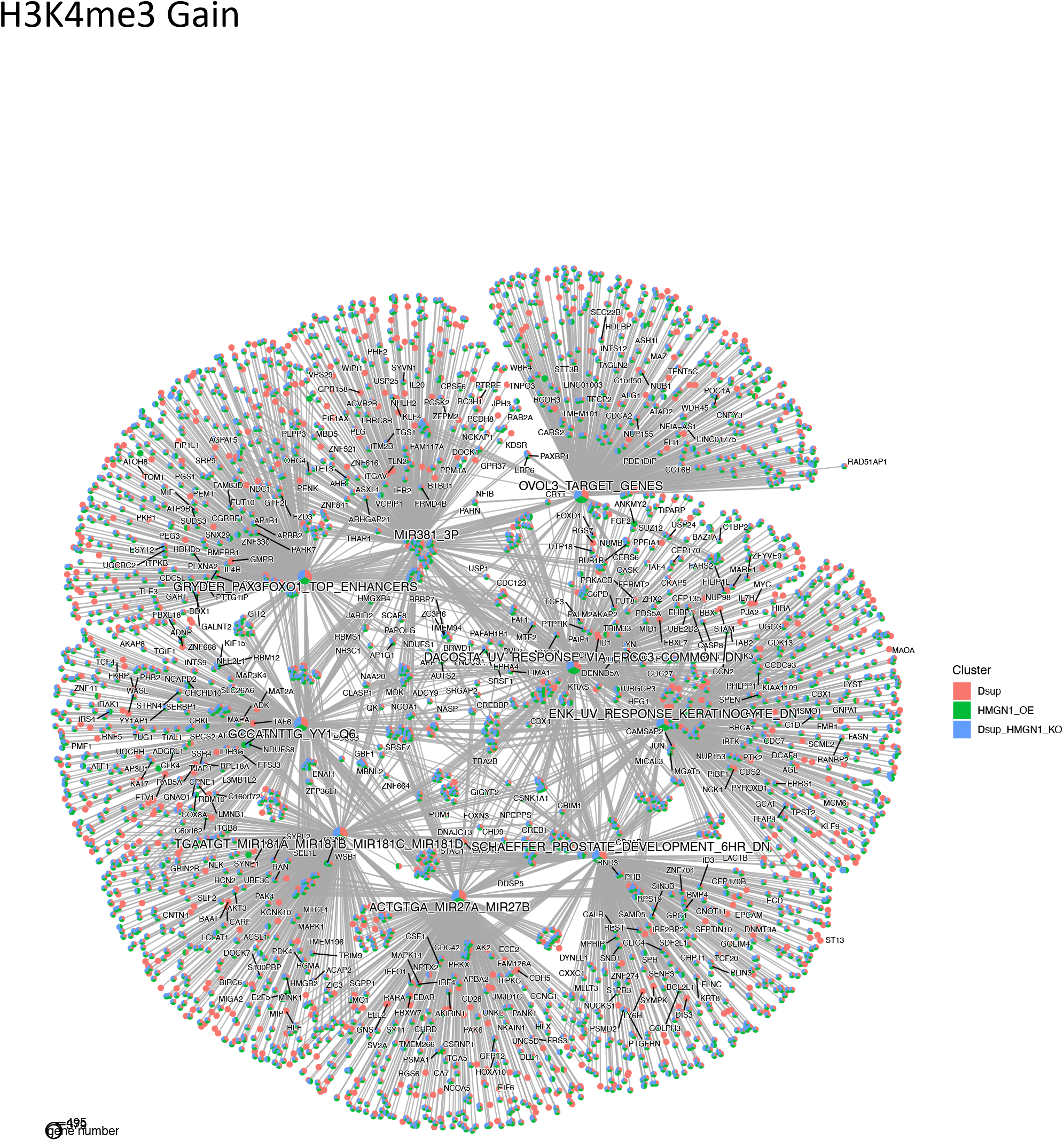
Gene concept network plot shows annotated H3K4me3 peaks gained in differential peak analysis assessed by RGT-THOR. Peaks were annotated to nearest gene using UCSC hg38 known genes within a ±3000bp range from the nearest TSS, p-adjusted value cutoff <0.01 with Bonferroni correction and pathway enrichments were obtained using MsigDB terms. Pie charts clusters show breakdown of HEK293 Dsup, HEK293 HMGN1-OE, and HEK293 Dsup HMGN1-KO and are displayed over each associated pathway and genes belonging to that pathway with segments connecting related nodes.

**Supplemental Figure S27.**
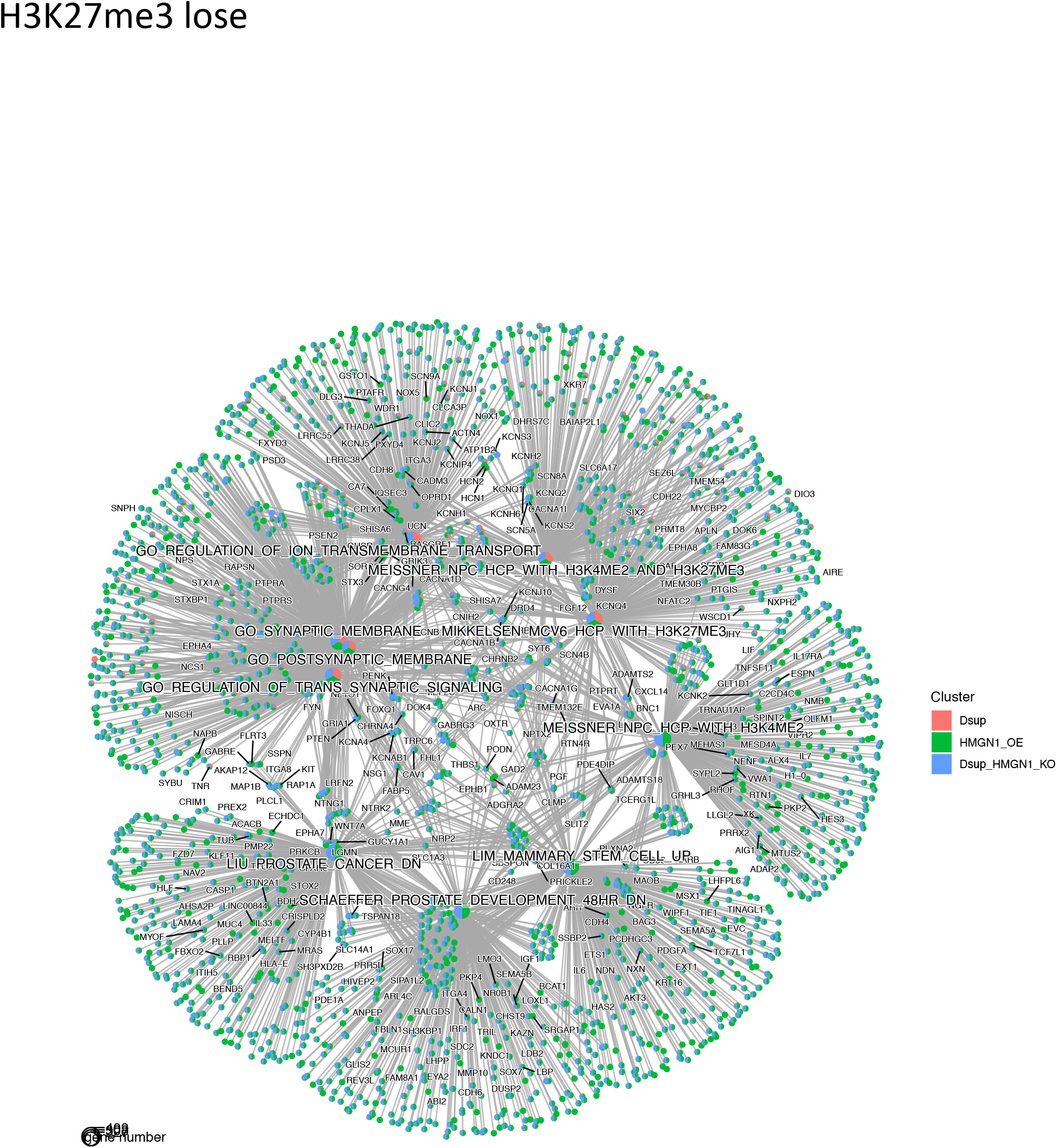
Gene concept network plot shows annotated H3K27me3 peaks lost in differential peak analysis assessed by RGT-THOR. Peaks were annotated to nearest gene using UCSC hg38 known genes within a ±3000bp range from the nearest TSS, p-adjusted value cutoff <0.01 with Bonferroni correction and pathway enrichments were obtained using MsigDB terms. Pie charts clusters show breakdown of HEK293 Dsup, HEK293 HMGN1-OE, and HEK293 Dsup HMGN1-KO and are displayed over each associated pathway and genes belonging to that pathway with segments connecting related nodes.

**Supplemental Figure S28.**
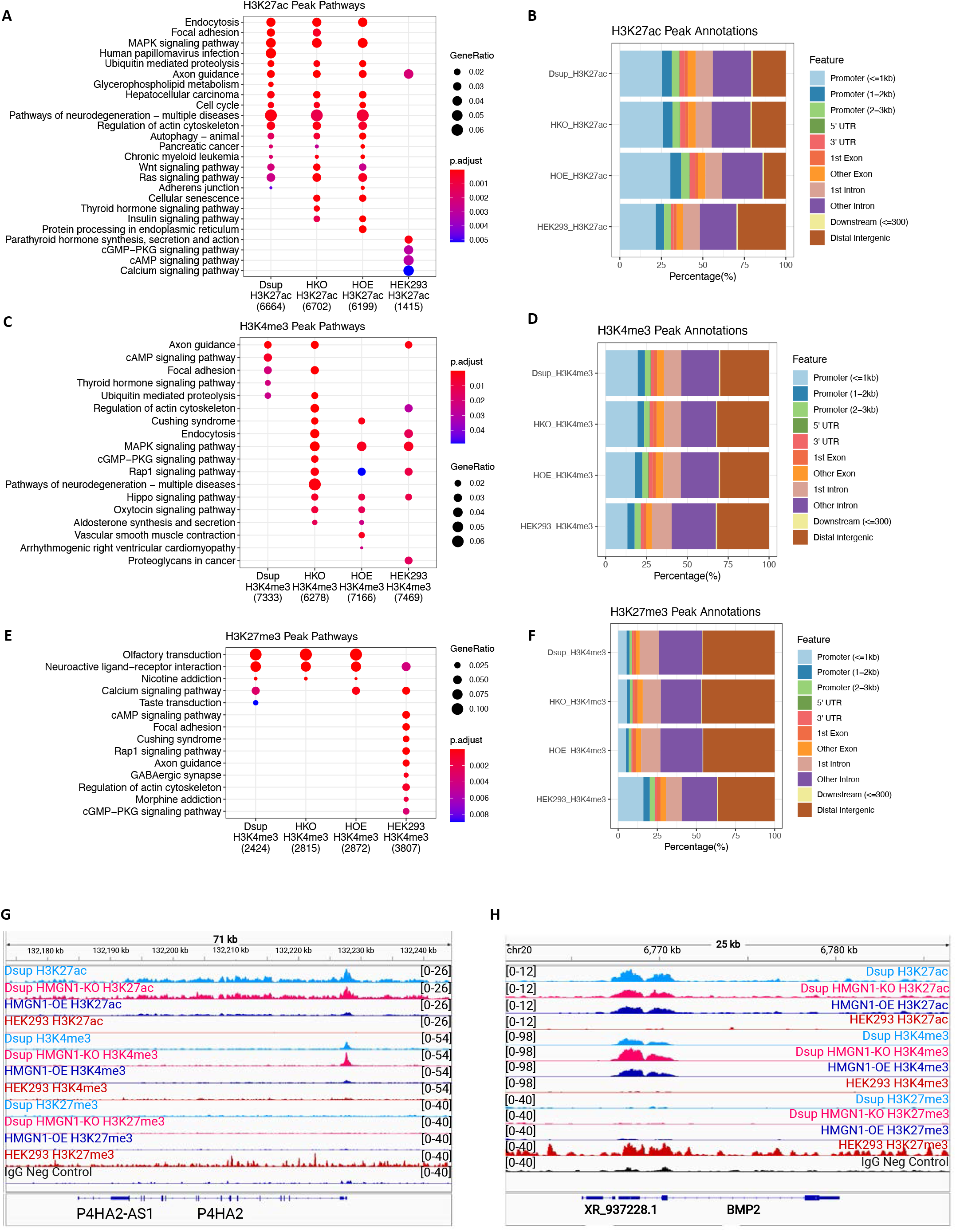
Annotation of histone PTMs for Dsup expressing cells and HEK293 HMGN1 Over Expression reveals unique pathway enrichments. **(A) (C) (E)** Dot plots show KEGG annotations for histone PTM MACS2 peak calls using narrow parameters for H3K27ac and H3K4me3 and broad peak calling for H3K27me3 with q-value <0.05 for narrow peaks and q-value<0.01 for broad peaks. Peaks were annotated to nearest gene with ±3000bp of TSS to UCSC hg38 known genes. Gene clusters comparisons were generated with a p-adjusted value cutoff of <0.01 and Bonferroni correction. Total number of peaks are shown below cell type name on the x-axis. Dot size represents the ratio of genes for a particular pathway belonging to the whole gene set with p.adjusted values represented as a gradient from red to blue with red being the highest p-adjusted value. **(B) (D) (F)** Bar plots show percent of annotated feature for MACS2 called peaks for each cell type. Peaks were annotated ±3000bp of TSS regions for annotated hg38 promoters. Dsup = HEK293 Dsup, HKO = HEK293 Dsup HMGN1-KO, HOE= HEK293 HMGN1-OE. **(G)** IGV tracks shows a representative locus on chromosome 5 for top differential Focal adhesion related Polycomb target P4HA2 for all cell types histone PTMs. Here histone PTM signal is only similar for Dsup expressing cells. **(H)** IGV tracks shows a representative locus on chromosome 20 for top differential Polycomb target BMP2 for all cell types histone PTMs. Here similar histone PTM signals can be seen for Dsup expressing cells and HEK293 HMGN1-OE cells.

**Supplemental Figure S29.**
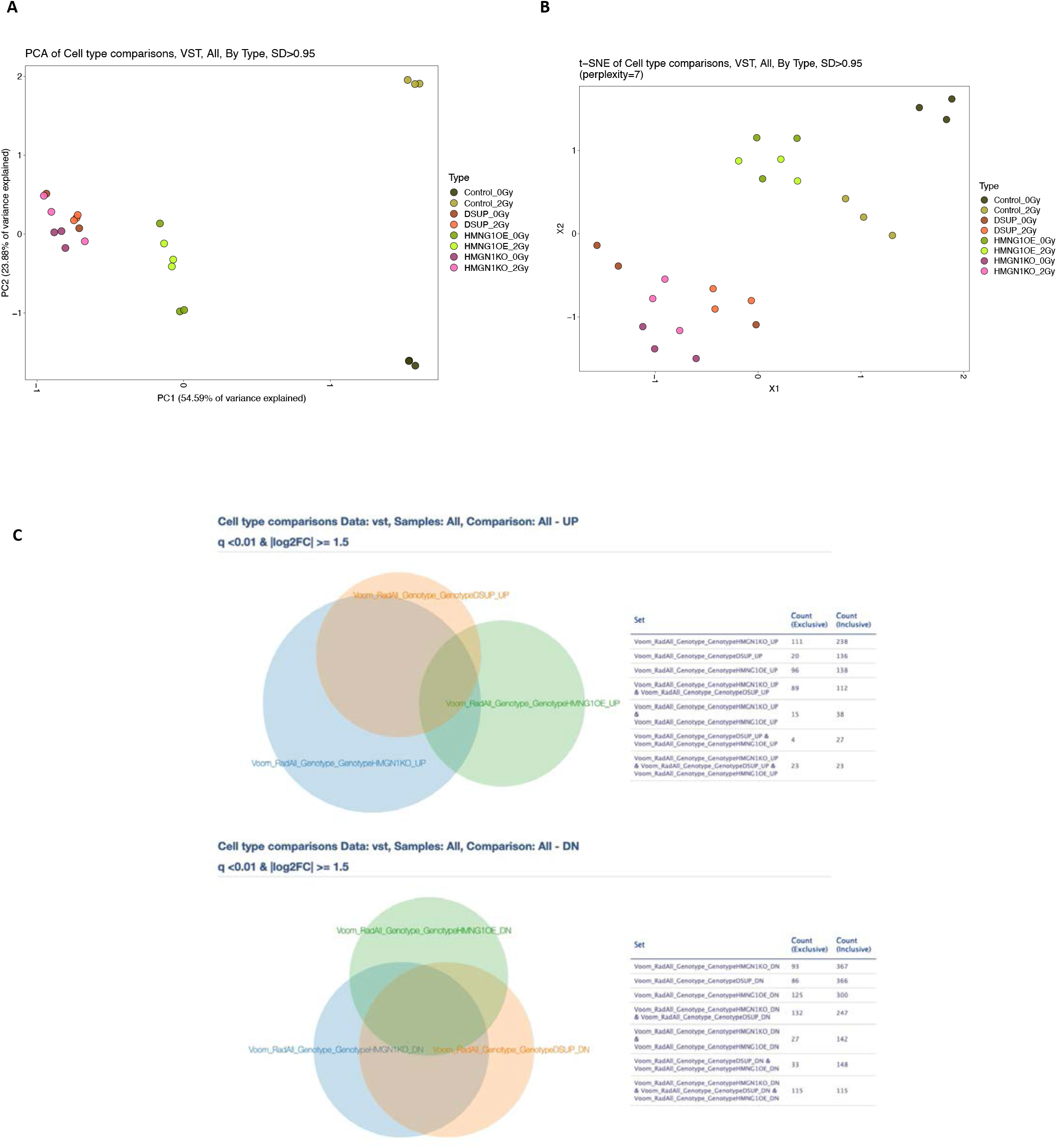
**(A)** Principal Component Analysis Clustering for expression data reveals sample to sample distance as calculated by unsupervised variance stabilizing transformation for each dose of radiation and cell type with PC1 variance on x-axis and PC2 variance on y-axis, SD>0.95. Strong clustering can be seen here among cell types with deviations between control HEK293 WT cells greatest between 0Gy and 2Gy of radiation. **(B)** T-Distributed Stochastic Neighbor Embedding shows relative distances between the same expression dataset in a 2 dimensional space, SD>0.95 and perplexity=7. Here strong clustering can be seen amongst each cell type with Control HEK293 WT cells at 2Gy of radiation more similar to HEK293 HMGN1-OE cells. Dsup expressing cells with or without HMGN1-KO clustered together as well. **(C)** Venn diagrams show exclusive and inclusive counts for differentially expressed upregulated and downregulated genes as assessed by Limma-Voom, q<0.01 and the absolute value of Log_2_FC ≥ 1.5.

**Supplemental Figure S30.**
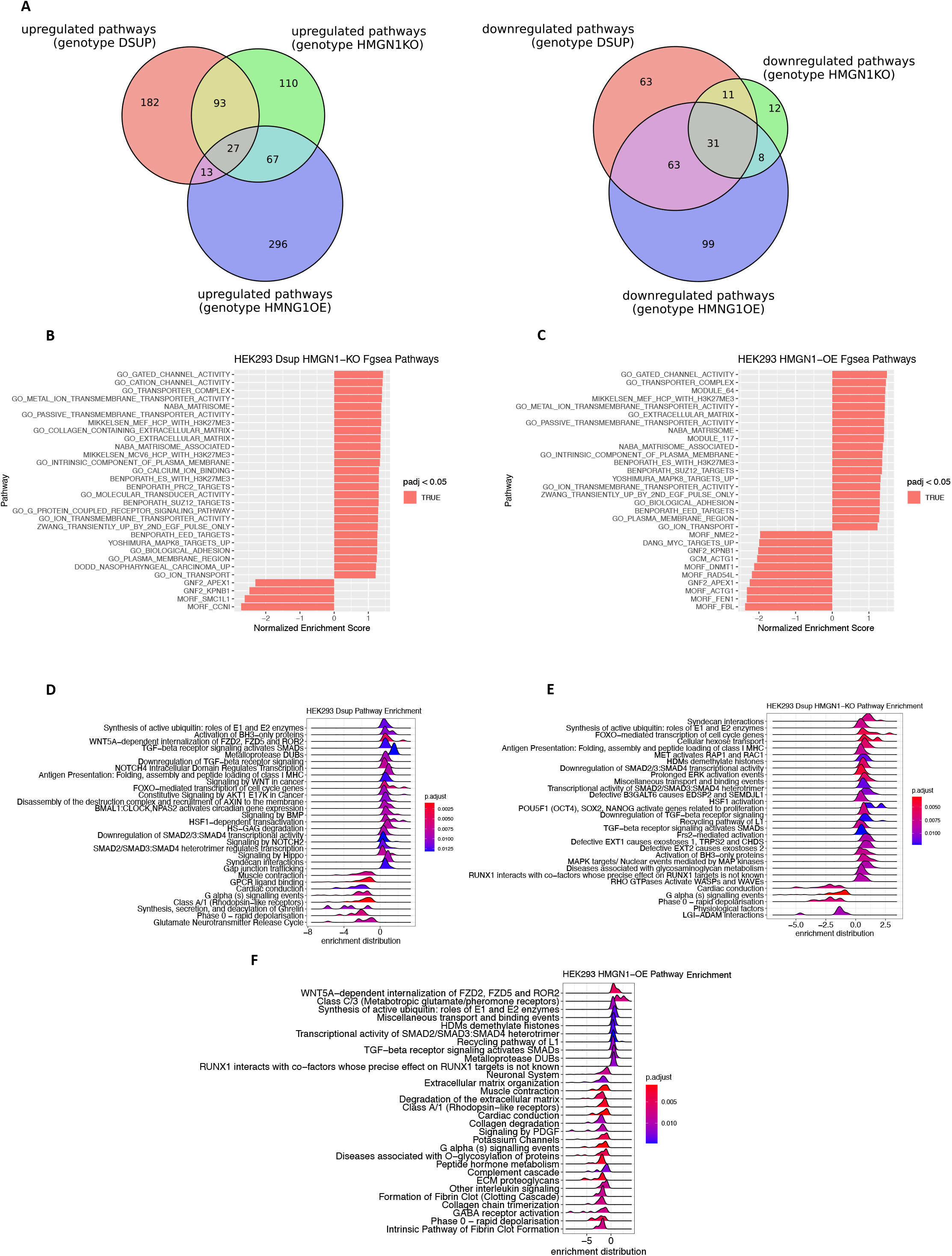
**(A)** Venn diagrams show the intersections of upregulated (left) and downregulated (right) Fgsea pathways for HEK293 Dsup, HEK293 Dsup HMGN1-KO, and HEK293 HMGN1-OE for all doses of radiation, q-value <0.01 and |NES| ≥1. **(B)** Bar plot shows top Fgsea pathways for HEK293 Dsup HMGN1-KO normalized for radiation dose. Normalized enrichment score is plotted on x-axis with significant p-adjusted value < 0.05 colored in orange. **(C)** Bar plot shows top multivariate Fgsea pathways for HEK293 HMGN1-OE normalized for radiation dose. **(D)** Ridge plot shows REACTOME enriched pathways for HEK293 Dsup normalized normalized to 0Gy and 2Gy of radiation obtained from Limma-voom differential expression analysis, p-adjusted value <0.01 and absolute value of Log_2_FC ≥ 1.5. Enrichment score distribution is plotted along the x-axis with gradient from red to blue representing p-adjusted values with blue being the most significant. **(E)** Ridge plot shows REACTOME pathway enrichments for HEK293 Dsup HMGN1-KO normalized to 0Gy and 2Gy. **(F)** Ridge plot shows REACTOME pathway enrichments for HEK293 HMGN1-OE normalized to 0Gy and 2Gy. Of note are shared upregulated pathways related to histone demethylation events and WNT signaling.

